# Long-read RNA sequencing identifies non-coding isoform switching as a regulator of cell fate

**DOI:** 10.1101/2025.02.21.639524

**Authors:** Victoire Fort, Gabriel Khelifi, Ruicen He, Valérie Watters, Melanie Pye, Nobuko Yamanaka, Marina Gertsenstein, Émeline I. J. Lelong, Jérémy Loehr, Victoria Micha, Jean-Philippe Lambert, Yojiro Yamanaka, Jeffrey Wrana, Samer M.I. Hussein

## Abstract

Development of long-read RNA sequencing technologies has paved the way to the exploration of RNA isoform diversity and its relevance in regulating cell fate. However, identifying new functional isoforms is still very difficult. Here, we leverage long-read RNA sequencing to study changes in isoforms during somatic cell reprogramming and identify novel isoforms occurring throughout cell state transitions. We demonstrate tight regulation of non-coding isoforms and show that isoform switching plays previously overlooked functional roles and outcomes in gene regulation and cell fate changes. We uncover a novel long non-coding RNA, *Snhg26*, that undergoes isoform switching during reprogramming to enhance the conversion of differentiated cells towards the pluripotent state. Knock-down of *Snhg26* in mouse and human pluripotency models reveals that it is important for pluripotency acquisition. Together, our study provides a resource to study full-length isoform usage during cell fate change. It also demonstrates the power of long-read sequencing to identify functionally relevant gene isoforms in the context of cell plasticity.

## Introduction

Among high-throughput technologies, the profiling of RNA transcripts, known as transcriptomics, has become a gold standard to identify the molecular markers that define cell identities and plasticity^1,2^. Despite the extensive collection of RNA sequencing (RNAseq) datasets generated by various independent research teams and international consortiums^3–6^, technical limitations have hindered our efforts to obtain a comprehensive and detailed overview of full transcriptomes. Notably, long coding and non-coding transcripts expression, repetitive sequences and transposable elements expression, as well as isoform diversity have been largely overlooked. Indeed, short-read RNAseq cannot cover the full-length sequence of a transcript and thus doesn’t have the power to unravel complex locus sequences and isoform diversity^7,8^. Long-read RNA sequencing (lrRNAseq) technologies, that are able to identify full-length transcripts^9,10^, can overcome this issue and deepen our understanding of cell biology and organism development. For instance, Leung *et al.* have uncovered the existence of novel isoforms in the context of both human and mouse cortices, and highlighted changes in alternative splicing between human foetal and adult cortices^11^. On the other hand, studies in human^12^ and mouse^13^ pre-implantation embryos have described thousands of unannotated isoforms, transcribed from both known and novel genes. In humans, Torre *et al.* have shown significant modulations of alternative splicing events associated with disruption of open reading frames during embryonic genome activation^12^. In the mouse, Qiao *et al*. have leveraged their lrRNAseq data analysis to identify new transcript isoforms where their depletion negatively impacted embryo development^13^. Surprisingly, aside from the latter study, none of these generated datasets have been used yet to identify functionally relevant new gene isoforms in a biological context.

Among the turning points of embryonic development, the establishment of pluripotent stem cells (PSCs) and their differentiation towards other cell types is crucial. Understanding the regulatory network of PSCs has enabled the seminal work by Yamanaka *et al.*, which has led to the discovery that a small set of genes (*Oct4, Klf4, c-Myc* and *Sox2*, termed “OKMS”) can reprogram cells into induced pluripotent stem cells (iPSCs)^14^. Since then, extensive studies have been conducted to understand the principles driving this process of cellular plasticity by characterizing reprogramming and PSCs in terms of epigenetics^15,16^, transcriptomics^16–18^, proteomics^19,20^, transcription factor networks^21,22^ and metabolomics^23,24^. This body of work has established that induction and maintenance of the pluripotent state are regulated by a gene regulatory network composed of hierarchical and interconnected regulatory circuits mediated by extrinsic signals, key transcription factors, chromatin modifying enzymes, microRNAs and long non-coding RNAs (lncRNAs)^25–27^. Additionally, post-transcriptional regulation (e.g. RNA splicing) has been identified as a potential player in the regulation of reprogramming towards the pluripotent cell state^28–30^. Indeed, short-read RNAseq analyses of mouse reprogramming has demonstrated that splicing patterns from the somatic cells are converted into that of pluripotent cells during iPSC induction, and that RNA-binding proteins (RBPs) regulate alternative splicing in a temporal manner^31,32^. Additionally, overexpressing or knocking-down splicing regulators enhanced or suppressed the efficiency of the reprogramming process, validating their importance in pluripotency regulation^31,33–35^. Despite the admitted role of alternative splicing and splicing regulators in reprogramming, isoform diversification that inherently comes with alternative splicing events has never been investigated in this context.

In this work, we characterize in detail stem cells as well as for the first time the process of reprogramming mouse somatic cells to iPSCs using lrRNAseq. We identify multiple new isoforms that result from previously undetected splicing patterns, and that are differentially regulated during the reprogramming process. More specifically, non-coding isoforms undergo negative regulation during the mid-phases of reprogramming, and about 6-16% of spliced genes exhibit a switch in expression of their isoforms. This included long non-coding RNAs, which suggests lncRNAs undergoing isoform switch may have important functions during the acquisition of pluripotency. We demonstrate that overexpression of a specific isoform of *Snhg26* lncRNA, *Snhg26.217*, enhances the reprogramming to the pluripotent state. We further show that *Snhg26* knock-out results in defects of an *in vitro* blastoid model, but only the *Snhg26.217* isoform rescues the observed developmental phenotype. Conversely, we show that knock-down of this lncRNA in mouse embryonic stem cells (ESCs) primes cells for differentiation, validating its function in the regulation of pluripotent stem cells. Finally, we show that knock-down of the human ortholog of *Snhg26* leads to a similar outcome in human pluripotent stem cells and also decreases the efficiency of human reprogramming. These results highlight the discovery of a new conserved lncRNA involved in the regulation of the pluripotent state. Together, our study demonstrates the benefits of lrRNAseq to identify new functionally relevant gene isoforms, in the face of exploring isoform diversity. The dataset we generated also stands as a valuable resource for other researchers to study the role of alternative splicing and isoform diversity in the regulation of cell fate.

## Results

### Defining reprogramming stages by lrRNAseq

Addressing isoform diversity in reprogramming is hindered by the low efficiency of the process^36^. Here, we designed two reprogramming systems that allow generation of sufficient amounts of reprogramming cells for high resolution analysis of isoform diversity. Because overactivation of *c-Myc* induces transcriptional and splicing events caused by its effect on promoting proliferation^37–39^, we devised our systems where *c-Myc* can be either included (OKMS) or omitted (OKS) from the cocktail of reprogramming factor transgenes. This way, reprogramming-induced events that are independent of *c-Myc* can be isolated (please refer to **Supplementary Note** for more details on the system). Briefly, we generated stable cell lines of mouse neural progenitor cells (NPCs) with doxycycline (DOX)-inducible reprogramming factor transgenes driving the expression of an *mCherry* reporter gene (**Fig. 1a** and **Supplementary Fig. 1a-b**). NPCs were chosen as they reprogram more efficiently than the commonly used mouse embryonic fibroblasts (MEFs), enabling shorter experimental timelines, especially for OKS reprograming systems^40^ (please refer to **Supplementary Note** for more details on this choice). As transgene silencing is a necessary step to reach *bona fide* pluripotency at the end of reprogramming^41–43^, DOX is removed at D6, and cells able to fully reprogram convert to iPSCs (please refer to **Supplementary Note** for more details on DOX removal). To monitor the appearance of iPSCs, the NPC lines also harbor a *GFP* transgene under the control of endogenous *Oct4* gene promoter expression, which is activated when cells reach the iPSC state (**Fig. 1a** and **Supplementary Fig. 1a-b**). In parallel, we also maintain cells with DOX during the whole reprogramming process to reproduce other previously shown reprogramming paths^16^. For these cells, transgene silencing naturally occurs in late timepoints^44^, causing a decrease in mCherry-positive cells (**Supplementary Fig. 1a-b**). Therefore, we collected samples with DOX at D2, D4, D6, D10, D14, D18, and D20. We also collected iPSC lines after DOX removal at either D14, D18, or D20 (**Fig. 1a**).

**Figure 1:**
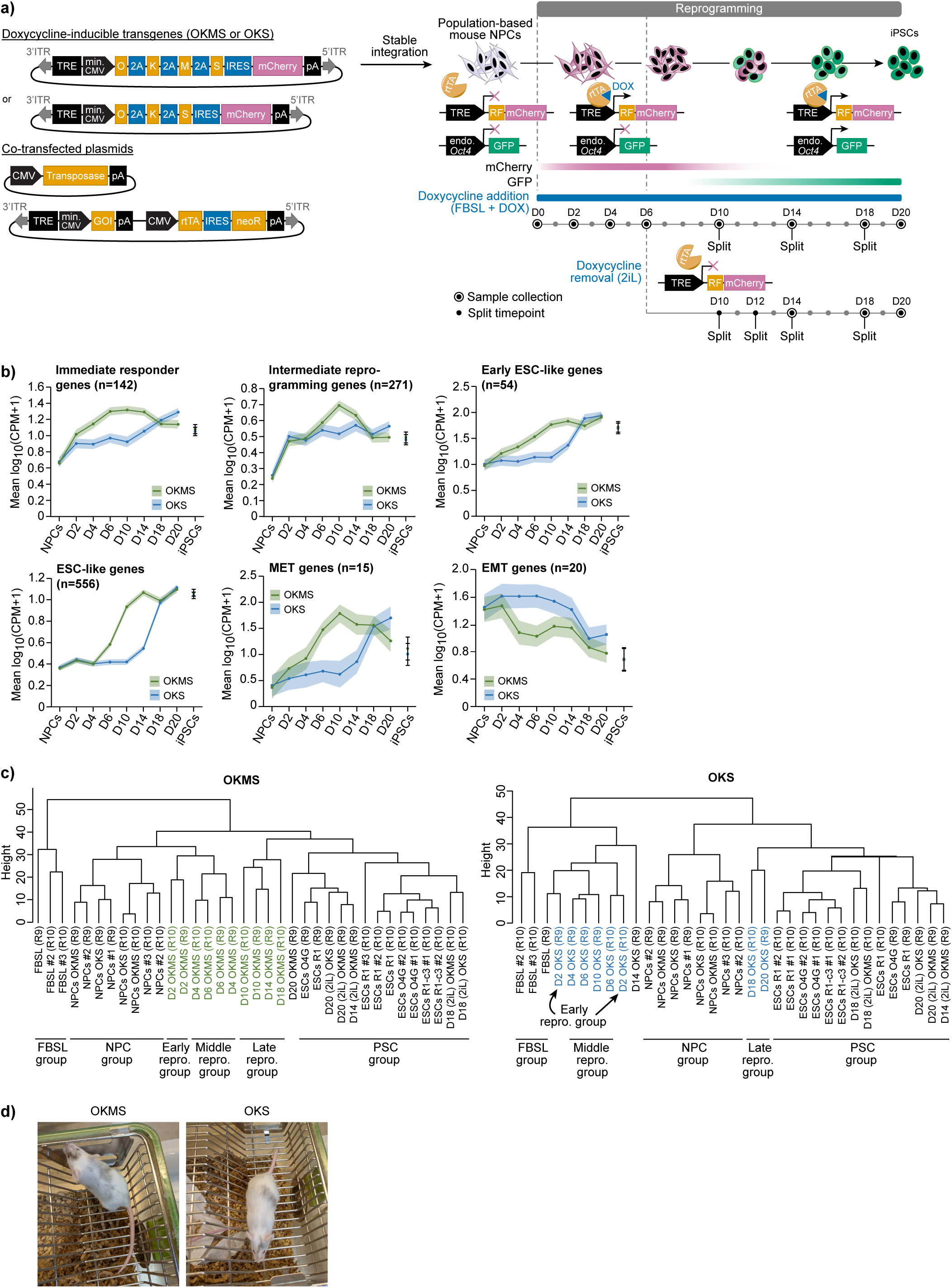
OKMS and OKS reprogramming systems that recapitulate expected gene expression. **a)** Schematic of the experimental reprogramming pipeline: NPCs harboring a *GFP* reporter under the control of the endogenous *Oct4* gene promoter were transfected with OKMS or OKS reprogramming factors driving the expression of an *mCherry* reporter. Co-transfection with a transposase, rtTA and a neomycin resistance gene allows for selection of cells with stably integrated cassettes through *piggyBac* transposition. Addition of doxycycline (DOX) to the medium induces overexpression of OKMS or OKS transgenes and leads to mCherry+ cells. Doxycycline is added either for the whole reprogramming process (upper track) or for six days only (lower track), leading to silencing of the transgenes and to GFP+ iPSCs. Samples were collected at different timepoints and processed for long-read RNA sequencing. ITR: Inverted Terminal Repeat (i.e. *piggyBac* transposition sequence); TRE: Tet Response Element; min. CMV: minimal CMV promoter; IRES: Internal Ribosome Entry Site; pA: poly(A) signal sequence; GOI: Gene Of Interest; rtTA: reverse tetracycline-controlled TransActivator; neoR: neomycin Resistance gene; NPCs: Neural Progenitor Cells; iPSCs: induced Pluripotent Stem Cells; RF: Reprogramming Factors (i.e. *Oct4* (O), *Klf4* (K), *c-Myc* (M) and *Sox2* (S)); endo.: endogenous; FBSL: FBS-LIF medium; 2iL: 2 inhibitors + LIF medium. **b)** Plots of logged expression for immediate responder, intermediate reprogramming, early ESC-like and ESC-like genes (gene lists from Hussein *et al*.^16^), or mesenchymal-to-epithelial transition (MET) and epithelial-to-mesenchymal transition (EMT) genes in OKMS or OKS reprogramming systems. Dots and lines represent average expression per timepoint, and shaded areas and error bars represent the standard error of the mean (SEM). CPM: Counts Per Million reads; ESC: Embryonic Stem Cell. **c)** Hierarchical clustering on top varying genes of the OKMS or OKS samples. The data is the same in OKMS and OKS for the NPC, FBSL and PSC group samples. repro.: reprogramming; PSC: Pluripotent Stem Cell. **d)** Images of chimeric weaned mice obtained from aggregation of OKMS or OKS iPS clones with 8-cell CD-1 albino embryos for pluripotency assessment.

To validate our reprogramming systems, we performed whole transcriptome analysis using lrRNAseq at different timepoints (n=45 total samples) of the process (**Fig. 1a** and **Supplementary Data 1**). Analyzing expression of reprogramming and PSC marker gene sets (previously described^16^), OKMS reprogramming rapidly activated immediate responder genes, intermediate reprogramming genes, genes associated with mesenchymal-to-epithelial transition (MET) (**Fig. 1b** and **Supplementary Data 1**), all of which are signs of proper of reprogramming^16^. Early and late ESC-like genes were activated as early as D4 and D6, respectively (**Fig. 1b**). This was accompanied by decreased expression of epithelial-to-mesenchymal transition (EMT) genes starting at D4 (**Fig. 1b**) suggesting our system recapitulates MET, a hallmark process observed during reprogramming to iPSCs^45,46^. OKS reprogramming showed slower dynamics, and the later expression of the gene sets compared to OKMS reflects the well-known delay in OKS kinetics and is due to lack of *c-Myc*, which is known to enhance reprogramming by increasing proliferation^47,48^. Nonetheless, OKS reprogramming showed rapid increase in expression of immediate responder and intermediate genes, and to some extent MET genes within the first 4 days (**Fig. 1b**) when compared to NPCs. These results suggest that both our OKMS and OKS systems constitute efficient reprogramming strategies, which faithfully recapitulate the global gene expression changes observed in other studies.

Because of the differences in OKMS and OKS kinetics, we undertook hierarchical clustering analysis in order to group OKMS and OKS timepoints that are similar based on expression of the top 10% varying genes (**Fig. 1c**). This segregated the samples into 5 different categories: First the “NPC group” that gathers four replicates of wild-type NPCs, the OKMS-NPCs and the OKS-NPCs at D0. OKMS and OKS reprogramming cells at D2 were classified as an “early reprogramming group”, while D4 to D6 for OKMS or D4 to D10 for OKS clustered together and form the groups of “middle reprogramming” phases. OKMS cells between D10 and D18, and OKS cells at D18 and D20 are clustered in a separate “late reprogramming group” that resembles a “PSC group”, which is composed of ESCs, OKMS- and OKS-iPSCs in 2i-LIF media. This is consistent with the OKMS samples being quickly redirected towards the induced pluripotent state and achieving pluripotency reacquisition earlier than OKS samples (**Fig. 1b** and **Supplementary Note Fig. 1c**). Finally, a sixth group comprising samples of NPCs placed in FBS-LIF medium without reprogramming factors (“FBSL”) (**Fig. 1c**) was used to assess transcriptional and splicing changes induced simply by the change of culture media. The segregation of these groups based on reprogramming phases was used in the subsequent analyses of our dataset. Looking at global gene expression, we observed a two-wave pattern (at the beginning of reprogramming upon DOX induction, and at the late phase), highlighting again known reprogramming dynamics, as previously described^41,49^ (**Supplementary Fig. 1c** and **Supplementary Data 1**).

Finally, we validated that iPSC clones isolated from our OKMS and OKS reprogramming experiments were able to contribute to the formation of chimeric mice when aggregated with 8-cell stage mouse embryos. The clones contributed 10-50% to chimera formation, as judged by coat color (**Fig. 1d**). This confirmed that our reprogramming systems give rise to *bona fide* iPSCs. Altogether, these results suggest that we generated both *Myc*-dependent and *Myc*-independent reprogramming systems that follow the well-accepted kinetics of reprogramming and can be compared in further steps of analysis.

### Unprecedented isoform diversity and tight regulation of non-coding isoforms

Alternative splicing has been extensively characterized in pluripotent stem cells and during reprogramming^28,32,50^. However, a detailed characterization of transcriptomes from the different cell states found on the trajectory of the reprogramming cells has not yet been undertaken. In order to analyze isoform diversity and confidently identify functional isoforms in our reprogramming systems, we combined single transcriptomes built from each of the 45 samples that we sequenced. We identified 279,998 isoforms that can be reliably detected and supported by several criteria (**Supplementary Fig. 2-6a**, **Supplementary Data 2-6**). Among these criteria, we focused on isoforms that are sufficiently expressed (≥ 1 transcript per million reads (TPM) in at least 3 samples), have a poly(A) motif, and have no reverse transcriptase switching events (**Supplementary Fig. 2** and **Supplementary Fig. 3a**), where the transcription start site (TSS) was supported by orthogonal approaches (e.g. CAGEseq, 5’ end sequencing, H3K4me3 ChIPseq peaks, etc.) (**Supplementary Fig. 3b-c** and **Supplementary Fig. 4**), and where splice junctions were supported by short-read RNAseq, presence of repeat elements, or mouse ESCs proteomics data (**Supplementary Fig. 3d**-**g** and **Supplementary Data 2-3**). Lastly, to compare supported and non-supported isoforms (5.7% not supported by TSS (**Supplementary Fig. 3b**), ∼20% not supported by junctions (**Supplementary Fig. 3d**)), we evaluated several metrics of isoform quality (i.e. length, number of exons, expression and coverage at the lowest-covered splice junction in short-read datasets) and show that non-supported isoforms do not exhibit any major differences from supported isoforms (**Supplementary Fig. 5**). From all isoforms, only 1.2% was not supported neither by TSS nor by junctions and were filtered out to obtain 276,608 transcripts (**Supplementary Fig. 2** and **Supplementary Fig. 6a**). We also compared our transcriptome to ones built by three orthogonal transcriptome-building tools: Bambu^51^, IsoQuant^52^, and StringTie^53^. Strikingly, all tools showed ∼50% reproducibility when compared to one another (**Supplementary Fig. 6b**, **Supplementary Data 3**, and **Supplementary Data 6**). This is concordant with a recent benchmarking study by the Long-read RNA-Seq Genome Annotation Assessment Project^54^.

Next, we used SQANTI3 software^55^ to classify the 276,608 isoforms. From these, only 35.3% of isoforms were annotated, whether they matched completely (20.53% “Full Splice Matches” or FSM) or partially (14.81% “Incomplete Splice Matches” or ISM) to the reference transcriptome (**Fig. 2a** and **Supplementary Data 2**). Indeed, reprogramming is a process where many cells undergo rapid proliferation, a mesenchymal-to-epithelial transition, DNA damage, global chromatin marks reorganization and a change in cell type, all of which can greatly alter the transcriptome^56^. Thus, this could explain the identification of the remaining 178,870 (64.70%) novel isoforms. From the total, 60.7% of the transcripts were new isoforms from known genes, but with differing splice-junction combinations, with 15.95% coming from new combinations of known splice junctions (“Novel-In-Catalog” or NIC) and 44.74% from novel splice junctions (“Novel-Not-in-Catalog” or NNC) (**Fig. 2a**). The remaining 4% of the detected isoforms were new transcripts generated from novel genes (i.e. 11,004 isoforms), with almost two third of these originating from antisense (1.32%) and genic (1.17%) isoforms, whereas intergenic (0.99%) and fusion (0.50%) transcripts accounted for the rest (**Fig. 2a**). These results are consistent with previous transcriptome analyses using lrRNAseq^11,13,55^.

**Figure 2:**
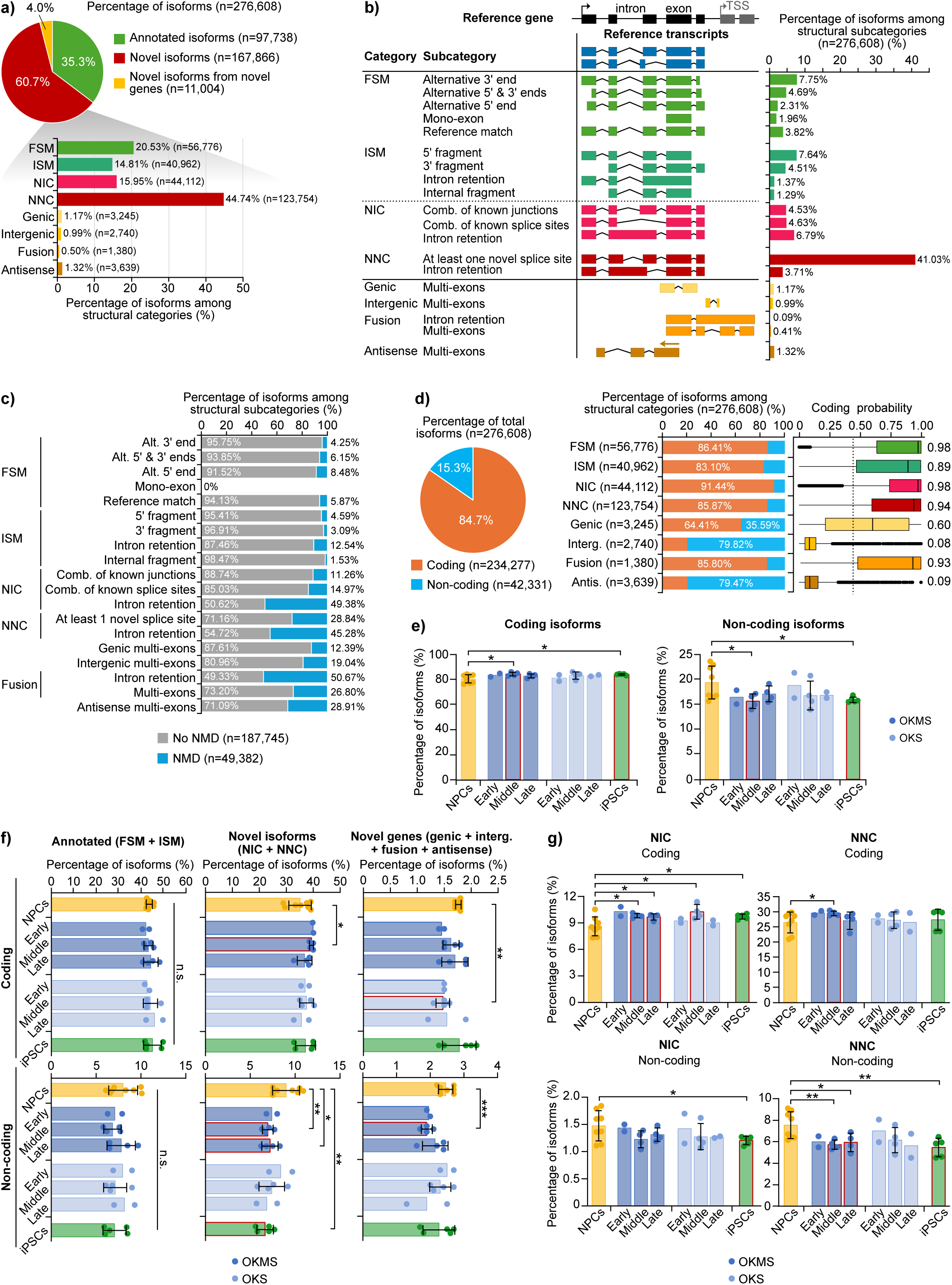
lrRNAseq reveals unprecedented isoform diversity and tight regulation of non-coding isoforms during reprogramming. **a)** Proportions of known and novel isoforms and of each isoform structural categories in the generated dataset after filtering out the transcripts lacking a poly(A) motif or with reverse-transcription (RT) switching events. FSM: Full-Splice Matches; ISM: Incomplete Splice Matches; NIC: Novel-In-Catalog; NNC: Novel-Not-in-Catalog. **b)** Proportions of isoforms in each structural subcategory, according to SQANTI3 classification^55^.TSS: Transcription Start Site; Comb.: Combination. **c)** Proportions of coding isoforms going through or avoiding nonsense-mediated decay (NMD) among the different structural subcategories of the filtered dataset. Alt.: Alternative. **d)** Proportions of coding and non-coding isoforms detected in the filtered dataset (pie chart), or separated by isoform structural category (bar graph), and box plots showing the coding probability of these isoforms. Central lines represent medians and boxes denote quartiles. Error bars represent the minimum and maximum values. Dots represent outlier cells. Values on the side of the box plots represent the median value of coding probabilities. Vertical dotted line represents the coding probability optimum cut-off indicating coding transcripts in mouse, according to the CPAT tool^129^. Interg.: Intergenic; Antis.: Antisense. **e)** Mean percentages of coding and non-coding isoforms detected in the NPCs (n=8), during early (n=2), middle (n=4) or late (n=4 for OKMS and n=2 for OKS) phases of reprogramming, or in the iPSCs (n=3) of the dataset, in total or **f)** separating annotated from novel isoforms. Error bars represent the standard deviation (STD) and each dot represents the replicates. Red outlines highlight significant data. P-values were calculated comparing to NPCs using two-sided t-test with unequal variance. Non significant (n.s.) ≥ 0.05, * < 0.05, ** < 0.01, *** < 0.001. **g)** Mean percentages of coding and non-coding NIC or NNC isoforms detected in NPCs (n=8), during early (n=2), middle (n=4) or late (n=4 for OKMS and n=2 for OKS) phases of reprogramming, or in the iPSCs (n=3) of the dataset. Error bars represent the STD and each dot represents replicates. Red outlines highlight significant data. P-values were calculated comparing to NPCs using two-sided t-test with unequal variance. * < 0.05, ** < 0.01.

Investigating subcategories of isoforms (**Fig. 2b**), we found that NNC isoforms with at least one novel splice site constituted the main subcategory (41.03%), followed by isoforms with shorter 3’ end (FSM *alternative 3’ end* and ISM *5’ fragment* subcategories, at 7.75% and 7.64%, respectively) and isoforms with *retained introns* (11.96% in total) (**Fig. 2b**). Interestingly, on average 48% of isoforms classified as having intron-retention were predicted to undergo nonsense-mediated decay (NMD) (NIC (49.38%), NNC (45.28%) and fusion (50.67%) *intron retention* subcategories) (**Fig. 2c** and **Supplementary Data 2**). This is consistent with the role of intron retention in NMD^57^, but more importantly, it recapitulates what we previously observed as a role for intron retention in regulating gene expression during reprogramming^16^. Taken together, our results highlight the importance of lrRNAseq in identifying new isoforms and confirm previous findings that the transcriptome undergoes significant splicing and RNA processing events during the reprogramming process, but most importantly, they suggest that isoform diversity has precedingly been underestimated^31,35^.

We next analyzed for coding potential and found that the proportion of non-coding isoforms was significantly lower than coding isoforms (15.3% compared to 84.7%, respectively) (**Fig. 2d** and **Supplementary Data 2**), and this is was especially true for known genes (FSM: 86.41%; ISM: 83.10%; NIC: 91.44%, and NNC: 85.87%, with coding probability medians between 0.89 and 0.98). Moreover, non-coding isoforms came predominantly from novel genes, with intergenic and antisense isoforms showing a very low coding score (median of 0.08 and 0.09, respectively, **Fig. 2d**), and almost 80% of these isoforms being non-coding (79.82% and 79.47%, respectively). This is consistent with intergenic regions generally harboring lncRNAs^58,59^ and with most antisense genes being transcribed as lncRNAs^60,61^. Given that the majority of non-coding isoforms arise mainly from the relatively small number of novel genes, this suggests that for known genes there is a preference for coding isoforms and a selection against non-coding isoforms.

To determine how the dynamics of non-coding isoform diversity is affected during reprogramming, we segregated isoforms by reprogramming phase, based on the groupings described in **Fig. 1c** (**Supplementary Data 2**). Overall, we found a significant decrease of the proportion of non-coding isoforms in the middle phase of OKMS reprogramming, which was also reflected in the final iPSC stage (**Fig. 2e**). Conversely, the proportion of coding isoforms increased. These opposing changes were mostly driven by novel isoforms from both known and novel genes (i.e. NIC, NNC, and novel genes) but not annotated genes, with a striking decrease in non-coding NNC isoforms (**Fig. 2f-g**). This was not due to bias introduced by the use of different types of sequencing platforms or flow cell technologies (i.e. MinION vs PromethION, and R9 vs R10 flow cells, respectively), as the bias was only limited to novel but not known genes (**Supplementary Fig. 6c-d**), and the majority of novel isoforms come from known genes. Altogether, our data expand on previous knowledge by characterizing for the first time both coding and non-coding isoforms in the context of reprogramming. Our results suggest that there is tight and controlled regulation of isoform diversity during reprogramming with a selection against novel non-coding transcripts.

### Alternative splicing follows a biphasic pattern during reprogramming

As NNC isoforms are transcripts that use novel donor and/or acceptor splice sites to create new splicing junctions, we aimed to investigate whether the observed changes in the abundance of non-coding transcripts could be explained by post-transcriptional regulation mechanisms such as alternative splicing (AS). Indeed, AS has been demonstrated as one of the underlying mechanisms affecting reprogramming efficiency, with different RBPs coordinating AS events at each timepoint of the process^32^. Thus, we first assessed expression of 19 splicing regulators known to modulate alternative splicing in pluripotent stem cells and reprogramming^32,35^, splitting them into 4 groups based on their expression: Increased expression during reprogramming to pluripotency (group 1), expressed throughout (group 2), decreased (group 3), or transiently increased (group 4) (**Fig. 3a** and **Supplementary Data 7**). The expression profiles of these RBPs did not globally differ when comparing the two reprogramming systems, although it highlighted again delayed kinetics in OKS reprogramming compared to the OKMS system, and was consistent with previous studies^31,32,35,62^. For examples, *Tia1*, *Mbnl1* and *Mbnl2* expression was down-regulated (group 3) in both systems, which agrees with their known function as negative regulators of ESC-like AS patterns^32,62,63^. Similarly, *Esrp1* and *Esrp2* expression transiently peaked between D6 and D10 for OKMS, and between D18 and D20 for OKS (group 4), following the same pattern as observed with markers of the MET (**Fig. 1b**). This is in line with their known role in promoting AS changes at the MET stage of reprogramming^32,64^. Thus, our results recapitulate the expression changes of regulators known to coordinate temporal AS patterns observed during induction to pluripotency.

**Figure 3:**
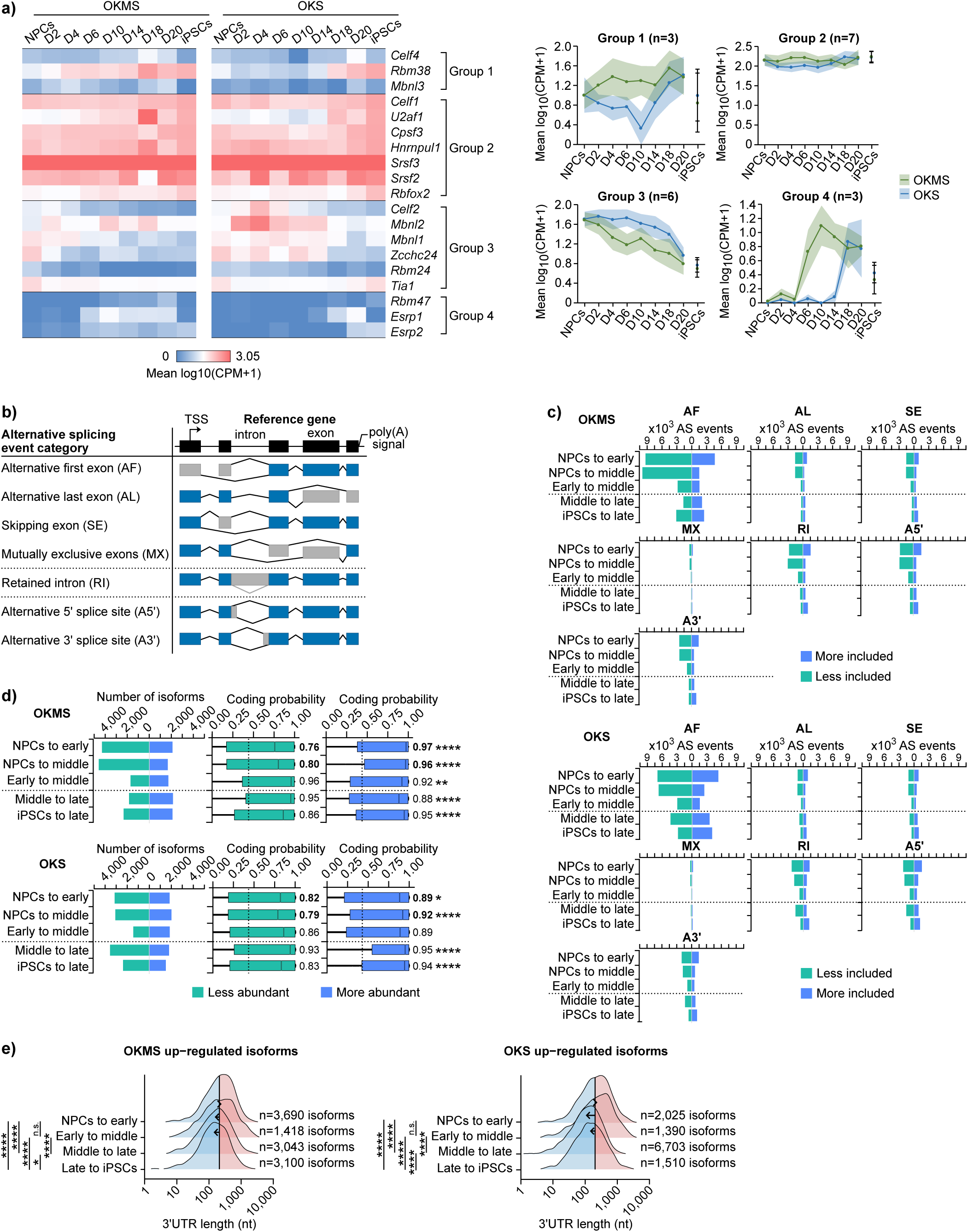
Alternative splicing follows a biphasic pattern during reprogramming. **a)** Heat map and plots depicting logged expression of pluripotency-related regulators of alternative splicing (gene lists from Cieply *et al.*^32^ and Vivori *et al.*^35^) in OKMS or OKS reprogramming. Dots and lines represent average expression per timepoint, and shaded areas and error bars represent the SEM. **b)** Schematic of the different types of local alternative splicing events analyzed in the filtered dataset, according to SUPPA2 classification^142^. **c)** Splicing events-based quantification of alternative splicing significantly more or less included, or **d)** isoform-based quantification of alternative splicing significantly more or less abundant, during OKMS and OKS reprogramming phase transitions (NPCs (n=8), early (n=2), middle (n=4), late (n=4 for OKMS and n=2 for OKS) phases of reprogramming, iPSCs (n=5)). Significance based on differential percent spliced-in (ΔPSI) ≥ 0.1 (more included) or ΔPSI ≤ -0.1 (less included) and an adjusted p-value ≤ 0.05 (P-values were calculated using one-sided t-test with unequal variance based on the direction of change, ex. more versus less included). Horizontal dotted line represents the inflexion point in the biphasic alternative splicing pattern observed in reprogramming. AF: Alternative First exon; AL: Alternative Last exon; SE: Skipped Exon; AS: Alternative Splicing; MX: Mutually exclusive exons; RI: Retained Intron; A5’: Alternative 5’ splice site; A3’: Alternative 3’ splice site. Box plots represent the coding probability for significant alternatively spliced isoforms at timepoint transitions in OKMS or OKS reprogramming. Values to the right of box plots represent the average coding probabilities. Vertical dotted lines represent the coding probability optimum cut-off indicating coding transcripts in mouse, according to the CPAT tool^129^. Central lines represent medians and boxes denote quartiles. Error bars represent the minimum and maximum values. Dots represent outlier cells. P-values were calculated using two-sided t-test with unequal variance. * < 0.05, ** < 0.01, **** < 0.0001. **e)** Distribution of 3’UTR lengths (log scale) in up-regulated isoforms from genes subject to alternative splicing during OKMS and OKS reprogramming phase transitions (NPCs (n=8), early (n=2), middle (n=4), late (n=4 for OKMS and n=2 for OKS) phases of reprogramming, iPSCs (n=5)). Lines represent means of the NPCs to early datasets. Arrow points show change in distribution means in each transition. P-values were calculated by ordinary one-way ANOVA with Tukey HSD *post hoc* test. n.s. ≥ 0.05, * < 0.05, **** < 0.0001. UTR: UnTranslated Region; nt: nucleotides.

To further investigate splicing regulation, we assessed AS events (defined in **Fig. 3b**) during reprogramming. For this, we analyzed our 276,608 identified isoforms (**Supplementary Fig. 2** and **Supplementary Data 2**) and removed events occurring when comparing NPCs to FBSL samples, which represent changes due to media change only, to focus on events only induced by reprogramming factors (**Supplementary Fig. 7a-b**). With this in mind, we detected a total of 140,645 AS and 33,657 differential isoforms usage events occurring throughout the reprogramming process (**Supplementary Fig. 2** and **Supplementary Data 7**). The majority of AS events, with alternative first exons dominating the events, occurred when NPCs transitioned to the early stages of reprogramming (n=13,816 for OKMS; n=12,166 for OKS), irrespective of the reprogramming system (**Fig. 3c**). AS events were less included during the early stages of reprogramming (NPCs to middle stage: A first phase), whereas the inclusion of splicing variations increased steadily in the later stages (**Fig. 3c**) (middle to late/iPSC stages: A second phase), suggesting a biphasic pattern that switches in the middle of reprogramming to include splicing events.

To investigate if these reprogramming specific AS events could lead to functional consequences, we performed an isoform switching (IS) analysis. Isoform switching can inform of AS events that lead to open reading frame (ORF) disruptions or changes that affect protein expression. IS can be measured by calculating differential isoform usage using differential isoform percentage spliced-in (ΔPSI) events (“ΔPSI events-based method”) or TPM values to obtain differentially expressed isoforms (DEIs) (“DEIs-based method”), where the number of ΔPSI events or DEIs is proportionally indicative of the number of IS events. Using the ΔPSI events-based method, we found that IS peaked early during reprogramming (NPCs to early/mid-reprogramming), with a higher number of less abundant events (∼4,000) than more abundant events (∼2,000) (**Fig. 3d**). Similar to AS events, IS was also biphasic, changing from less abundant in the early stages of reprogramming to more abundant in later stages (**Fig. 3d**). Interestingly, less abundant isoforms in early and middle phase reprogramming showed lower coding probabilities (i.e. decreased or disrupted ORFs) than more abundant isoforms (**Fig. 3d**). This is consistent with the specific decrease in non-coding isoforms in the middle phase of reprogramming (**Fig. 2e-g**). In parallel, a global decrease was observed in 3’ untranslated region lengths for alternatively spliced isoforms that are up-regulated in those later reprogramming stages (**Fig. 3e**, **Supplementary Fig. 7c-d**, **Supplementary Data 2**, and **Supplementary Data 8**), which has been shown to be associated with higher protein-coding potential^65^. Taken together, these results suggest that AS leads to gain of protein-coding potential and function, and tight regulation of non-coding transcripts in the early phases of reprogramming.

### Selective interplay between gene expression and splicing regulates reprogramming

Having identified major splicing and isoform usage changes during reprogramming, especially in the early phases, we investigated how this affects isoform expression and gene function, and whether common or different genes are affected in other datasets. In total, we identified 10,460 genes that undergo splicing changes during reprogramming, and of these 49 genes (among them *Tpm1, Mbnl2, Zfp207, Fgfr1, Palm, Fn1, Dst, Fyn*) were found in common between our dataset and two other reprogramming datasets^32,35^ (**Fig. 4a** and **Supplementary Data 7**). Strikingly, we found that close to 90% of the genes identified as differentially spliced in these datasets overlapped with our dataset, whether the cells used for reprogramming were NPCs (our study), MEFs (637 genes out of 676, i.e. 94.2%) or B cells (131 genes out of 148, i.e. 88.5%). However, less than 7% of the genes undergoing AS we detected were found in the two other studies. Thus, our results highlight the strength of using lrRNAseq to detect novel isoforms and splicing events compared to other technologies.

**Figure 4:**
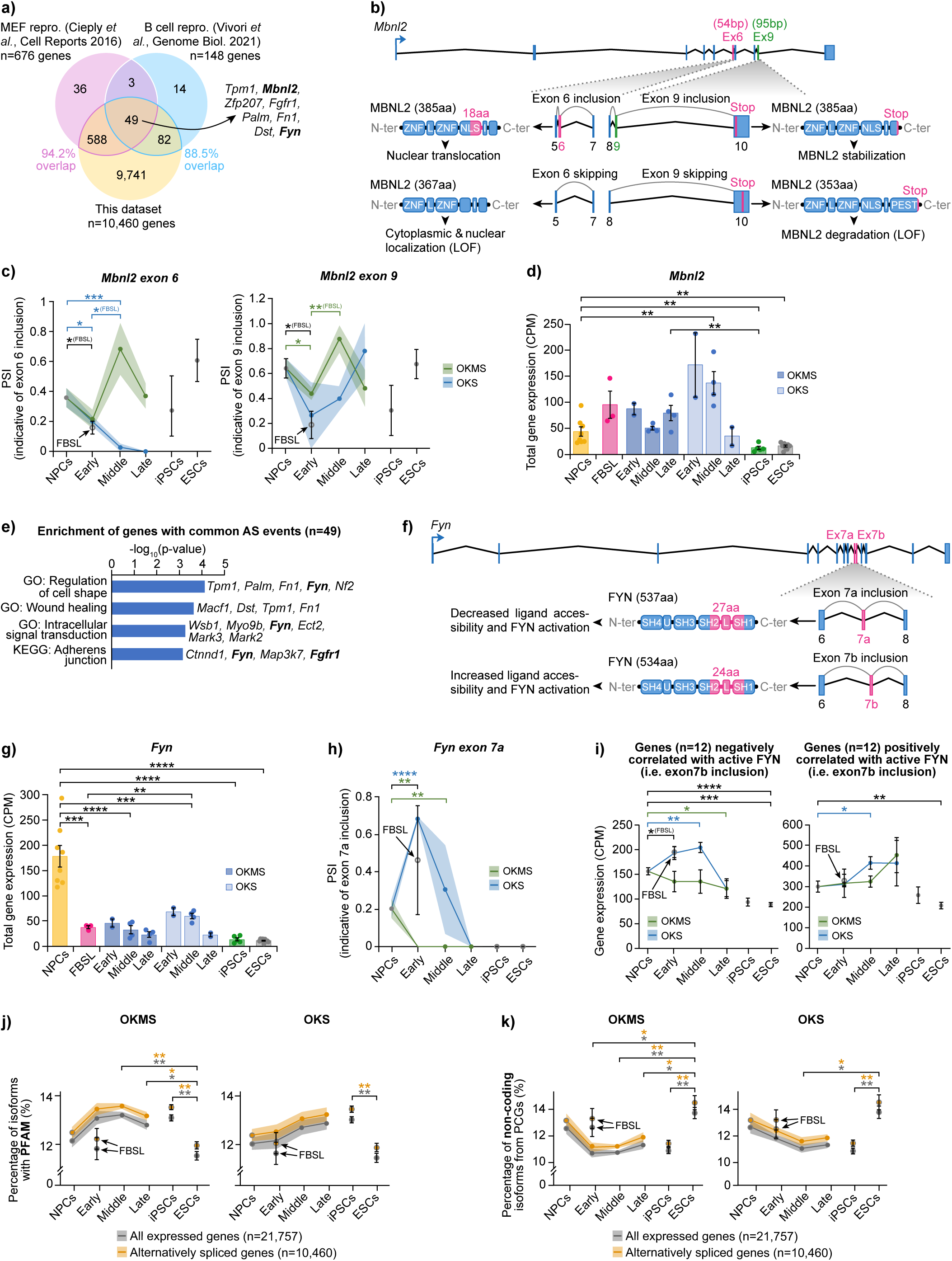
Selective interplay between gene expression and alternative splicing regulates reprogramming. **a)** Overlap between genes undergoing any significant alternative splicing detected from our lrRNAseq dataset (reprogramming from NPCs) and the datasets from Cieply *et al.*^32^ (reprogramming from MEFs) and Vivori *et al.*^35^ (reprogramming from B cells). MEF: Mouse Embryonic Fibroblast. **b)** Schematic highlighting two alternative splicing events of exons 6 and 9 within the *Mbnl2* gene locus, and their respective consequences. Ex: Exon; aa: amino acids; N/C-ter: N-/C-terminus; ZNF: Zinc Finger domain; L: Linker domain; NLS: Nuclear Localization Signal; I: Interaction domain; LOF: Loss-Of-Function; PEST: PEST domain. **c)** PSI values of *Mbnl2* exon 6 or exon 9 inclusion in NPCs (n=8), FBSL (n=3), during early (n=2), middle (n=4) or late (n=4 for OKMS and n=2 for OKS) phases of reprogramming, or in the iPSCs (n=5) and ESCs (n=9) of the dataset. Dots and lines represent average expression per timepoint, and shaded areas and error bars represent the standard error of the mean (SEM). P-values were calculated using one-sided t-test with unequal variance to ask if the exon is included or not. * < 0.05, ** < 0.01, *** < 0.001. The data is the same in OKMS and OKS for NPCs, FBSL, iPSCs and ESCs samples because those samples were in common for OKMS and OKS (grey dots). **d)** Mean expression of *Mbnl2* gene in NPCs (n=8), FBSL (n=3), during early (n=2), middle (n=4) or late (n=4 for OKMS and n=2 for OKS) phases of reprogramming, or in the iPSCs (n=5) and ESCs (n=9) of the dataset. Error bars represent the SEM and each dot represents replicates. P-values were calculated using one-sided t-test with unequal variance. ** < 0.01. **e)** Logged p-values from GO-term enrichment analysis (biological processes and KEGG pathways) for the genes commonly spliced between our lrRNAseq dataset and the datasets from Cieply *et al.*^32^ and Vivori *et al.*^35^. P-values were calculated by one-sided Fisher’s exact test. FDR ≤ 0.1. **f)** Schematic highlighting the mutually exclusive alternative splicing of exons 7a and 7b within the *Fyn* gene locus, and their respective consequences. SH: Src Homology domain; U: Unique domain. **g)** Mean expression of *Fyn* gene in NPCs (n=8), FBSL (n=3), during early (n=2), middle (n=4) or late (n=4 for OKMS and n=2 for OKS) phases of reprogramming, or in the iPSCs (n=5) and ESCs (n=9) of the dataset. Error bars represent the SEM and each dot represents replicates. P-values were calculated using one-sided t-test with unequal variance. ** < 0.01, *** < 0.001, **** < 0.0001. **h)** PSI values of *Fyn* exon 7a inclusion in NPCs (n=8), FBSL (n=3), during early (n=2), middle (n=4) or late (n=4 for OKMS and n=2 for OKS) phases of reprogramming, or in the iPSCs (n=5) and ESCs (n=9) of the dataset. Dots and lines represent average expression per timepoint, and shaded areas and error bars represent the standard error of the mean (SEM). P-values were calculated using one-sided t-test with unequal variance to ask if the exon is included or not. ** < 0.01, **** < 0.0001. The data is the same in OKMS and OKS for NPCs, FBSL, iPSCs and ESCs samples because those samples were in common for OKMS and OKS (grey dots). **i)** Mean expression of genes that negatively or positively correlate with *Fyn* in CRISPR screens from DepMap Public 25Q3 Release dataset^69^, during the phases of OKMS and OKS reprogramming. Error bars represent the SEM. The data is the same in OKMS and OKS for NPCs, FBSL, iPSCs and ESCs samples because those samples were in common for OKMS and OKS (grey dots). P-values were calculated using two-sided t-test with equal variance. * < 0.05, ** < 0.01, *** < 0.001, **** < 0.0001. **j)** Percentages of isoforms with annotated PFAM domains or **k)** percentages of non-coding isoforms from protein-coding genes, for either all expressed genes, or genes that undergo alternative splicing, in NPCs (n=8), FBSL (n=3), during early (n=2), middle (n=4) or late (n=4 for OKMS and n=2 for OKS) phases of reprogramming, or in the iPSCs (n=5) and ESCs (n=9) of the dataset. Dots and line represent average and shaded area represent the SEM for reprogramming samples. For the NPCs, FBSL, iPSCs and ESCs samples, error bars represent the SEM. The data is the same in OKMS and OKS for NPCs, FBSL, iPSCs and ESCs samples because those samples were in common for OKMS and OKS (grey circles). P-values were calculated by ordinary one-way ANOVA with Tukey HSD *post hoc* test. n.s. ≥ 0.05, * < 0.05, ** < 0.01. PCGs: Protein-Coding Genes.

Of the 49 common genes, we found that *Mbnl2*, a negative regulator of AS in pluripotency^32,62,63^ (**Fig. 3a**), undergoes two exon inclusion/skipping events during reprogramming: Inclusion of exon 6 yields a bipartite nuclear localization signal, thus exon 6 skipping can affect the localization and function of MBNL2^66^. Similarly, skipping exon 9 causes a frameshift and decreased protein stability^66^ (**Fig. 4b** and **Supplementary Data 7**). Interestingly, exons 6 and 9 were specifically skipped in both OKMS and OKS reprogramming from the early phase (**Fig. 4c** and **Supplementary Fig. 8a**), suggesting loss of function of MBNL2. Those exons were then included in OKMS mid-reprogramming, and the global gene expression remained stable (**Fig. 4c-d**). As for OKS, expression strongly decreased in the late phase, suggesting that different mechanisms of gene regulation are selectively used between the two systems (i.e. splicing versus expression changes in OKMS and OKS, respectively). Other commonly spliced genes are involved in cell signaling and adhesion (**Fig. 4e** and **Supplementary Data 7**). Among them are negative regulators of pluripotency, such as tyrosine kinases *Fgfr1* and *Fyn* (**Fig. 4a** and **Fig. 4e**), which promote differentiation of ESCs^67^. *Fyn*, a SRC family kinase, contains a mutually exclusive exon 7 (exons 7a and 7b), that leads to differences in FYN SH2 linker and N-terminal of the catalytic domain SH1 (**Fig. 4f** and **Supplementary Data 7**). Inclusion of exon 7a decreases FYN accessibility to its ligand and thus results in lower activation of the kinase^68^. We found that both decrease in gene expression and inclusion of exon 7a could lead to a decrease in FYN functionality during reprogramming, more specifically OKS reprogramming (**Fig. 4g-h**, **Supplementary Fig. 8b-c**, and **Supplementary Data 7**). Interestingly, expression of genes negatively-correlated with active *Fyn* (based on DepMap^69^ analyses) increased in the early and middle phases of reprogramming following the inclusion of exon 7a in OKS (i.e. gain of inactive form of FYN, thus behaving like loss of FYN). Comparatively, no significant changes were observed in OKMS, consistent with the low exon 7a inclusion, but there is a decreasing trend in the expression of those FYN targets (**Fig. 4i**, left and **Supplementary Data 7**). As for positively-correlated genes, we noticed an increase mostly in the middle and late phases of OKS and OKMS reprogramming, respectively, where only the active form of FYN is expressed (i.e. exclusion of exon 7a and inclusion of exon 7b) (**Fig. 4g-h** and right panel of **Fig. 4i**). These results suggested that expression of an inactive isoform of FYN at the beginning of reprogramming corresponds accordingly with the regulation of its target genes.

To investigate possible functional effects of alternative splicing on a global level during reprogramming, we assessed the proportion of isoforms that included PFAM protein domains, as well as non-coding isoforms that were generated from protein-coding genes (**Fig. 4j-k** and **Supplementary Data 8**). We observed an increasing trend in isoforms containing annotated PFAM domains throughout both the OKMS and OKS reprogramming timelines (**Fig. 4j**), which is consistent with the higher coding potential of more abundant isoforms versus less abundant isoforms during reprogramming (**Fig. 3d**). Inversely, the proportion of non-coding isoforms from protein-coding genes decreased significantly during reprogramming (**Fig. 4k**). This pattern was true for all expressed genes and genes undergoing alternative splicing. Taken together, our results are consistent with the notion of selection against non-coding isoforms during reprogramming and suggest that a broader selection of functional isoforms (i.e. isoforms containing PFAM domain) is at play to ensure proper reprogramming to pluripotency. This again highlights the strength of using lrRNAseq to identify isoforms which are functionally relevant for studying specific biological processes.

### Isoform switching selectively regulates non-coding isoforms during early reprogramming

Given that both expression and AS selectively regulate reprogramming, we sought to determine if these splicing patterns affected isoform expression. From the 276,608 identified transcripts, we focused only on those that are differentially expressed in at least one sample comparison in our dataset. This refined the list to 195,889 isoforms (**Supplementary Fig. 2** and **Supplementary Data 9**), which are significantly more or less expressed during the different phases of reprogramming (**Fig. 5a**). Similar to our splicing analysis, we excluded isoforms affected by media change only (FBSL to NPCs comparison, **Supplementary Fig. 9a**). Similar to differentially expressed genes (**Supplementary Fig. 1c**), we observed a 2-wave pattern of DEIs, where more isoforms are up-regulated in the early and lates phases of reprogramming (much more pronounced in OKS system) and where there are less DEIs between the early and middle phases of the process (**Fig. 5a**). This held true for both coding and non-coding isoforms (**Supplementary Fig. 9b**). This also suggests that the early phases of reprogramming are in part conserved in terms of isoform expression regulation and coincides with the fact that several molecular and cellular mechanisms important for reaching the pluripotent state take place in the early steps of reprogramming^41,45,70,71^. Therefore, we narrowed our subsequent analyses to isoform expression changes occurring as the cells progress from NPCs to mid-reprogramming. We classified isoform changes into 4 categories based on their expression profiles: Up-regulated (“up”), down-regulated (“down”), up-then down-regulated (“up-down”), or down-then up-regulated (“down-up”) (**Supplementary Fig. 9c**). This reduced our list of DEIs to 35,262 transcripts (18% of total DEIs, corresponding to 12,428 genes) (**Supplementary Fig. 2** and **Supplementary Data 9**) but also allowed to define and track isoform expression patterns. For example, we found that isoforms that were up-regulated in early to mid-reprogramming were later down-regulated in late reprogramming (n=9,516 for OKMS; n=6,872 for OKS) (**Supplementary Fig. 9c**).

**Figure 5:**
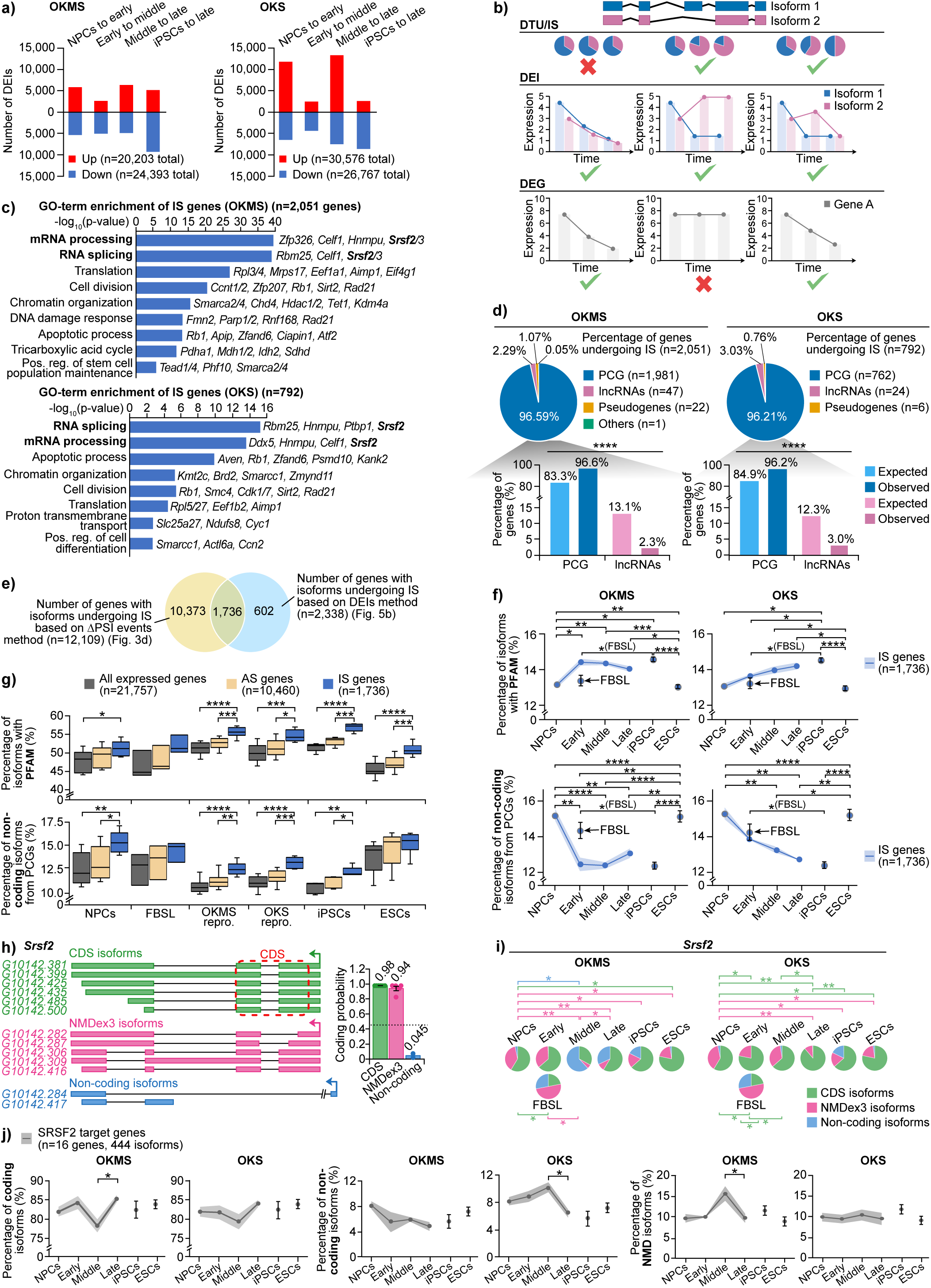
Isoform switching has functional impacts on reprogramming. **a)** Number of up- or down-regulated isoforms between the different phases of reprogramming among the total isoforms (n=195,889) detected in the dataset. Significance based on log2(fold change) ≥ 2 for up-regulated isoforms or ≤ 0.5 for down-regulated isoforms with adjusted p-values ≤ 0.05 (calculated by Benjamini-Hochberg method). DEIs: Differentially Expressed Isoforms. **b)** Schematic defining the difference between differential transcript usage (i.e. isoform switching) and differential isoform or gene expression. DTU/IS: Differential Transcript Usage/Isoform switch; DEG: Differentially Expressed Gene. **c)** Logged p-values from GO-term enrichment analysis (biological processes) for genes undergoing isoform switch (IS) between D0 and the middle phase of OKMS or OKS reprogramming. P-values were calculated by one-sided Fisher’s exact test. FDR ≤ 0.1. Pos. reg.: Positive regulation. **d)** Proportions of gene biotype among the genes undergoing isoform switching, according to GENCODE annotation^145^. Bar graphs depict the observed percentage of protein-coding genes or lncRNA genes undergoing isoform switch, compared to their expected percentages found for genes undergoing differential isoform expression. P-values were calculated by Chi-squared test. **** = 5.3x10^-53^ (OKMS), 7.3x10^-17^ (OKS). lncRNAs: long non-coding RNAs. **e)** Overlap between the number of genes which undergo isoform switch between D0 and the middle phase of our reprogramming systems, based on two methods. **f)** Percentages of differentially expressed isoforms with annotated PFAM domains or non-coding isoforms from protein-coding genes, for genes that undergo isoform switching, in NPCs (n=8), FBSL (n=3), during early (n=2), middle (n=4) or late (n=4 for OKMS and n=2 for OKS) phases of reprogramming, or in the iPSCs (n=5) and ESCs (n=9) of the dataset. Dots and line represent average and shaded area represent the SEM for reprogramming samples. For the FBSL, iPSCs and ESCs samples, error bars represent the SEM. The data is the same in OKMS and OKS for NPCs, FBSL, iPSCs and ESCs samples because those samples were in common for OKMS and OKS (grey circles). P-values were calculated by ordinary one-way ANOVA with Tukey HSD *post hoc* test. * < 0.05, ** < 0.01, *** < 0.001, **** < 0.0001. **g)** Box plots of percentages of isoforms with annotated PFAM domains or the percentage of non-coding isoforms, for either all expressed genes, alternatively spliced genes (AS), and differentially expressed isoforms of genes that undergo isoform switching, in NPCs (n=8), FBSL (n=3), during early (n=2), middle (n=4) or late (n=4 for OKMS and n=2 for OKS) phases of reprogramming, or in the iPSCs (n=5) and ESCs (n=9) of the dataset. Bounds of the box represent 25^th^ and 75^th^ quartiles, bounds of the whiskers represent minimum and maximum values, and center line represents median. P-values were calculated by ordinary one-way ANOVA with Tukey HSD *post hoc* test. * < 0.05, ** < 0.01, *** < 0.001, **** < 0.0001. **h)** Schematic depicting the transcripts of the *Srsf2* gene detected in our dataset, as visualized with IGV software^144^, and bar graph showing the coding probability of the *Srsf2* transcript categories (n=6 CDS isoforms; n=5 NMDex3 isoforms; n=2 non-coding isoforms). Horizontal dotted line represents the coding probability optimum cut-off indicating coding transcripts in mouse, according to the CPAT tool^129^. Error bars represent the SEM and each dot represents replicates. CDS: Coding Sequence. **i)** Proportions of each *Srsf2* transcript category in OKMS and OKS reprogramming phases. P-values were calculated using one-sided t-test with unequal variance. * < 0.05, ** < 0.01. The data is the same in OKMS and OKS for NPCs to FBSL and ESCs to iPSCs comparisons because those samples were in common for OKMS and OKS. **j)** Percentage of SRSF2 target genes (gene list from Han *et al*.^72^) expression coming from coding isoforms, non-coding isoforms or isoforms that undergo nonsense-mediated decay in NPCs (n=8), during early (n=2), middle (n=4) or late (n=4 for OKMS and n=2 for OKS) phases of reprogramming, or in the iPSCs (n=5) and ESCs (n=9) of the dataset. Dots and line represent average expression and shaded areas represent SEM. For the iPSCs and ESCs samples, error bars represent the SEM. P-values were calculated for each successive phases using a two-sided t-test. * = 0.019 (coding OKMS), 0.041 (non-coding OKS), 0.046 (NMD OKMS).

With this refined DEIs list and given that both gene expression and AS are selectively regulated as we previously showed, we now sought to integrate the notion of isoform expression with proportionality in our IS analysis (i.e. “DEIs-based method”). This is different from our ΔPSI event-based IS analysis (**Fig. 3d**), which mainly represents isoform proportions. For this we devised an in-house matrix to recapitulate the different possible scenarios of IS occurring between NPCs and mid-reprogramming (**Fig. 5b** and IS matrix in **Supplementary Data 9**). We found 2,051 and 792 genes undergoing IS for OKMS and OKS, respectively, affecting about 16.5% and 6.4% of genes where isoforms are differentially expressed between NPCs and mid-reprogramming (**Supplementary Fig. 9d** and **Supplementary Data 9**). GO-term enrichment analysis showed that genes undergoing IS between NPCs and mid-reprogramming for both OKMS and OKS were associated with processes such as mRNA processing, RNA splicing, apoptosis, cell division, DNA repair, chromatin remodeling, translation, cellular respiration and stem cell population maintenance and differentiation (**Fig. 5c**, **Supplementary Fig. 9d**, and **Supplementary Data 9**). This is expected as these biological mechanisms are key actors in induction of pluripotency. Thus, isoform switching is involved in every layer of pluripotency induction regulation, and this is conserved in our two different reprogramming systems.

Interestingly, while no differences in the patterns of DEIs could be detected between coding and non-coding genes (**Supplementary Fig. 9b**), we observed a striking lack of non-coding genes undergoing IS (**Fig. 5d**). Indeed, around 96% of IS genes corresponded to protein-coding genes, whereas only about 2-3% were lncRNA genes, and around 1% constituted pseudogenes or other biotypes. We compared these numbers to the proportion of biotypes in the list of genes undergoing differential isoform expression but not necessarily IS (n=12,347 genes for OKMS; n=8,350 genes for OKS) and found that lncRNAs were significantly less enriched within the IS gene list: lncRNAs were expected to amount for 12-13% of genes, but they represented only 2-3% in our datasets for IS (**Fig. 5d**). In contrast, protein-coding genes were significantly more enriched within the IS gene list (96.6% for OKMS; 96.2% for OKS) compared to expected values (83.3% for OKMS; 84.9% for OKS). These results suggest that there is a negative constraint against lncRNAs to undergo IS, consistent with our results demonstrating a constraint and tight regulation of non-coding transcripts during the early phases of reprogramming (**Fig. 2e-g**).

When looking at genes classified as undergoing IS by both the ΔPSI events-based and DEIs-based methods of IS analysis (n=1,736 genes) (**Fig. 5e** and **Supplementary Fig. 2**), we observed that, similar to the previous patterns for all genes (**Fig. 4j-k**), reprogramming is associated with an increase in the proportion of isoforms containing annotated protein domains (**Fig. 5f**), and a concomitant and significant decrease in the number of non-coding isoforms. Interestingly, genes subject to IS exhibited higher proportions of both isoforms with annotated protein domains (i.e. putative functional isoforms) and non-coding isoforms across reprogramming samples compared to all expressed genes or genes subject to AS (**Fig. 5g**). This suggests that IS allows for specific selection of both functional coding isoforms, encoding annotated protein domains, and non-coding isoforms from protein-coding genes.

One example of a protein-coding gene with demonstrated non-coding isoforms regulation during reprogramming is *Srsf2. Srsf2* is a well-known regulator of AS in early reprogramming^31,72^, and was among the genes in the main GO-terms associated with IS (i.e. “RNA splicing” and “mRNA processing”) (**Fig. 5c**). *Srsf2* gene-level expression did not significantly change during both OKMS and OKS reprogramming but tended to increase when cells reach the pluripotent state (**Supplementary Fig. 9e**). We identified 13 DEIs for *Srsf2*, six coded for the proper protein, five were predicted to undergo NMD while retaining a strong coding probability, and two were non-coding (**Fig. 5h**). Of the five NMD transcripts (NMDex3), three were NIC (new combinations of known splice junctions) isoforms that included a splicing in exon 3 that was distinct from the protein-coding (main) isoforms (**Fig. 5h** and **Supplementary Data 10**). We compared expression profiles and fractions of the main, NMDex3, and non-coding isoforms. In OKMS reprogramming, the proportion of the protein-producing isoforms and the NMDex3 isoforms decreased significantly (going from 58.6% and 38.1% in NPCs to 32.9% and 5.2% in middle phase, respectively) (**Fig. 5i** and **Supplementary Fig. 9f-g**), when their expression levels were not significantly altered. This was due to a significant increase in the expression and thus proportion of the non-coding isoforms (going from 3.3% in NPCs to 61.9% in middle phase) (**Fig. 5i**, **Supplementary Fig. 9f-g**, and **Supplementary Data 10**). This pattern was reversed in the late and iPSC phases of OKMS reprogramming. As for OKS reprogramming, this time the non-coding isoforms remained very lowly expressed throughout the process (**Supplementary Fig. 9g**). However, expression of the NMDex3 isoforms increased until the middle phase of reprogramming, regulating the proportion of functional isoforms. In later phases, the proportion of the NMDex3 isoforms decreased significantly (going from 38.1% in NPCs to 10.5% in the late phase), thus allowing an increase in proportion of the main protein-coding isoforms (going from 58.6% in NPCs to 88.8% in late phase) (**Fig. 5i**, **Supplementary Fig. 9f-g**, and **Supplementary Data 10**). Thus, in both reprogramming systems, isoforms leading to the proper SRSF2 protein dominate later in reprogramming through expression control of different isoforms, and more specifically for OKMS by IS of the non-coding isoforms to the benefit of the main coding isoforms. A similar effect could be observed when analyzing reprogramming from B cells with OSKM reprogramming factors (data from Stadhouders *et al.*^73^). Indeed, a significant increase in the proportion of isoforms leading to the proper SRSF2 protein can be seen throughout reprogramming, accompanied by a significant increase of the non-coding isoforms in the early phase, and their decrease in late reprogramming (**Supplementary Fig. 9h** and **Supplementary Data 10**). This suggests that this phenomenon is conserved across reprogramming systems from multiple differentiated cell types. To assess whether the variations in *Srsf2* isoform levels could have an impact on the role of this splicing regulator, we investigated the fate of known SRSF2 target gene transcripts^72^ in our dataset (**Supplementary Data 10**). The first half of reprogramming was associated with an increase in non-functional isoforms (whether non-coding or NMD isoforms) for SRSF2 target mRNAs in both reprogramming systems (**Fig. 5j**, **Supplementary Data 2**, and **Supplementary Data 8**), which was consistent with a diminished proportion of protein-coding isoforms. Conversely, increase of functional *Srsf2* isoforms in late reprogramming occurred in the same phase as the decrease in the non-functional isoforms of SRSF2 target mRNAs (**Fig. 5i-j**). Overall, these results suggest again that selective regulation through AS and isoform switching allows proper control of genes important for proper reprogramming and for the resulting iPSCs.

This regulation of genes by expression of non-coding isoforms is also consistent with changes in coding probability decreasing in the middle phase of reprogramming, to then increase in later phases as observed in our ΔPSI-based IS analysis (**Fig. 3d**). This also offers interesting parallels with the specific regulation of non-coding isoforms from both coding genes (**Fig. 5f-g**) and lncRNAs (**Fig. 5d**) observed during reprogramming and suggests that coding and non-coding isoforms are regulated differently and at different phases of reprogramming. With this in mind, and because we previously showed how during reprogramming IS can select for isoforms of protein-coding genes containing functional domains (**Fig. 4** and **Fig. 5i-j**), we reasoned that some lncRNAs undergoing IS may also harbor important functions that need to be regulated during the first half of the reprogramming process and thus for pluripotency acquisition.

### Identification of a non-coding transcript regulating the pluripotent state

Because isoform switching of lncRNAs is under strict control during early reprogramming (**Fig. 5d**), we hypothesized that isoform switching could regulate several lncRNAs involved in the pluripotency regulatory network. To increase chances of identifying functionally relevant IS events in lncRNAs, we focused on lncRNAs undergoing isoform switching during NPCs to mid-reprogramming in either OKMS or OKS systems and that are identified in both our ΔPSI events-based analysis (n=1,664 lncRNA genes) and our DEIs-based analysis of (n=58 lncRNA genes) (**Fig. 6a**). We obtained 45 lncRNA candidates that were common to these two lists (**Fig. 6a** and **Supplementary Data 11**). In order to validate the importance of these candidates for reprogramming, we cross-referenced our list with lncRNAs that were reproducibly detected and differentially expressed in at least one other reprogramming system (i.e. OSKM or OKMS with 2° MEFs as a starting cell type) and in ESCs^16,25,42^. We identified ten candidates (**Fig. 6b-c**, **Supplementary Fig. 10a-b**, and **Supplementary Data 11**). From these, *2700038G22Rik*, *2010204K13Rik*, *Gm11944*, and *Mir17hg* were the only four lncRNAs with expression increasing throughout both reprogramming datasets and remained consistently higher in iPSCs than in MEFs (**Fig. 6b-c** and **Supplementary Fig. 10a-b**). More specifically, the lncRNA *2700038G22Rik* showed this pattern consistently across multiple reprogramming datasets (**Fig. 6c**) suggesting a possible role in the acquisition of the pluripotent state. Moreover, a human syntenic ortholog could only be found for three of the four candidates: *2700038G22Rik, 2010204K13Rik* and *Mir17hg* lncRNAs (**Fig. 6e**, **Supplementary Fig. 10c**, and **Supplementary Data 11**). However, the human ortholog for *2010204K13Rik* lncRNA appeared to be the coding gene *PAGE4*, which was only expressed in late reprogramming (**Supplementary Fig. 10c**). For this reason, *2010204K13Rik* was not studied further. As for *Mir17hg*, it is a conserved precursor for the *miR-17-92* microRNAs cluster^74,75^, which have been shown to be highly expressed in ESCs to regulate cell cycle progression^76^ and expression of pluripotency genes^77,78^. These miRNAs are also induced during the early phase of reprogramming^78,79^ and enhance human fibroblast OKMS and OKS reprogramming^79,80^ by decreasing TGFBR2 and p21 protein levels to overcome reprogramming barriers^80^. Identification of *Mir17hg* as a lncRNA whose by-products are well studied in ESCs and reprogramming confirmed that our identification pipeline is thus appropriate to discover lncRNAs with a function in pluripotency regulation, and suggested that the other top candidate *2700038G22Rik* could be important as well.

**Figure 6:**
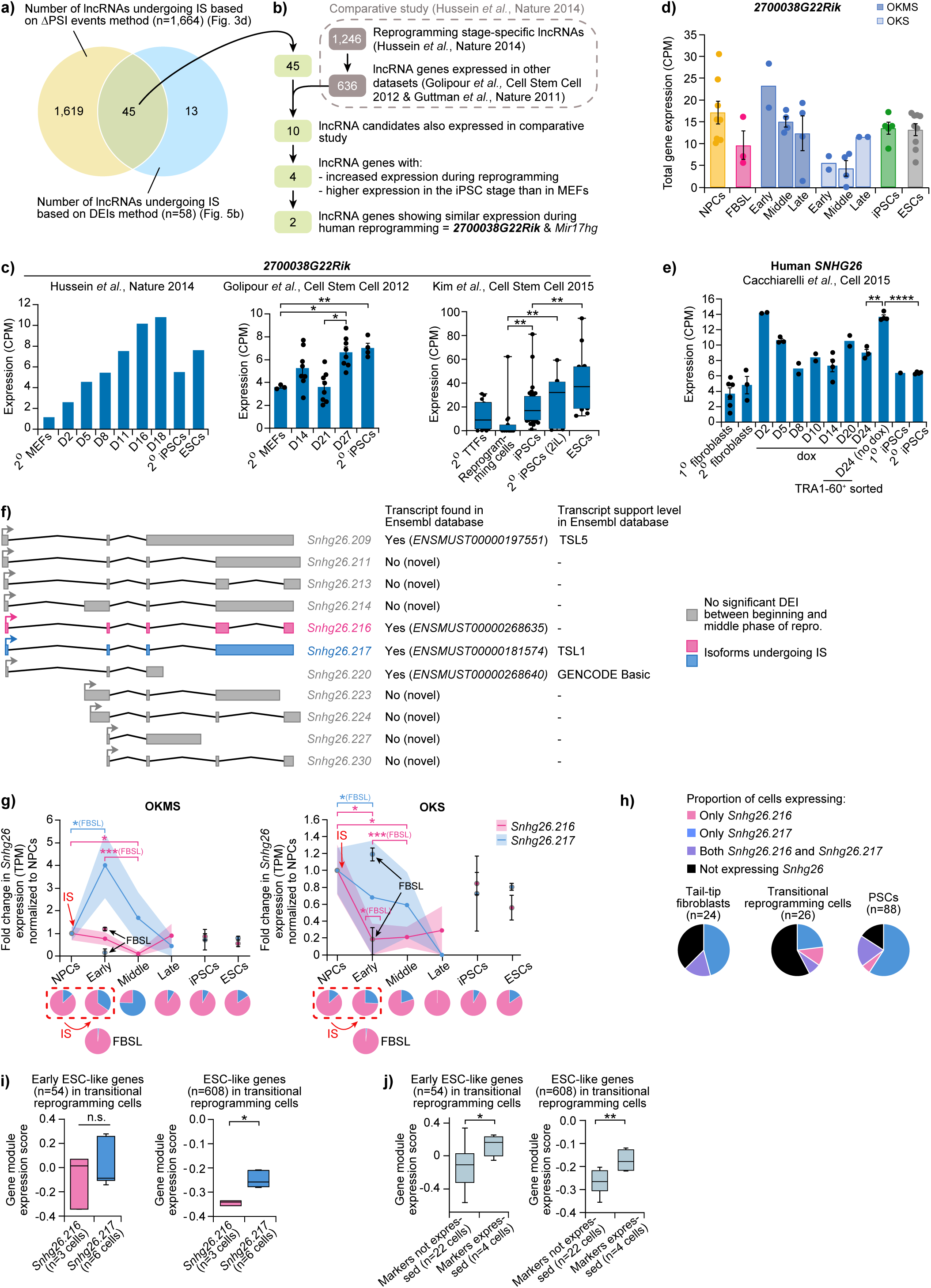
Identification of a candidate non-coding transcript regulating the pluripotent state. **a)** Overlap between the number of lncRNA genes which undergo isoform switch between D0 and the middle phase of our reprogramming systems, based on two methods. **b)** Pipeline used to isolate lncRNA candidate genes potentially involved in the regulation of pluripotency induction. **c)** Average expression of *2700038G22Rik* lncRNA at different timepoints of two 2° MEFs reprogramming RNAseq datasets^16,42^ (Golipour *et al.*, Cell Stem Cell 2012: 2° MEFs (n=3), D14 (n=8), D21 (n=8), D27 (n=8), 2° iPSCs (n=4). Error bars represent the SEM and each dot represents replicates.) and one 2° TTFs reprogramming single-cell RNAseq dataset^18^ (Kim *et al.*, Cell Stem Cell 2015: 2° TTFs (n=14), reprogramming cells (n=15), iPSCs (n=31), 2° iPSCs (n=8), ESCs (n=14). Central lines represent medians and boxes denote quartiles. Error bars represent the minimum and maximum values. Dots represent outlier cells). P-values were calculated using two-sided t-test with unequal variance. * < 0.05, ** < 0.01. TTFs: Tail-Tip Fibroblasts. **d)** Mean expression of *2700038G22Rik* gene in NPCs (n=8), FBSL (n=3), during early (n=2), middle (n=4) or late (n=4 for OKMS and n=2 for OKS) phases of reprogramming, or in the iPSCs (n=5) and ESCs (n=9) of the dataset. Error bars represent the SEM and each dot represents replicates. **e)** Average expression of *SNHG26* lncRNA at different timepoints of a 2° human fibroblasts reprogramming RNAseq dataset^81^ (1° fibroblasts n=6, 2° fibroblasts n=3, D2 n=2, D5 n=3, D8 n=2, D10 n=2, D14 n=4, D20 n=2, D24 n=3, D24 no dox n=4, 1° iPSCs n=1, 2° iPSCs n=4). Error bars represent the SEM and each dot represents replicates. P-values were calculated using two-sided t-test with unequal variance. ** < 0.01, **** < 0.0001. **f)** Schematic depicting the transcripts of the *Snhg26* gene detected in our dataset and their level of support on the latest version of Ensembl database (release 113)^82^. TSL: Transcript Support Level. **g)** Expression fold change normalized to NPCs, of two *Snhg26* lncRNA isoforms in NPCs (n=8), FBSL (n=3), during early (n=2), middle (n=4) or late (n=4 for OKMS and n=2 for OKS) phases of reprogramming, or in the iPSCs (n=5) and ESCs (n=9) of the dataset. Dots and lines represent average expression, and shaded areas and error bars represent % change. Pie charts represent the proportion of each transcript. P-values were calculated using one-sided t-test with unequal variance. * < 0.05, *** < 0.001. The data is the same in OKMS and OKS for NPCs, FBSL, iPSCs and ESCs samples because those samples were in common for OKMS and OKS (grey circles). TPM: Transcripts Per Million reads. **h)** Proportions of cells expressing *Snhg26* isoforms in a 2° TTFs reprogramming single-cell RNAseq dataset^18^. **i)** Box plots of Seurat gene module normalized expression scores for early and ESC-like genes (gene lists from Hussein *et al.*^16^) between cells expressing only isoform *Snhg26.216* or *Snhg26.217*, or **j)** cells based on expression of coding isoforms from at least two of three markers of proper reprogramming (*Dppa2, Utf1,* and *Esrrb*). Bounds of the box represent 25^th^ and 75^th^ quartiles, bounds of the whiskers represent minimum and maximum values, and center line represents median. P-values were calculated by a two-sided Mann-Whitney non-parametric U-test. n.s. > 0.05, * = 0.024 (ESC-like genes), 0.048 (early ESC-like genes), ** = 0.0071 (ESC-like genes).

Interestingly, NPCs also express *2700038G22Rik* (**Fig. 6d**), and its expression increased even further during OKMS reprogramming but decreased and then increased in the OKS system to reach iPSCs/ESCs levels (**Fig. 6d**). This increase in expression was reproducible in different datasets using 2° MEFs or 2° tail-tip fibroblasts, with expression levels in iPSCs comparable to those in control ESCs^16,18,42^ (**Fig. 6c**). Moreover, expression of the human ortholog of *2700038G22Rik*, *SNHG26*, showed a similar expression pattern of up-regulation during human reprogramming^81^ (**Fig. 6e**). *2700038G22Rik* lncRNA thus appeared as a promising regulator of pluripotency, as its expression is increasing during reprogramming, it is highly expressed in pluripotent stem cells, and has a conserved expression pattern in different mouse and human reprogramming systems.

Because *2700038G22Rik* (named *Snhg26* from here on) was identified as undergoing IS, we analyzed its different isoforms and identified 11 transcripts expressed among the 276,608 isoforms detected in our dataset (**Fig. 6f**). Only four could be identified in the latest Ensembl database version (release version 113.39, Aug. 2024)^82^ (**Fig. 6f**), highlighting dataset specific isoform diversity and the importance of long-read data integration into public databases. In our reprogramming dataset, two isoforms showed differential expression between the early and middle phases. Those two isoforms, *G38722.216* (“*Snhg26.216*”) and *G38722.217* (“*Snhg26.217*”), demonstrated an opposite expression pattern: *Snhg26.216* level was more elevated than the one of *Snhg26.217* in NPCs (**Supplementary Fig. 10d-e**), but decreased significantly in the middle phase of both OKMS and OKS reprogramming (**Fig. 6g** and **Supplementary Fig. 10d-e**). *Snhg26.217* expression increased drastically in OKMS reprogramming early phase (relative to NPCs) so that in middle phase, the proportion of *Snhg26.217* was significantly more (going from 13.2% in NPCs to 75.5% in middle phase) (**Fig. 6g**, **Supplementary Fig. 10d-e,** and **Supplementary Data 11**), thus leading to a clear isoform switching event. In OKS reprogramming, both *Snhg26.217* and *Snhg26.216* expression decreased, but the fraction of *Snhg26.217* isoform increased in early reprogramming (going from 13.2% in NPCs to 25.7% in early phase) at the expense of the *Snhg26.216* isoform (**Fig. 6g** and **Supplementary Data 11**). The strong increase of *Snhg26.217* over *Snhg26.216* isoform was further validated by RT-PCR followed by lrRNAseq (**Supplementary Fig. 11a** and **Supplementary Data 12**). Similarly, in OSKM reprogramming from B cells^73^, opposed expression dynamics were observed for *Snhg26.217* and *Snhg26.216* (**Supplementary Fig. 11b**). Early reprogramming from B cells is associated with an increase in the proportion of *Snhg26.216* isoforms, at the expense of *Snhg26.217* expression. Later in reprogramming, *Snhg26.217* levels rise back to their original levels, suggesting that *Snhg26* isoform switching is conserved across multiple reprogramming systems.

To assess whether *Snhg26* isoforms are associated with reprogramming outcome, we analyzed single-cell RNA sequencing data from reprogramming cells^18^ (**Supplementary Fig. 11c** and **Supplementary Data 10**). During reprogramming (i.e. “transitional reprogramming cells”), 23% of cells express only *Snhg26.217*, 11% express only *Snhg26.216*, and 7% express both, and the proportion of *Snhg26.217* expressing cells increased significantly when cells reach the pluripotent state (**Fig. 6h**). Moreover, transitional cells expressing *Snhg26.217* isoform have significantly higher expression of ESC-like genes when compared to cells expressing *Snhg26.216* (**Fig. 6i** and **Supplementary Fig. 11d**). This mirrors the subset of transitional reprogramming cells expressing known markers associated with favorable reprogramming outcome (e.g. *Dppa2*, *Utf1* and *Esrrb*^49^) (**Fig. 6j** and **Supplementary Fig. 11e-f**). This data, as well as *Snhg26* conserved expression pattern among different reprogramming systems and species, suggests that this lncRNA, more specifically the *Snhg26.217* isoform, may function as an important regulator of the early stages of pluripotency induction.

### *Snhg26.217* lncRNA enhances the reprogramming route to pluripotency

In order to investigate if *Snhg26*, more specifically isoform *217*, could have a function during the acquisition of mouse pluripotency, we performed reprogramming experiments where we overexpressed either a control RNA or the *Snhg26.217* lncRNA isoform with the reprogramming factors OKS (**Supplementary Data 11**). Upon DOX addition, the proportion of reprogramming mCherry+ cells peaked earlier for OKS + *Snhg26.217* (80% of cells at D8) than for OKS + control (58.35% of cells at D14) (**Fig. 7a**, **Supplementary Fig. 1**, and **Supplementary Fig. 12a-b**). However, activation of the *Oct4* locus (i.e. GFP+ cells) was similar starting from D14 (2.12% for OKS + control; 1.44% for OKS + *Snhg26.217*) and increasing similarly between the conditions to reach their maximum at D20 (21.78% for OKS + control; 18.50% for OKS + *Snhg26.217*) (**Fig. 7b**). By this time, most cells had spontaneously silenced transgenes expression (11.05% of mCherry+ cells for OKS + control; 13.39% for OKS + *Snhg26.217* at D20) (**Fig. 7a**). Surprisingly, the increased number of mCherry+ cells when *Snhg26.217* was expressed did not lead to significantly more GFP+ cells upon DOX removal (63.35% for OKS + control; 75.55% for OKS + *Snhg26.217* at D14) (**Fig. 7b**), even if mCherry levels remained high (going from 51.61% at D6 to 73.4% at D10 for OKS + *Snhg26.217,* compared to 37.70% at D10 for OKS + control) (**Fig. 7a**). For both conditions, most cells silenced the exogenous reprogramming factors upon DOX removal, starting by D14 (5.72% for OKS + control; 7.07% for OKS + *Snhg26.217*) (**Fig. 7a**). The fact that the proportion of mCherry+ cells for OKS + *Snhg26.217* was more than twice the proportion of OKS + control reprogramming cells, and that this was not reflected in the percentage of GFP+ cells emerging, suggested two scenarios which could explain how *Snhg26.217* expression increased the number of reprogramming cells: 1) *Snhg26.217* could directly affect proliferation of cells that commit to reprogramming but not the establishment of iPSCs (e.g. cells are not able to complete reprogramming to the final iPSC stage, possibly remaining blocked at the pre-iPSC stage), or 2) *Snhg26.217* may influence the increase in iPSC colony formation (i.e. more cells commit to iPSCs) (**Fig. 7c**). For the latter scenario, the total number of cells would be greater under *Snhg26.217* expression because of an increase in reprogramming cells (i.e. mCherry+ cells); thus, looking at percentages will not reflect this increase as global cell confluency is not considered in such calculations. For example, an increase in mCherry+ cells leads to an increase in the number of cells that commit to reprogramming; however, for cells to become iPSCs (i.e. GFP+ cells) and for reprogramming to complete, transgene expression must be silenced. This will lead to a mixed population of GFP+ and negative cells, which will show an increase in cell density but not necessarily the proportion of iPSCs (please refer to **Supplementary Note Fig. 2** for details).

**Figure 7:**
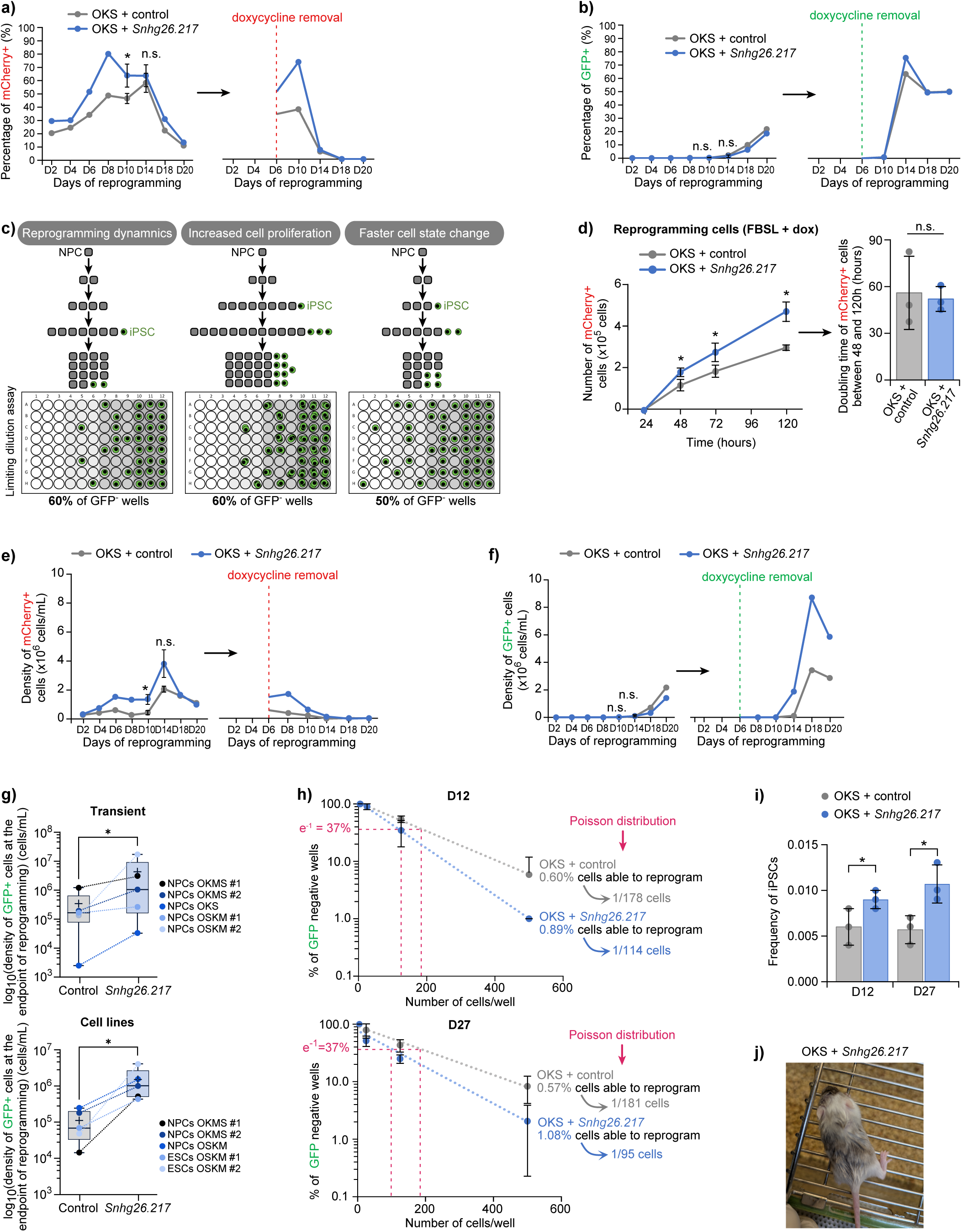
*Snhg26.217* isoform enhances the reprogramming route to pluripotency. **a)** Percentages of mCherry+ or **b)** GFP+ cells in FBSL + doxycycline medium, or after doxycycline removal in 2i-LIF medium, during reprogramming with OKS + control or OKS + *Snhg26.217* transgenes. Error bars represent SEM (n=3). P-values were calculated using one-sided t-test with paired samples to specifically ask if *Snhg26.217* enhances reprogramming. n.s. > 0.05, * = 0.029. The data is the same as Supplementary Note Fig. 1b-c for OKS + control because experiments were conducted at the same time for OKMS, OKS and OKS + *Snhg26.217* lncRNA. **c)** Schematic representing the possible effect of *Snhg26.217* expression on the dynamics of somatic cell reprogramming and on the results of a limiting dilution assay, depending on whether it increases cell proliferation or the kinetics of cells to reprogram. **d)** Average number of mCherry+ cells and their mean doubling times from the OKS + control (n=3 biological replicates) and OKS + *Snhg26.217* (n=3 biological replicates) reprogramming cell lines through the first 5 days of reprogramming. Error bars represent the SEM. P-values were calculated using one-sided t-test on paired samples to specifically ask if *Snhg26.217* increases doubling time of mCherry+ cells. * = 0.029 (48h), 0.014 (72h), 0.026 (120h). For the bar graph of mean cell doubling times, error bars represent the STD. P-values were calculated using two-sided t-test with unequal variance. n.s. ≥ 0.05. **e)** Average density of mCherry+ or **f)** GFP+ cells in FBSL + doxycycline medium, or after doxycycline removal in 2i-LIF medium, during reprogramming with OKS + control or OKS + *Snhg26.217* transgenes. Error bars represent the SEM (n=3 biological replicates). P-values were calculated using one-sided t-test with paired samples to specifically ask if *Snhg26.217* increases density of mCherry+ or GFP+ cells. n.s. > 0.05, * = 0.029. **g)** Logged density of GFP+ cells at the endpoint of reprogramming, using either transient transfections (n=5 biological replicates) or stable cell lines (n=5 biological replicates) to express the control or *Snhg26.217* transgenes. Horizontal lines represent medians, crosses represent averages, and boxes denote quartiles. Error bars represent the minimum and maximum values. Dots represent replicates. P-values were calculated by one-tailed Wilcoxon matched-pairs signed-rank test to specifically ask if *Snhg26.217* increases density of GFP+ cells. * = 0.031. The data is the same as each endpoint presented in Supplementary Fig. 13a-f. **h)** Mean percentages of GFP-negative wells for the different cell densities FBSL + doxycycline medium (D12) and in 2i-LIF medium (D27) (n=3), allowing the use of Poisson distribution. Error bars represent the SEM. **i)** Average iPSC frequencies in OKS + control (n=3 biological replicates) and OKS + *Snhg26.217* (n=3 biological replicates) conditions at D12 and D27 of reprogramming. Error bars represent the STD and each dot represents replicates. P-values were calculated using one-sided t-test with unequal variance to specifically ask if *Snhg26.217* increases frequency of iPSCs. * = 0.044 (D12), 0.019 (OKS). **j)** Representative image of a chimeric mouse obtained from aggregation of OKS + *Snhg26.217* iPS clones with 8-cell CD-1 albino embryos for pluripotency assessment.

To test these two scenarios, we first addressed whether *Snhg26.217* expression could increase the proliferation of reprogramming cells. We found that *Snhg26.217* expression significantly and consistently increased the population of mCherry+ cells (297,253 cells for OKS + control; 471,693 cells for OKS + *Snhg26.217* at 120 hours post-DOX induction) (**Fig. 7d**) and the total number of cells (**Supplementary Fig. 12c**). However, no significant difference was observed in mean doubling time of these mCherry+ or total populations between 24h and 120h of DOX induction between control and *Snhg26.217* conditions (**Fig. 7d** and **Supplementary Fig. 12c**), suggesting that *Snhg26.217* expression does not influence cell proliferation. We also quantified the doubling times of the starting NPC lines following the expression of a control or *Snhg26.217* RNA, in NPC medium and independent of reprogramming medium conditions. Once again, their average proliferation rates were similar (**Supplementary Fig. 12d**), suggesting that *Snhg26.217* overexpression did not impact the growth of the starting NPC lines. Finally, to make sure that *Snhg26.217* expression does not influence the division rate of pluripotent cells themselves, we compared the doubling times of ESCs overexpressing *Snhg26.217* or a control RNA in 2i-LIF + DOX medium every 12 hours. Again, there was no difference in the proliferation curves of ESCs when comparing control to *Snhg26.217* overexpression (**Supplementary Fig. 12e**). These results demonstrated that *Snhg26.217* did not affect cell proliferation and that scenario 2 (i.e. increase in iPSC colony formation) was more plausible.

To address the second scenario, and since the increased number of reprogramming cells led to a global cell population increase, it was thus important to account for the global density of cells in the calculation of mCherry+ and GFP+ cells during reprogramming to properly measure the *Snhg26.217* effect on the process (please refer to **Supplementary Note Fig. 2** for details). By doing this, we observed that the mean density of mCherry+ cells progressively increased to reach its maximum at D14 for both OKS + control (2,065,072 cells/mL) and OKS + *Snhg26.217* (3,823,884 cells/mL) conditions and subsequently decreased to give rise to GFP+ cells (**Fig. 7e-f**). Upon DOX removal, transgene expression dropped rapidly between D10 and D14 for both conditions, and we observed a strong induction of iPSCs at D14, with 16 times more GFP+ cells than in control reprogramming (112,328 cells/mL for OKS + control; 1,877,379 cells/mL for OKS + *Snhg26.217*) (**Fig. 7f**). These results highlight that scenario 2 is the most plausible explanation for our observations and that *Snhg26.217* may accelerate reprogramming to iPSCs. Therefore, by analyzing cell density, it allowed us to visualize that the number of both reprogramming cells and iPSCs were enhanced by *Snhg26.217* overexpression compared to the control condition. To validate the robustness of this result, we analyzed the density of GFP+ cells using other reprogramming strategies. We found that the density of GFP+ cells was always higher when *Snhg26.217* was expressed compared to the control condition regardless of reprogramming factors order, cell origin, or whether it was done using transient or stable transfections (**Fig. 7g** and **Supplementary Fig. 13a-f**). Together, these results validated that *Snhg26.217* significantly enhanced the induction of iPSCs in various reprogramming systems.

We next asked if *Snhg26.217* could influence the frequency of cells able to engage into reprogramming and to reach the iPSC stage by changing cell identity and enhancing the cells’ rate of reprogramming. To assess this, we performed a limiting dilution assay, a robust assay usually used to assess the frequency of stem cells in a cell population *in vitro* (please refer to **Supplementary Note Fig. 3** for details)^83–85^. We used the same OKS reprogramming system as before, but this time we seeded the starting cell lines as graded dilutions (5, 25, 125 or 500 cells/well) with 24 replicates in a culture plate. We induced the transgenes with doxycycline and calculated the percentage of wells negative of any reprogramming events at D12 and then at D27 after removing DOX at D18. We found that in the control condition, about 0.6% of cells were able to give rise to iPSCs during OKS reprogramming (0.60% (1/178 cells) at D12; 0.57% (1/181 cell) at D27) (**Fig. 7h-i**). However, when we overexpressed *Snhg26.217* along with the reprogramming factors, this frequency increased to about 1% (0.89% (1/114 cells) at D12; 1.08% (1/95 cells) at D27) (**Fig. 7h-i**). Together, these results led us to conclude that *Snhg26.217* accelerates the reprogramming kinetics by enhancing the overall efficiency and increasing cell state changes rather than increasing cell division (for a more detailed explanation, see **Supplementary Note Fig. 2**).

To validate that iPSCs generated with *Snhg26.217* are *bona fide* iPSCs, we isolated clones from the OKS + *Snhg26.217* reprogramming experiments and tested their contribution to the formation of chimeras using aggregations. E8-8.5 embryos showed high contribution for OKS *+ Snhg26.217* iPSCs, as visualized with the negative cells/area in tdTomato-positive embryos (**Supplementary Fig. 13g**). As expected, iPSC contribution was also limited to the embryonic area (i.e. epiblast derivatives), with extraembryonic tissues remaining tdTomato-positive. When aggregated to albino mouse embryos, the iPSC clones contributed 5-55% to chimera formation, as judged by coat color (**Fig. 7j** and **Supplementary Fig. 13h**). Finally, OKS + *Snhg26.217* iPSCs also successfully formed teratoma when injected into mice, and cells derived from the three germ layers were represented to the same extent as OKMS/OKS + control iPSCs (**Supplementary Fig. 14**). This confirmed that *Snhg26.217* expression during reprogramming gave rise to *bona fide* iPSCs.

### *Snhg26* lncRNA is a developmental gene important for the pluripotent state

To further investigate the regulation and function of *Snhg26* lncRNA in pluripotency, we used our previous ChIPseq during reprogramming of 2°MEFs to iPSCs^16^ and analyzed the chromatin landscape around *Snhg26* locus. The *Snhg26* promoter was marked by H3K4me3 and H3K27me3 and devoid of DNA methylation in 2°MEFs, but lost the H3K27me3 mark as cells progressed through reprogramming towards iPSCs (**Fig. 8a**), suggesting a bivalent locus that is activated upon loss of the repressive mark H3K27me3. This is in accordance with up-regulation of *Snhg26* during reprogramming and a sign of a developmentally regulated genes^86,87^. Moreover, a previous study identified *Snhg26* as part of a cohort of ESC-specific lncRNAs and showed that knock-downs of KLF2 or N-MYC pluripotency-associated transcription factors, both lead to a down-regulation of *Snhg26* (named *linc1533* in that study) in mouse ESCs^25^. They also showed the binding of KLF4, C-MYC and N-MYC to *Snhg26* promoter^25^, suggesting that *Snhg26* may be regulated by the pluripotency-transcriptional network.

**Figure 8:**
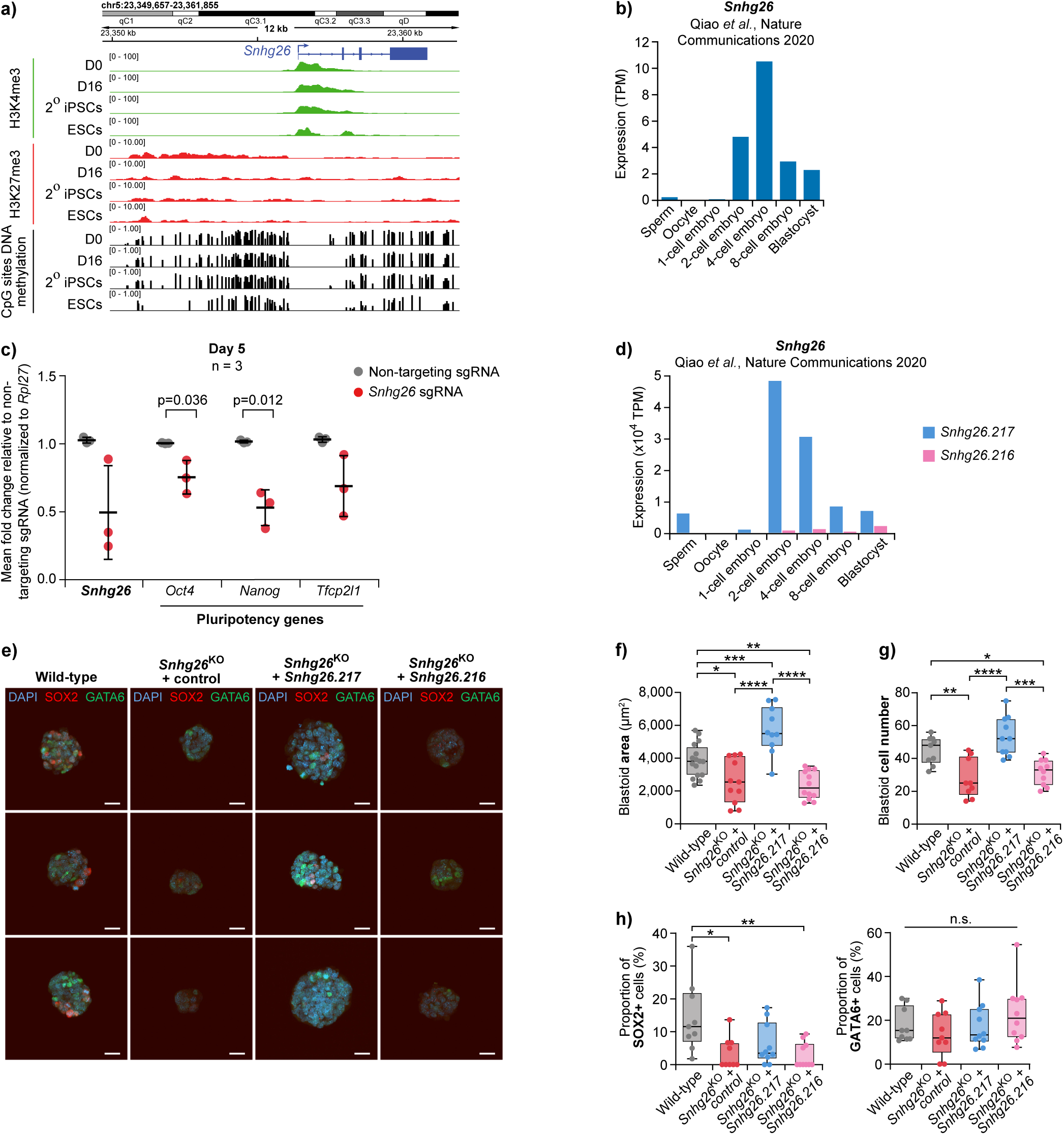
*Snhg26* is a developmental gene important for mouse pluripotency. **a)** Signal from ChIPseq data^16^ at the genomic locus of *Snhg26*. Tracks represent 2° MEFs (D0) and their reprogrammed counterparts after 16 days of reprogramming (D16), 2° iPSCs and control ESCs. **b)** Gene-level expression of *Snhg26* lncRNA in different stages of a mouse pre-implantation embryo dataset^13^. **c)** Mean fold changes in expression for *Snhg26* lncRNA and pluripotency-associated genes, assessed by RT-qPCR after *Snhg26* knock-down in mouse ESCs (n=3) using the CRISPRi system. Expression is normalized to a control sample with non-targeting single-guide RNA (sgRNA) and to a housekeeping gene. Horizontal line represents the average, error bars represent the STD and each dot represents replicates. P-values were calculated using one-sided t-test with unequal variance to specifically ask if *Snhg26* knock-down decreases pluripotency genes expression. p: p-value. **d)** Expression of *Snhg26* isoforms in different stages of a mouse pre-implantation embryo dataset^13^. **e)** Representative images in fluorescence microscopy of wild-type or *Snhg26*^KO^ blastoids obtained after 3 days of overexpression of either a control transcript or *Snhg26* isoforms. Scalebars = 20 µm. KO: Knock-Out. **f)** Area of the wild-type (n=16), *Snhg26*^KO^ + control (n=11), *Snhg26*^KO^ + *Snhg26.217* (n=10), or *Snhg26*^KO^ + *Snhg26.216* (n=12) blastoids. Central lines represent medians and boxes denote quartiles. Error bars represent the minimum and maximum values. Dots represent replicates. P-values were calculated by ordinary one-way ANOVA with Tukey HSD *post hoc* test. * < 0.05, ** < 0.01, *** < 0.001, **** < 0.0001. **g)** Total number of cells, or **h)** proportion of cells positive for SOX2 or GATA6 staining in the wild-type (n=9), *Snhg26*^KO^ + control (n=9), *Snhg26*^KO^ + *Snhg26.217* (n=10), or *Snhg26*^KO^ + *Snhg26.216* (n=10) blastoids. Central lines represent medians and boxes denote quartiles. Error bars represent the minimum and maximum values. Dots represent replicates. P-values were calculated by ordinary one-way ANOVA with Tukey HSD *post hoc* test. n.s. ≥ 0.05, * < 0.05, ** < 0.01, *** < 0.001, **** < 0.0001.

Because those results suggested a role for *Snhg26* in ESCs, we wondered if *Snhg26* expression could be conserved in the broader context of pluripotency and development. To this end, we evaluated total *Snhg26* expression during mouse early embryo development using a published dataset^13^. Interestingly, *Snhg26* expression peaks at the 4-cell stage of pre-implantation embryos, and remains expressed up to the blastocyst stage (i.e. pluripotent state), albeit at a lower expression level (**Fig. 8b**). This peak expression corresponds to the zygotic genome activation stages in mice and suggests a role for *Snhg26* in embryo development. Furthermore, if we examine *Snhg26* expression during *in vitro* differentiation of ESCs towards cortical neurons^88^, we observe that it drops drastically from the first days of the differentiation process (**Supplementary Fig. 15a**). These results suggest that expression of *Snhg26* is specific to early embryo development and the associated pluripotent state.

To address its role in pluripotency, we first sought to determine where *Snhg26* lncRNA is localized as localization of a lncRNA is generally indicative of its function^89,90^. We found *Snhg26* to be nuclear (83.8% on average) according to quantifications performed after cellular fractionation of mouse ESCs (**Supplementary Fig. 15b** and **Supplementary Data 13**). Its expression was similar to other well-characterized nuclear lncRNAs such as *Xist* (88.2% on average), *Malat1* (85.4% on average) and *Neat1* (89.5% on average). Moreover, *in situ* hybridization of probes targeting *Snhg26* transcripts demonstrated strong co-localization with DAPI staining (**Supplementary Fig. 15c**), further validating *Snhg26* nuclear localization in mouse ESCs. Therefore, we knocked-down *Snhg26*’s transcription in mouse ESCs by targeting its promoter with a CRISPR-interference (CRISPRi) system^91,92^, which also ensures that all its transcripts were targeted. Knock-down of *Snhg26*’s transcription for 5 days in mouse ESCs led to down-regulation of pluripotency genes *Oct4, Nanog* and *Tfcp2l1* (**Fig. 8c**), confirming that *Snhg26* was involved in the pluripotency regulatory network of ESCs.

Finally, in order to link the role of the specific *Snhg26.217* isoform in enhancing reprogramming with the function of the *Snhg26* gene in pluripotency, we asked whether *Snhg26* isoforms had similar or divergent roles during early embryonic development. First, we assessed their expression in a mouse embryo published dataset^13^ and found that while *Snhg26.216* was expressed between the 2-cell and blastocyst stages, *Snhg26.217* expression levels were always higher, particularly high at the 2-cell and 4-cell stages (**Fig. 8d** and **Supplementary Data 13**). This suggested that *Snhg26.217* may have a more important role in early development than *Snhg26.216*. To assess this, we generated a *Snhg26* knock-out (KO) mouse ESC line by removing the last exon of *Snhg26*, which represents 92.5% of its whole exonic sequence using CRISPR-Cas9. We were able to generate an *Snhg26*^KO^ ESC clone, that was then engineered to overexpress either *Snhg26.217* or *Snhg26.216* transcripts using DOX induction. Three-dimension blastoid models, *in vitro* models of pre-implantation blastocysts, were derived from the KO + *Snhg26* transcript lines, along with wild-type cells, over three days of DOX induction and stained for markers of the inner cell mass (SOX2) and the primitive endoderm (GATA6). We found that *Snhg26* KO led to smaller blastoids (**Fig. 8e-f** and **Supplementary Fig. 15d**), with a significantly reduced number of cells (**Fig. 8g**) compared to wild-type ones. Moreover, they displayed less SOX2-positive cells and an unchanged number of GATA6-positive cells, suggesting that lack of *Snhg26* lncRNA prevented the formation of inner cell mass-like cells (**Fig. 8h**). Strikingly, overexpression of *Snhg26.217* isoform rescued the *Snhg26*^KO^ blastoid size and total number of cells, even increasing those features (**Fig. 8f-g** and **Supplementary Fig. 15d**). This time, the number of SOX2 and GATA6-positive cells was not significantly affected (**Fig. 8e-h**), suggesting that *Snhg26.217* isoform rescues development of inner cell mass-like cells. On the other hand, *Snhg26*^KO^ blastoids overexpressing *Snhg26.216* isoform displayed a phenotype similar to the *Snhg26*^KO^ blastoids (**Fig. 8e-h** and **Supplementary Fig. 15d**). These results suggest that *Snhg26.217* isoform expression, and not that of *Snhg26.216,* is important for proper embryo development and formation of the inner cell mass of mouse early blastocysts.

Altogether, the effects of overexpression, knock-down, and knock-out of *Snhg26* and its isoforms in pluripotency models such as reprogramming, ESCs and blastoids demonstrate the role of *Snhg26* lncRNA, and more specifically of *Snhg26.217* isoform, in the instauration of the pluripotent state. Thus, lrRNAseq analysis of our reprogramming systems enabled us to identify a functional isoform of a new lncRNA involved in regulating the pluripotent state.

### *SNHG26* lncRNA has a conserved function in human pluripotency

Because *Snhg26* was selected from a lncRNA candidate list for its conserved expression pattern during mouse and human reprogramming, we wondered if it could have a similar function in the regulation of human pluripotency. To this end, we assessed the expression of its human counterpart*, SNHG26* lncRNA, during human embryo development using a published dataset^93^. Similarly to the mouse lncRNA, *SNHG26* expression started at the 8-cell stage of pre-implantation embryos, which corresponds to the zygotic genome activation stages in humans (**Fig. 9a**). *SNHG26* remained quite highly expressed in human pluripotent stem cells. Because mouse and human ESCs (hESCs) correspond to different states of pluripotency^94,95^, we examined *SNHG26* expression in primed and *naïve* hESCs, as well as *in vitro* equivalent of 8 cell-like cells^96^. We found that *SNHG26* was highly expressed in *naïve* and 8 cell-like cells compared to primed hESCs (**Fig. 9b**), which is in keeping with results from pre-implantation embryo datasets. Finally, exploring *SNHG26* expression during *in vitro* differentiation of hESCs towards cortical neurons^97^, we observed that its expression dropped drastically from the beginning of the process (**Supplementary Fig. 16a**). These expression patterns suggest that *Snhg26* may have a specific and conserved role in early embryo development and the associated pluripotent state.

**Figure 9:**
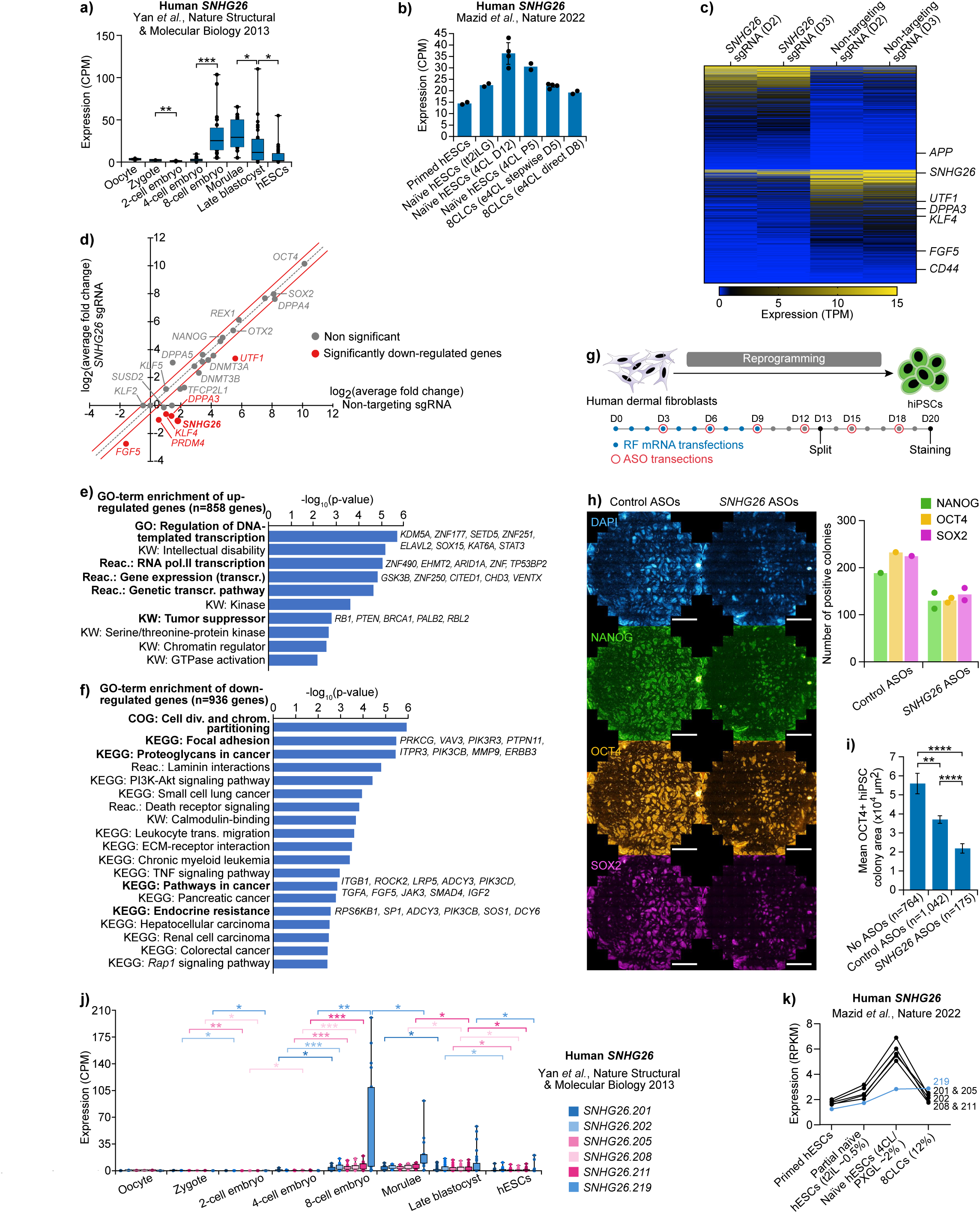
*Snhg26* is a developmental gene important for human pluripotency. **a)** Average expression of *SNHG26* lncRNA in different stages (oocyte n=3; zygote n=3; 2-cell embryo n=6; 4-cell embryo n=12; 8-cell embryo n=20; morulae n=16; late blastocyst n=30; hESCs n=34) of a human pre-implantation embryo dataset^93^. Central lines represent medians and boxes denote quartiles. Error bars represent the minimum and maximum values. Dots represent outlier cells. P-values were calculated using two-sided t-test with unequal variance. * < 0.05, ** < 0.01, *** < 0.001. hESCs: human Embryonic Stem Cells. **b)** Expression of *SNHG26* lncRNA in primed (n=2) and *naïve* hESCs (tt2iLG n=2; 4CL D12 n=4; 4CL P5 n=2) or 8CLCs (D5 n=4; D8 n=2) samples from an available dataset^96^. Error bars represent the SEM and each dot represents replicates. tt2iLG: titrated 2 inhibitors with LIF and Gö6983 medium; e4CL: enhanced 4 chemical with LIF medium; 8CLCs: 8 Cell stage embryo-Like Cells. **c)** Heat map depicting the expression values of genes significantly up- or down-regulated (n=1,794) at D2 and D3 of *SNHG26* knock-down in hESCs, compared to control hESCs with a non-targeting sgRNA. Significance based on a Shannon entropy score below tenth percentile. Maximum expression value was set at ≥ 15 TPM. **d)** Logged expression of pluripotency genes (n=29) (significantly decreased are shown in red) following *SNHG26* knock-down in hESCs, as compared to conditions with a non-targeting sgRNA. Significance based either on fold change ≤ 0.67 and adjusted p-value ≤ 0.05 (calculated by Benjamini-Hochberg method), or on a Shannon entropy score below tenth percentile. **e)** Logged p-values from GO-term enrichment analysis for genes up-regulated or **f)** down-regulated following *SNHG26* knock-down in hESCs. GO-term significance based on fold enrichment ≥ 1.5, p-value ≤ 0.05 and FDR ≤ 0.1. P-values were calculated by one-sided Fisher’s exact test. KW: Key Words; Reac: Reactome pathway; pol.: polymerase; transcr.: transcription; COG: Clusters of Orthologous Genes; div.: division; chrom.: chromosome; trans.: transendothelial. **g)** Schematic of *SNHG26* knock-down during human reprogramming. hiPSCs: human induced Pluripotent Stem Cells; RF: Reprogramming Factors; ASO: AntiSense Oligonucleotide. **h)** Representative whole-well immunofluorescence images and quantification of iPSC colonies stained with DAPI, NANOG, OCT4, and SOX2 antibodies at D20 of human reprogramming with control (n=1) or *SNHG26-*targeting (n=2) antisense oligonucleotides transfection. Bars represent the average and dots represent biological replicates. Scalebars = 3,000 µm. **i)** Mean area of OCT4-positive iPSC colonies at D20 of human reprogramming with control (n=764 no ASOs; n=1,042 control ASOs) or *SNHG26-*targeting ASOs (n=175). Error bars represent the SEM. P-values were calculated using two-sided t-test with unequal variance. ** = 0.001, **** = 9.2x10^-9^ (no ASOs to *SNHG26* ASOs), 2.3x10^-6^ (control ASOs to *SNHG26* ASOs). **j)** Expression of *SNHG26* isoforms in different stages (oocyte n=3; zygote n=3; 2-cell embryo n=6; 4-cell embryo n=12; 8-cell embryo n=20; morulae n=16; late blastocyst n=30; hESCs n=26) of a human pre-implantation embryo dataset^93^. Central lines represent medians and boxes denote quartiles. Error bars represent the minimum and maximum values. Dots represent outlier cells. P-values were calculated using two-sided t-test with unequal variance. * < 0.05, ** < 0.01, *** < 0.001. **k)** Expression of *SNHG26* isoforms in primed (n=2), partial *naïve* (n=2), *naïve* (n=4) hESCs or 8CLCs (n=2) samples from a publicly available dataset^96^. RPKM: Reads Per Kilobase of transcript per Million reads; PXGL: medium with PD0325901, XAV939, Gö6983 and LIF.

As with mouse ESCs, we knocked-down *SNHG26*’s transcription in hESCs by CRISPRi. Contrary to results in mouse ESCs, in hESCs, no significant decrease was observed for the core pluripotency genes *OCT4* and *NANOG* (**Supplementary Fig. 16b**). As previously mentioned, mouse and human ESCs do not correspond to the same state of pluripotency, and *SNHG26* is more expressed in the *naïve* state. Thus, the use of primed hESCs limits the study and the full extent of *SNHG26* knock-down effect. Interestingly, the two markers of primed pluripotency *SALL4* and *OTX2* were down-regulated (24% and 33%, respectively), and expression of some *naïve* pluripotency markers was also decreased (40%, 32% and 42% for *TFCP2L1*, *REX1* and *DPPA4*, respectively) (**Supplementary Fig. 16b**). Deeper look at how overall gene expression was affected by *SNHG26* knock-down in hESCs by lrRNAseq revealed that additional markers of pluripotency were down-regulated, such as *DPPA3, UTF1* and *FGF5* (**Fig. 9c-d** and **Supplementary Data 13**). These results suggested that similar to its mouse ortholog, knock-down of *SNHG26* in hESCs was associated with acute pluripotency perturbation. Interestingly, GO-term enrichment analysis showed that genes up-regulated following *SNHG26* knock-down were linked to regulation of gene transcription and were tumor suppressor genes (**Fig. 9e** and **Supplementary Data 13**). This suggested that *SNHG26* lncRNA could act as a transcriptional regulator in hESCs, in keeping with its nuclear localization in mouse ESCs. On the opposite, down-regulated genes were associated to cell migration, cancer-related pathways and cancer resistance (**Fig. 9f** and **Supplementary Data 13**). This is interesting as *SNHG26* lncRNA has been shown to be up-regulated in many types of resistant cancers^98–102^, and genes from the pluripotency gene network are overexpressed in cancer, promoting stem cell-like properties and uncontrolled growth, thus driving tumor initiation, progression and resistance to treatment^103^. This suggests that *SNHG26* could have a role in tumorigenesis by promoting stem cell-like properties. Consistent with this notion, *SNHG26* has been shown to be a pivotal regulator of keratinocyte progenitors and their transition from an inflammatory phase to a subsequent proliferative phase during both mouse and human wound healing^104^.

Next, in order to validate the conserved role of *SNHG26* lncRNA in regulation of pluripotency, we used a pool of antisense oligonucleotides (ASOs) to knock-down *SNHG26* during reprogramming of human dermal fibroblasts to iPSCs (**Fig. 9g**). ASOs were employed due to the technical challenges associated with generating CRISPRi lines from fibroblasts. Transfection of ASOs throughout reprogramming resulted in a 70% reduction in the expression of *SNHG26* by day 12, relative to cells treated with non-targeting control ASOs (**Supplementary Fig. 16c**). Although no morphological differences were observed at the end of reprogramming factors induction (i.e. D10), iPSC colonies with *SNHG26* knock-down showed a lower density and size compared to reprogramming without and with control ASOs (**Supplementary Fig. 16d**). Quantification of NANOG, OCT4, and SOX2-positive colonies at D20 of reprogramming also demonstrated a reduction in the number of colonies following *SNHG26* knock-down, with noticeably smaller sizes than the control conditions (**Fig. 9h-i** and **Supplementary Fig. 16e**). These results demonstrated that *SNHG26* lncRNA has a conserved role in the acquisition of the pluripotent state, and that the function of this lncRNA is not limited to reprogramming of neural progenitor cells but also to fibroblasts.

Next, we asked whether we could find specific isoforms that mediate the function of human *SNHG26*. *SNHG26* has 22 human isoforms according to Ensembl database (release version 107, July 2022)^105^, with only 10 being expressed in human PSCs or early embryo datasets (**Supplementary Fig. 17** and **Supplementary Data 13**)^93,96,97,106–110^. We identified 4 that have a conserved exon-intron structure with the mouse *Snhg26.217*, and 4 resembling the *Snhg26.216* isoform (**Supplementary Fig. 17**). Among the isoforms resembling *Snhg26.217, SNHG26.201* has already been shown to regulate cell fate and proliferation of keratinocytes^104^. However, when looking at a dataset from early human embryonic development samples, only 6 of the resembling transcripts were detected, and *SNHG26.201* was lowly expressed (**Fig. 9j**). Strikingly, only *SNHG26.219,* which has a conserved structure with *Snhg26.217,* had the same expression pattern as its mouse ortholog, being highly expressed at the 8-cell stage (**Fig. 9j**). Moreover, *SNHG26.219* was the transcript the most expressed in *in vitro* 8 cell-like cells (**Fig. 9k**). These results suggest that *SNHG26.219* transcript may be the human ortholog of *Snhg26.217* and possess a similar function in the regulation of pluripotency. Further work is required to establish if this is the case.

Thus, the genomic conservation and identical expression profiles over several biological systems suggested that *Snhg26* lncRNA could share a conserved functional role in pluripotency regulation (refer to **Fig. 10** for a working model). Thus, lrRNAseq analysis of our optimized reprogramming systems enabled us to identify a functional isoform of a new lncRNA whose function is conserved in regulating the pluripotent state.

**Figure 10:**
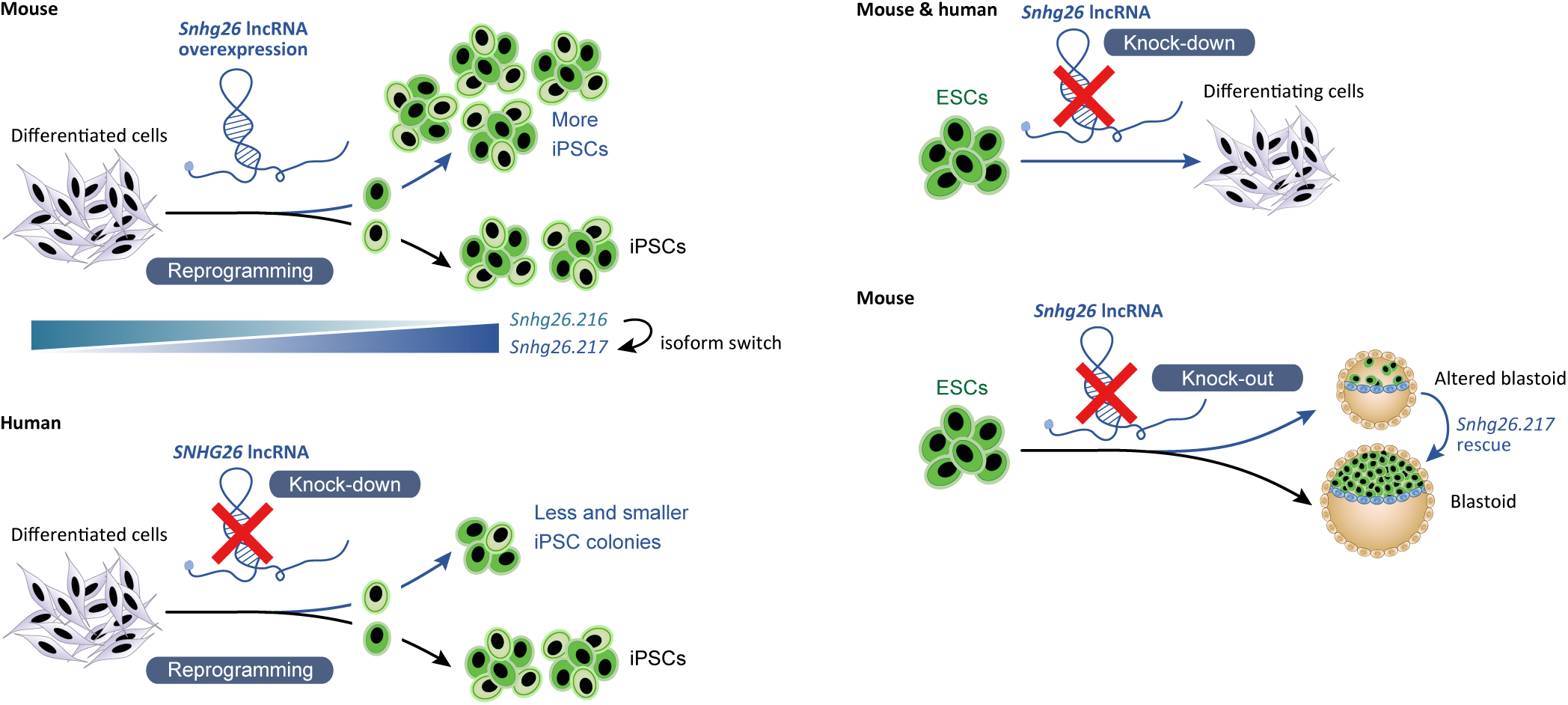
*Snhg26* is a conserved lncRNA that acts in the pluripotency regulatory network. Schematic representing the enhancing effect of *Snhg26* lncRNA on iPSC generation and the isoform switch that occurs during mouse reprogramming. Accordingly, knock-down of *SNHG26* during human reprogramming leads to less and smaller iPSC colonies. Comparatively, *Snhg26* knock-down promotes mouse and human ESC differentiation. *Snhg26* knock-out impairs mouse blastoids formation and generation of inner cell mass-like cells, which are both rescued by *Snhg26.217* isoform.

## Discussion

Previous studies have demonstrated that AS regulation is an important factor in cell fate determination, for instance during somatic cell reprogramming to pluripotency. Two reports^32,35^, using different reprogramming systems, have explored genome-wide AS across different timepoints of reprogramming and demonstrated that specific splicing patterns and regulators are linked to reprogramming. Taking advantage of full-length transcripts generated by long-read RNAseq, we explored this further and undertook the first lrRNAseq-mediated characterization of isoform diversity during the reprogramming of cells towards an induced pluripotent state. Our work identifies for the first time that isoform diversity and switching are highly implicated in the process. We also show that this is not limited to protein-coding RNA isoforms, but rather non-coding RNA isoform switching plays a major role in regulating gene function and reprogramming outcome. In fact, among genes that are affected by this IS control mechanism is *Srsf2*, which was also identified by these previous studies and others^29,31^ as an important AS regulator during reprogramming and pluripotency. Expression of non-coding isoforms was down-regulated and made way for expression of isoforms that lead to functional proteins. We also show that the splicing and isoform expression patterns of these regulators differ between our reprogramming systems and may affect their functional outcome. Both OKMS and OKS reprogramming systems have the same start (NPCs) and end (iPSCs) points but use different interplay between isoform expression and AS to achieve the same outcome. This also suggests that isoform diversification and switching promote functional outcomes that are favorable for reprogramming and exposes a level of isoform regulation that has not been shown before. Thus, in our study, we aimed at identifying such outcomes and their effect on the regulation of the pluripotent state.

We began by first devising an appropriate reprogramming system model that recapitulates known features of reprogramming in terms of timelines, gene expression profile and alternative splicing events. Running lrRNAseq at different timepoints of reprogramming, we brought to light the underestimated diversity of isoforms expressed during this process, with about 65% of the total detected isoforms being previously unannotated and many originating from new splice junctions (i.e. “Novel-Not-in-Catalog” transcripts). This reinforces the utility of lrRNAseq to address the shortcomings of short-read RNAseq in identifying full-length isoforms. Closer examination of isoforms showed that the transcripts that were classified as non-coding were regulated in the early and middle phases of both reprogramming systems. We found that most differentially abundant isoforms were at the transition to the early phase of reprogramming. By the middle phase of the process, a switch was observed in the abundance of alternatively spliced isoforms. We also found that less abundant spliced isoforms had a lower coding potential before this switch, and this hinted at a pressure to regulate specifically non-coding isoforms in the early phases of reprogramming. This demonstrates that non-coding isoforms undergoing isoform switch may have important functions during acquisition of pluripotency. These findings show that isoform diversification and switching of non-coding genes, such as lncRNAs, add a new level of regulation in pluripotency establishment.

Previous research has shown involvement of lncRNAs in pluripotency regulation^25,111–113^, but only a few examples of functionally tested lncRNAs exist. Moreover, unlike protein-coding genes, which give rise to a functional product, it is often difficult to identify the functional isoform and the mechanism of action of a lncRNA. Our study provides a new way of identifying functionally relevant lncRNAs in the context of pluripotency. By integrating our data with several datasets on reprogramming and pluripotency, we identified the *Snhg26* lncRNA that undergoes isoform switch during early reprogramming. The overexpression of the *Snhg26.217* isoform greatly increased the number of reprogramming cells and *bona fide* iPSCs. Furthermore, knock-down of human *SNHG26* resulted in lower reprogramming efficiency. Decrease of *Snhg26* in mouse and human ESCs down-regulated expression of several pluripotency markers, validating the conserved role for this new lncRNA in the regulation of pluripotency. Moreover, the conserved expression pattern of mouse and human *Snhg26* during embryo development and their common perturbance of the pluripotent state when absent is of interest, as it illustrates biological contexts in which the *Snhg26* lncRNA is evolutionary conserved. Indeed, *SNHG26* knock-down in hESCs up-regulated several tumor suppressor genes, linking its role in promoting stem cell features with several studies that showed its association with multiple cancer types. One of the limitations of this study is the use of a reprogramming system where transgenes are randomly integrated within the genome of the reprogramming cells. We can thus not exclude that this parameter could further affect the highlighted differences between OKMS and OKS reprogramming, nor the resulting enhanced reprogramming to iPSCs when *Snhg26* lncRNA is expressed. However, our system is based on a heterogenous cell population, and the effects of *Snhg26* on reprogramming kinetics were validated multiple times using different cell lines and reprogramming strategies.

In summary, it is important to note that only by analyzing isoform switching of *Snhg26* we were able to identify a functional isoform of this lncRNA. Given the current state of lncRNA annotations, which is highly complex and of unknown functional consequence, our study highlights the benefits of using lrRNAseq to recognize isoform diversity and switching and how in-depth analysis of such processes is key to identifying functionally relevant lncRNA isoforms. The dataset we generated also stands as a valuable resource for other researchers to study the role of alternative splicing and isoform diversity in the regulation of cell fate.

## Methods

### Generation of the destination vectors

For reprogramming, the *piggyBac* (PB) transposon vectors PB-TAC-OKMS (Addgene #80480), PB-TAC-OKS (OKS sequence was amplified from Addgene #19771 and reverse Gateway^TM^ cloned into PB-TAC-OKMS to replace the OKMS sequence), and PB-TAC-OSKM (Addgene #80481) containing a doxycycline-inducible TetO promoter, as well as a polycistronic reprogramming cassette with either *Oct4*, *Klf4*, *c-Myc* and *Sox2* genes (OKMS), *Oct4*, *Klf4* and *Sox2* genes (OKS), or *Oct4, Sox2, Klf4* and *c-Myc* (OSKM) and an *mCherry* reporter gene were kindly gifted by Dr. Andras Nagy (University of Toronto). The pCMV-hyPBase plasmid^114^ from the Sanger Institute was used to express the PB transposase.

For transgene overexpression, the PB transposon destination vectors PB-TetO-AIO carrying our genes of interest (e.g. *Snhg26*) were generated by digesting an in-house PB-TetO-Gateway plasmid and the PB-TAC-ERP2 (Addgene #80478) or PB-TAC-ERN (Addgene #80475) plasmids with MluI-HF® (New England Biolabs (NEB) #R3198) and SbfI-HF® (NEB #R3642), followed by ligation using a T4 DNA ligase (ThermoFisher Scientific #EL0016) according to the manufacturer’s instructions. They carry a doxycycline-inducible TetO promoter, a Gateway^TM^ cloning cassette as well as the constitutive reverse tetracycline-controlled TransActivator (rtTA) and a puromycin (P) or neomycin (N) resistance gene.

For CRISPRi-mediated knock-downs, an in-house dCas9-BFP-KRAB construct was PCR-amplified from the pHR-SFFV-dCas9-BFP-KRAB plasmid (Addgene #46911) using the Deoxynucleotide (dNTP) Solution Set (NEB #N0446S) and the Q5® High-Fidelity polymerase (NEB #M0491) according to the manufacturer’s instructions. The fragment was then inserted into the PB-TetO-AIO destination vector using Gateway^TM^ cloning (ThermoFisher Scientific #11791020) strategy according to the manufacturer’s instructions. The sequences of all plasmids were verified by Sanger sequencing and inducible expression validated by RT-qPCR. Complete sequences of all plasmids are available upon request.

### Cloning of the *Snhg26* and control transcripts into destination vectors

Total RNA was extracted from mouse embryonic stem cells (ESCs) and reversed-transcribed into cDNA using SuperScript™ III Reverse Transcriptase (ThermoFisher Scientific #18080044) according to the manufacturer’s instructions. Primers designed to target *Snhg26* transcript 217 (*Ensmust00000181574* or “*Snhg26.217*”) were used with the dNTP Solution Set (NEB #N0446S) and the Q5® High-Fidelity polymerase (NEB #M0491) according to the manufacturer’s instructions. *Snhg26* PCR-amplified cDNA transcript sequence was validated by Sanger sequencing and cloned into the PB-TetO-AIO destination vector using Gateway^TM^ cloning (ThermoFisher Scientific #11789020 & #11791020) strategy according to the manufacturer’s instructions. For *Snhg26* transcript 216 (*Ensmust00000268635* or “*Snhg26.216*”), the *Snhg26.217* containing plasmid was digested with BglII (NEB #R0144) and XcmI (NEB #R0533) to remove a part of exon 4. Primers to ligate back the plasmid were designed so that they also contain a missing part of the truncated exon 4 sequence. Those were added by ligation using a T4 DNA ligase (ThermoFisher Scientific #EL0016) according to the manufacturer’s instructions. Similarly to *Snhg26.217*, a cDNA antisense to the *Luciferase* gene (control condition) was PCR-amplified using primers designed to target the pGL3-Basic plasmid (Promega #E1751) and cloned by Gateway^TM^ cloning as described above. The sequences of all plasmids were verified by Sanger sequencing and inducible expression validated by RT-qPCR. Complete sequences of the resulting plasmids are available upon request (primer sequences are listed in **Supplementary Data 11**).

### Mouse ESC culture

ESC lines, including R1 (RRID: CVCL_2167) and Oct4-GFP (harboring a *GFP* reporter gene under the control of the *Oct4* gene promoter from Viswanathan *et al*.^115^) (both gifted by Dr. Andras Nagy, University of Toronto) were routinely maintained in a humidified incubator at 37°C under 5% CO2. Cells were propagated in 2i-LIF medium on 0.1% porcine gelatin (Sigma-Aldrich #G1890). 2i-LIF medium consists of 20ng/mL LIF (home-made from pGEX-2T-MLIF^116^), 3μM CHIR-99021 (Selleck Chemicals #S2924) and 1μM PD0325901 (Selleck Chemicals #S1036) in N2B27 media. N2B27 media consists of a 1:1 mixture of DMEM/F-12 media mix (ThermoFisher Scientific #11330057) supplemented with 1X N-2 supplement (ThermoFisher Scientific #17502048) and 20μg/mL human recombinant Insulin (ThermoFisher Scientific #12585014), and Neurobasal^TM^ media (ThermoFisher Scientific #21103049) supplemented with 1X B-27^TM^ supplement (ThermoFisher Scientific #17504044), 1mM sodium pyruvate (ThermoFisher Scientific #11360070), 2mM L-glutamine (ThermoFisher Scientific #35050061), 1X MEM Non-Essential Amino Acid solution (ThermoFisher Scientific #11140050) and 10^-4^M 2-Mercaptoethanol (ThermoFisher Scientific #21985023). Medium was refreshed every other day and ESCs were passaged every 2-3 days. For passaging, cells were treated with TrypLE^TM^ Express Enzyme (ThermoFisher Scientific #12604021) and passaged as single-cells (usually 1/10 ratio). All cell lines were negative for mycoplasma.

### Generation of Oct4-GFP neural progenitor cell (NPC) line

Oct4-GFP ESCs were dissociated into single-cells with TrypLE^TM^ Express Enzyme (ThermoFisher Scientific #12604021) and seeded on 0.1% porcine gelatin (Sigma-Aldrich #G1890) in FBS-LIF medium. FBS-LIF (FBSL) medium consists of DMEM (high glucose, ThermoFisher Scientific #11965118) supplemented with 15% Fetal Bovine Serum (FBS, ThermoFisher Scientific #12483020), 1mM sodium pyruvate (ThermoFisher Scientific #11360070), 2mM L-glutamine (ThermoFisher Scientific #35050061), 1X MEM Non-Essential Amino Acid solution (ThermoFisher Scientific #11140050), 10^-4^M 2-Mercaptoethanol (ThermoFisher Scientific #21985023) and LIF at 20ng/mL. The resulting primed cells were passaged at day 2 (D2) and day 4 in FBSL medium on 0.1% porcine gelatin using TrypLE^TM^ Express Enzyme. On D4, cells were seeded at a density of 1 x 10^3^ cells/cm^2^ in FBSL medium on 0.1% porcine gelatin. Twenty-four hours later, medium was switched to N2B27 medium with 35μg/mL human recombinant Insulin (ThermoFisher Scientific #12585014) and replaced every other day. At D10, cells were dissociated with TrypLE^TM^ Express Enzyme and transferred as single-cells into Corning^TM^ Costar^TM^ Ultra-Low Attachment microwell plates (Fisher Scientific #07-200-601) in Selection medium (2:1 mixture of N2B27 medium with 35μg/mL human recombinant Insulin supplemented with 20ng/mL human recombinant EGF (Peprotech #AF-100-15), 20ng/mL human recombinant FGF-2 (Peprotech #100-18B) and 500ng/mL heparin sodium salt (Sigma-Aldrich #H3393)) to promote neurosphere generation. Medium was replaced every other day and the neurospheres were passaged every 3-4 days with TrypLE^TM^ Express Enzyme until D20. Dissociated neurospheres were then seeded in NPC medium on Geltrex^TM^-coated plates (hESC-qualified, growth factor-reduced, ThermoFisher Scientific, #A1413302). NPC medium consists of N2B27 supplemented with 20ng/mL human recombinant EGF, 20ng/mL human recombinant FGF-2 and 500ng/mL heparin sodium salt. Medium was refreshed every other day and NPCs were passaged every 2-3 days. For passaging, cells were treated with StemPro™ Accutase™ Cell Dissociation Reagent (ThermoFisher Scientific #A1110501) and passaged as single-cells (usually 1/6-1/8 ratio). All cell lines were maintained in a humidified incubator at 37°C under 5% CO2 and were negative for mycoplasma.

### Human embryonic stem cells (hESCs) culture

Primed hESCs line H9 (RRID: CVCL_9773) were routinely maintained in a humidified incubator at 37°C under 5% CO2. Cells were propagated in mTeSR^TM^1 medium (StemCell Technologies #85850) on Geltrex^TM^-coated plates (hESC-qualified, growth factor-reduced, ThermoFisher Scientific #A1413302). Medium was refreshed every day and hESCs were passaged every 4-5 days. For passaging, cells were treated with Dispase (StemCell Technologies #07923) and passaged as small clumps using a Corning^TM^ Cell Lifter (Fisher Scientific #07-200-364) (usually 1/10 ratio). All cell lines were negative for mycoplasma. Human ESCs were obtained commercially, and all experiments pertaining to hESCs were approved by the Stem Cell Oversight Committee as outlined by Canadian Institutes of Health Research as per the Tri-Council Policy Statement: Ethical Conduct for Research Involving Humans (TCPS 2) and the Agreement on the Administration of Agency Grants and Awards by Research Institutions.

### Transfection of mouse ESCs and NPCs

ESC lines were seeded at a density of 13 x 10^3^ cells/cm^2^ on plates coated with 0.1% porcine gelatin (Sigma-Aldrich #G1890) in 2i-LIF medium. Alternatively, NPC lines were seeded at a density of 30 x 10^3^ cells/cm^2^ on Geltrex^TM^-coated plates (hESC-qualified, growth factor-reduced, ThermoFisher Scientific, #A1413302) in NPC medium.

The following day, cells were transfected with 200-500ng of hyPBase transposase and 1.8-2µg of PB-TetO-AIO constructs using Lipofectamine^TM^ Stem Transfection Reagent (ThermoFisher Scientific #STEM00015) according to the manufacturer’s instructions. For reprogramming cell lines, 500ng of hyPBase transposase were co-transfected with 750ng of PB-TAC-OKMS or PB-TAC-OKS vectors and 750ng of PB-TetO-AIO constructs using Lipofectamine^TM^ Stem Transfection Reagent according to the manufacturer’s instructions. Cells were then selected with 1μg/mL puromycin (Bio Basic #PJ593) for 4 days or 400μg/mL G-418 sulfate for 6 days.

### Reprogramming towards iPSCs from mouse cells

Stably transfected Oct4-GFP ESC-derived NPC cell lines were seeded at 3 x 10^3^ cells/cm^2^ on a 0.1% porcine gelatin (Sigma-Aldrich #G1890) coating in NPC medium supplemented with 2% FBS (ThermoFisher Scientific #12483020) to promote adhesion. The following day, medium was replaced with FBSL supplemented with 1µg/mL doxycycline hyclate (Sigma-Aldrich #D9891) to induce overexpression of the reprogramming factors. Medium was refreshed every other day until D6. From D6 (unless mentioned otherwise), cells were maintained either in FBSL or 2i-LIF medium and passaged every 2-4 days depending on colonies density. For passaging, cells were treated with TrypLE^TM^ Express Enzyme (ThermoFisher Scientific #12604021) and passaged as single-cells (1/3-1/30 ratio depending on density). mCherry+ and GFP+ cells were monitored using an Axio Vert.A1 (Zeiss) inverted fluorescence microscope with a 10X objective. Images were acquired with an Axiocam 305 color camera (Zeiss) and the ZEN 2 (blue edition) software (v3.9.100). Reprogramming cells were harvested using TrypLE^TM^ Express Enzyme for dissociation, washed with DPBS and pelleted for RNA extraction or resuspended in 500μL of DPBS for analysis using a BD Accuri^TM^ C6 Plus flow cytometer (BD Biosciences). iPSCs could readily be maintained in 2i-LIF medium after 20 days. All cell lines were negative for mycoplasma.

### Assessment of mouse iPSC pluripotency in iPSC-embryo chimeras

Prior to aggregation, iPS cells were grown in KSR + 2i medium^117^ consisting of high glucose DMEM with 15% KSR, 2mM GlutaMAX^TM^, 1mM Na pyruvate, 0.1mM non-essential amino acids, 0.1mM 2-Mercaptoethanol (Gibco), 1000U/mL LIF, 5μg/mL Insulin (Sigma #I0516), 1μΜ PD0325901 (StemGent #04-0006), and 3μΜ CHIR-99021 (StemGent #04-0004).

For OKMS/OKS and OKS + *Snhg26.217* reprogramming: Embryos from superovulated CD-1 (Charles River) females were collected at 2.5 days post-coitum (dpc). Zonae pellucidae of embryos were removed using acid Tyrode’s solution (Sigma) and clumps of cells from the iPS clones were obtained using Accutase (Millipore #SF006). Embryos were then aggregated with a clump of 6-10 iPSCs inside a depression well made in the dish with an aggregation needle (BLS Ltd). Aggregates were cultured overnight in microdrops of Life Global medium (Cooper Surgical LGGG-050) with 1mg/mL Protein Supplement (Cooper Surgical LGPS-020) covered by embryo-tested mineral oil (Cooper Surgical LGPO-500) at 37°C and 6% CO_2_. Embryos were transferred the following morning into the uteri of 2.5-dpc pseudo pregnant CD-1 females. Chimerism was evaluated at birth based on the presence of black eyes and at weaning by coat color pigmentation. CD-1 outbred albino stock was used as embryo donors for aggregation with iPSCs and as pseudo pregnant recipients. Animals were maintained on 12h light/dark cycle and provided with food and water *ad libitum* in individually ventilated units (Techniplast) in the specific pathogen-free facility at The Centre for Phenogenomics (TCP). All procedures involving animals were performed in compliance with the Animals for Research Act of Ontario and the Guidelines of the Canadian Council on Animal Care. Animal Care Committee reviewed and approved all procedures conducted on animals at TCP.

For OKS + control and OKS + *Snhg26.217* reprogramming: An iPSC clamp consisting of 4-5 cells of a clone was aggregated with two tdTomato mouse embryos at the 8-cell morula stage in a depression well of a small drop of EmbryoMax® Advanced KSOM Embryo media (Sigma-Aldrich #MR-101-D) under mineral oil. Embryos had ubiquitous tdTomato expression derived from the Ai14 strain (RRID: IMSR_JAX:007914). After overnight incubation at 37°C with 5% CO2, the blastocysts were transferred to the uteri of pseudo pregnant females. The embryos were dissected at E8-8.5 stage. Contribution of tdTomato negative iPSC-derived cells was visualized as negative cells/area in tdTomato-positive chimeric embryos. This was performed at the McGill Integrated Core for Animal Modeling (MICAM).

### Assessment of mouse iPSC pluripotency using teratoma formation assay

Prior to the transplantation iPSCs were cultured as described above for aggregation. On the day of transplantation, iPSCs were harvested and dissociated into a single-cell suspension using Accutase (Millipore #SF006). The pellets were washed in PBS, and resuspended in Matrigel^TM^ Basement Membrane (BD Biosciences #356234) - diluted 1 volume of Matrigel + 2 volumes of cold DMEM to get 5 × 10^6^ cells per injection site. Mice were anesthetized using isoflurane, 100μL of cell suspension in diluted Matrigel were injected subcutaneously into each hind limb. Mice were monitored to determine when the tumor is palpable and visible, measured with calipers every 2-3 days to log tumor growth. Once the tumor has met end point guidelines, mice were euthanized and tumors dissected analysis. Briefly, teratoma were fixed in formalin and embedded in paraffin. Tissue cuts were stained by immunohistochemistry and scanned by Axioscan 7 (Zeiss) with a 20X objective. All procedures involving animals were performed in compliance with the Animals for Research Act of Ontario and the Guidelines of the Canadian Council on Animal Care. Animal Care Committee reviewed and approved all procedures conducted on animals at TCP.

### Doubling time assays

ESC or NPC cell lines were seeded in replicates at 5 x 10^4^ cells/cm^2^ on a 0.1% porcine gelatin (Sigma-Aldrich #G1890) coating in 2i-LIF or NPC medium supplemented with 1% FBS (ThermoFisher Scientific #12483020) to promote adhesion. The following day, one replicate of cells was harvested using TrypLE^TM^ Express Enzyme (ThermoFisher Scientific #12604021) and counted using a BD Accuri^TM^ C6 Plus flow cytometer (BD Biosciences) as the starting timepoint (0h). Medium was replaced with 2i-LIF, NPC medium or FBSL medium supplemented with 1µg/mL doxycycline hyclate (Sigma-Aldrich #D9891) to induce transgenes overexpression. The cells were counted again after 24, 48, 72 and 120 hours. The total numbers of cells of 2-4 independent experiments were used to generate the average growth curves and to calculate the mean doubling times of the cell lines. Growth curves were represented with SEM using Microsoft Excel and doubling time graphs with STDs using GraphPad Prism (v7.04). The expression of an *mCherry* reporter gene allowed to calculate specifically the doubling time of the reprogramming cells. The cell specific growth rate (*µ*) was determined from the slope of the natural logarithm of cell count as a function of time and doubling time (DT) using the formula [*DT* = ln 2/*µ*]. Calculation used for doubling time was performed as follow: *DT* = (*T2*-*T1*) x ln(2)/ln(*q2*/*q1*) where *T* = time and *q* = quantity of cells. P-values between the different timepoints and doubling times were calculated using t-test (unpaired and unequal variance).

### Limiting dilution assays

Stably transfected Oct4-GFP ESC-derived NPC cell lines were seeded at 5, 25, 125, or 500 cells per well of a Falcon^TM^ 96-well Flat-Bottom microplate (Fisher Scientific #08-772-2C) on a 0.1% porcine gelatin (Sigma-Aldrich #G1890) coating in NPC medium. The following day, medium was replaced with FBSL medium supplemented with 1µg/mL doxycycline hyclate (Sigma-Aldrich #D9891) to induce the reprogramming into iPSCs. Medium was refreshed every other day for two weeks and switched at D18 to 2i-LIF medium until reprogramming completion. The number of wells negative for mCherry+ and GFP+ cells were quantified manually using an Axio Vert.A1 (Zeiss) inverted fluorescence microscope with a 10X objective after 12 and 27 days. In Microsoft Excel, the average percentage of negative wells was plotted on a semi-logarithmic graph in function of the number of cells seeded as an exponential line, with error bars representing the SEM. The Poisson distribution allowed to determine the number of cells (*x* = 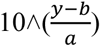, where *y* = 37, *a* = slope and *b* = y intercept) that must be seeded to have 37% of negative wells. The frequency (1/*x*) and percentage (1/*x* *100) of cells able to reach the iPSC stage were calculated based on three independent experiments (n=3). Frequencies of reprogramming cells and iPSCs were represented with STDs using GraphPad Prism (v7.04). P-values between the frequencies were calculated using t-test (unpaired and unequal variance).

### Flow cytometry

Cells were harvested using TrypLE^TM^ Express Enzyme (ThermoFisher Scientific #12604021) for dissociation, resuspended in 500μL of DPBS and filtered through a 35μm cell strainer (Fisher Scientific #08-771-23). Samples were loaded into a BD Accuri^TM^ C6 Plus flow cytometer (BD Biosciences). Flow cytometry data were collected with the C6 Plus analysis software and analyzed with FlowJo^TM^ (v10.8.1). Cell debris were gated apart using an FSC-A x SSC-A plot. Then, singlets were gated using an FSC-A x FSC-H plot and the gating for fluorescent reporter-positive cells was determined using a reporter-negative control sample as well as separated samples each positive for the different reporter genes. The total number of live single cells, as well as the percentages of mCherry+ and GFP+ cells and the volume used by the flow cytometer were collected directly from the .FCS files to calculate the density of mCherry+ and GFP+ cells as follow: *density of mC*ℎ*erry cells* = % *of live single mC*ℎ*erry cells* × 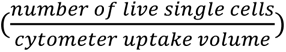. From this, the fraction of mCherry+ cells was calculated as follow: 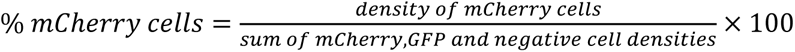. The average values of 3-9 independent experiments were used and represented with the SEM using Microsoft Excel. P-values were calculated using t-test (paired samples).

### ESC nuclear/cytoplasmic fractionation

Fractionation was performed following Gagnon *et al*.^118^. Briefly, R1 ESCs maintained in 2i-LIF medium on 0.1% porcine gelatin (Sigma-Aldrich #G1890) were expanded to a total of 3 x 10^6^ cells. Cells were dissociated using TrypLE^TM^ Express Enzyme (ThermoFisher Scientific #12604021). 1 x 10^6^ cells were kept for total RNA and the rest was processed for cell fractionation: Cells were resuspended in ice-cold RNAse-free hypotonic lysis buffer (10mM Tris pH7.5 (Bio Basic #TB0194), 10mM NaCl (Sigma-Aldrich #S3014), 3mM MgCl2 (Sigma-Aldrich #M4880), 0.3% (vol/vol) IGEPAL® CA-630 (Sigma-Aldrich #I3021), 10% (vol/vol) glycerol (Fisher Scientific #BP229-1), 400U/mL of buffer of Murine RNase Inhibitor (NEB #M0314)) and incubated for 10 minutes on ice. The lysate was vortexed briefly and centrifuged 3 minutes at 4°C at 1,000g. The supernatant (cytoplasmic fraction) was collected on ice, added to 1mL of fresh RNA precipitation solution (5% 3M sodium acetate pH5.5 (Sigma-Aldrich #S8750) and 95% ethanol) and incubated for 1 hour at -20°C. The solution was vortexed briefly and centrifuged 15 minutes at 4°C at 18,000g before washing the pellet in ice-cold 70% (vol/vol) ethanol and centrifuging 5 minutes at 4°C at 18,000g prior to extraction. The pellet (nuclear fraction) was resuspended in ice-cold RNAse-free hypotonic lysis buffer and centrifuged 2 minutes at 4°C at 200g three times prior to extraction. Non-fractioned cells, cytoplasmic and nuclear fractions were finally resuspended in RLT buffer (Qiagen #74106) to extract RNA using RNeasy Kit (Qiagen #74106) according to the manufacturer’s instructions. *Snhg26* lncRNA localization was evaluated by RT-qPCR by calculating the percentage of *Snhg26* cDNA PCR quantity in each fraction normalized to total *Snhg26* cDNA PCR quantity. *Xist, Malat1* and *Neat1* lncRNAs, and *Oct4* and *Klf2* mRNAs were used as controls for nuclear and cytoplasmic transcripts, respectively (qPCR primer sequences are listed in **Supplementary Data 13**). The average values of three independent experiments were used and represented with the STD using GraphPad Prism (v7.04). P-values were calculated using t-test (unpaired and unequal variance).

### RNAScope *in situ* hybridization

Wild-type or *Snhg26*^KO^ R1 ESC colonies were grown in 2i-LIF medium on microscope slides coated with 0.1% porcine gelatin (Sigma-Aldrich #G1890). After 3 days, ESCs were fixed with 4% paraformaldehyde (Fisher Scientific #AA433689L) for 30 minutes at room temperature, sequentially dehydrated using 50%, 70% and 100% ethanol for 5 minutes at room temperature and stored at -20°C. *In situ* hybridization of a set of probes targeting *Snhg26* transcript (Advanced Cell Diagnostics #1574981) was performed using the RNAScope® Multiplex Fluorescent Reagent Kit v2 (Advanced Cell Diagnostics #323100) according to the manufacturer’s instructions. Briefly, slides were sequentially re-hydrated using 100%, 70% and 50% ethanol for 5 minutes at room temperature. Cells were then permeabilized using 0.1% Tween 20 (Fisher Scientific #BP337-500). Slides were incubated for 10 minutes at room temperature after addition of H2O2. Protease III digestion was performed for 10 minutes at room temperature. The *Snhg26* probes were hybridized for 2 hours at 40°C. RNAScope® amplification steps with preamplifiers AMP1 (30 minutes), background reducers AMP2 (30 minutes) and amplifiers AMP3 (15 minutes) were then performed at 40°C. Amplifiers were conjugated to 1:2000 TSA Vivid Fluorophore 650 (Advanced Cell Diagnostics #323273). Nuclear staining was performed using DAPI from the RNAScope® Kit and slides were mounted in ProLong^TM^ Gold Antifade Mountant (ThermoFisher Scientific #P36930). Imaging was performed with an Axio ObserverZ.1 (Zeiss) microscope with LSM900 confocal head using 3µm z-steps. LD C-Apochromat 40X 1.1 water objective was used and images were acquired with the ZEN (blue edition) software (v3.3). The following parameters were used: LSM scan speed 4; DAPI 500V detector gain and 3.5% laser wavelength; Cy5 (TSA Vivid Fluorophore 650) 500V detector gain and 0.2% laser wavelength. Stacks were reconstructed using the Z project plugin in Fiji (ImageJ v2.14.0 Fiji).

### Knock-down of *Snhg26* in mouse and human ESCs

The inducible knock-down cell lines were generated using CRISPRi-based methods following Gilbert *et al*.^92^. Single guide RNAs (sgRNAs) were designed using the CRISPR 10K Tracks of the UCSC genome browser^119^ and ordered from IDT^TM^. They were resuspended and annealed before their ligation into an in-house PB-U6-Gateaway plasmid digested beforehand with BbsI-HF® (NEB #R3539) using a T4 DNA ligase (ThermoFisher Scientific #EL0016) according to the manufacturer’s instructions (sgRNA sequences are listed in **Supplementary Data 13**). The sequences of all plasmids were verified by Sanger sequencing. Complete sequences of all plasmids are available upon request.

For mouse *Snhg26* knock-down, 400ng of hyPBase transposase were co-transfected with 850ng of PB-TetO-AIO-dCas9-BFP-KRAB vector and 350ng of control or *Snhg26*-targeting sgRNA constructs using Lipofectamine^TM^ Stem Transfection Reagent according to the manufacturer’s instructions. Cells were then selected with 1μg/mL puromycin (Bio Basic #PJ593) for 4 days or 400μg/mL G-418 sulfate for 6 days. The mouse ESC lines were seeded prior to the knock-down at 2.6 x 10^4^ cells/cm^2^ on 0.1% porcine gelatin (Sigma-Aldrich #G1890) in 2i-LIF medium. The knock-down was induced by adding 1 µg/mL doxycycline hyclate (Sigma-Aldrich #D9891) and ESCs were harvested after 5 days for RNA extraction. For human *SNHG26* knock-down, primed hESC lines were seeded at a density of 2.6 x 10^4^ cells/cm^2^ on Geltrex^TM^-coated plates (hESC-qualified, growth factor-reduced, ThermoFisher Scientific, #A1413302) in mTeSR^TM^1 medium (StemCell Technologies, #85850) supplemented with 10µM Y-27632 (Tocris Bioscience #1254). The following day, medium was replaced by fresh mTeSR^TM^1 medium and cells were co-transfected with 500ng of hyPBase transposase, 1µg of PB-TetO-AIO-dCas9-BFP-KRAB vector and 875ng of control or four *SNHG26-*specific pooled sgRNAs using Lipofectamine^TM^ Stem Transfection Reagent (ThermoFisher Scientific #STEM00015) according to the manufacturer’s instructions. Cells were then selected with 500ng/mL puromycin (Bio Basic #PJ593) for 6 days. The hESCs were seeded prior to the knock-down on Geltrex^TM^-coated plates (hESC-qualified, growth factor-reduced, ThermoFisher Scientific, #A1413302) in mTeSR^TM^1 medium (StemCell Technologies, #85850). The knock-down was induced by adding 1 µg/mL doxycycline hyclate and ESCs were harvested after 2 or 3 days for RNA extraction.

*Snhg26* and *SNHG26* knock-down efficiencies as well as the mean relative fold-change in pluripotency-related mRNAs expression were measured via RT-qPCR by comparing to the control condition (non-targeting sgRNA) (qPCR primer sequences are listed in **Supplementary Data 13**). The average values of 2-3 independent experiments were used and represented with the STD using GraphPad Prism (v7.04). P-values were calculated using t-test (unpaired and unequal variance).

For analysis of global gene expression through lrRNAseq: The gene expression values in transcripts per million reads (TPM) of hESCs with control or *SNHG26* sgRNAs at D2 or D3 were averaged (**Supplementary Data 13**). Significantly up- or down-regulated genes were established either as genes with a fold change ≥ 1.5 or ≤ 0.67, respectively, and adjusted p-value ≤ 0.05 (any reference to adjusted p-values throughout the text is done using the Benjamini-Hochberg (FDR) method); or as genes with a Shannon entropy score below tenth percentile cutoff of the entropy scores distribution as previously described^16^. The Shannon entropy modelling to compute an expression-specificity index for each gene was generated as previously described^16,120^. Briefly, for each gene, a relative expression (Ri) value was calculated per timepoint (i.e. D2 or D3) and per condition (i.e. control or *SNHG26* sgRNAs), where Ri = (expression per timepoint or average expression per condition)/sum of TPM values in all the timepoints or all the conditions. The entropy index score (Hi) across all samples or groups was calculated as Hi = −1 × Σ(Ri × log2(Ri)). Represented down-regulated genes have a significance based either on fold change and adjusted p-value or on Shannon entropy score. Heat map expression of significant genes (n=1,794) was generated using the TPM expression values of these genes using GraphPad Prism (v7.04), setting the maximum color for values ≥ 15 TPM. Represented up- and down-regulated genes have a significance based only on Shannon entropy score (**Supplementary Data 13**). GO-term enrichment was assessed using the Database for Annotation, Visualization and Integrated Discovery (DAVID)^121^ (v2022q3) on the genes significantly up- or down-regulated based on the Shannon entropy score only. Significant GO-terms were selected based on a fold enrichment ≥ 1.5, a p-value ≤ 0.05, and an FDR ≤ 0.1 (**Supplementary Data 13**).

### Reprogramming of human fibroblasts towards iPSCs

Human reprogramming was performed using mRNAs generated by the HiScribe® T7 mRNA Kit with CleanCap® Reagent AG (NEB #E2080S) using constructs for *OCT4, SOX2, KLF4, C-MYC, LIN28* and nuclear-destabilized *GFP* (NDG) described in Mandal *et al*.^122^. BJ fibroblasts (ATCC #CRL-2522) were plated at 20,000 cells per well of a 24-well plate and transfected with 150ng of mRNA cocktail containing a 3:1:1:1:1:1 molar ratio for *OCT4, SOX2, KLF4, C-MYC, LIN28*, and *NDG* for 10 days with Lipofectamine^TM^ RNAiMAX (ThermoFisher Scientific #13778030). For the first 12 days, cells were cultured in NutriStem® hPSC XF GF-free (Sartorius #06-5100-01-1A) that was conditioned on mitotically arrested human neonatal dermal fibroblasts (amsbio #DFN-F) treated with mitomycin C for 3 hours. Starting at D13, cells were cultured in in-house generated MEF-conditioned media on mitotically-arrested mouse embryonic fibroblasts treated with mitomycin C for 3 hours and generated by the Lunenfeld-Tanenbaum Research Institute.

### Knock-down of *SNHG26* in human reprogramming

Knock-down of *SNHG26* during human reprogramming began with transfection of parallel, antisense oligonucleotides (ASOs) control or targeting *SNHG26* lncRNA (ASOs were custom-ordered and generated by IDT^TM^, sequences are listed in **Supplementary Data 13**) at D3 using Lipofectamine^TM^ RNAiMAX (ThermoFisher Scientific #13778030) with 30nM of ASO. Knock-down was subsequently performed every 3 days. Cells were passaged 1/3 at D13 and fixed at D20 before staining by immunofluorescence with NANOG (R&D Systems #AF1997), OCT4 (Santa Cruz Biotechnology #sc-9081), and SOX2 (R&D Systems #MAB2018) antibodies. Nuclear staining was performed using DAPI (Sigma #D9542-1MG) and whole-well immunofluorescence imaging was performed with the InCell Auto 3000 (GE Healthcare). Quantification of NANOG, OCT4, and SOX2-positive colonies was performed manually. Quantification of OCT4-positive colonies area was performed in Fĳi (ImageJ v1.54p Fiji) by smoothing granularity with a *Gaussian blur* function at sigma = 4μm. Thresholding was performed by adjusting the upper slider to 40. Threshold value was visually validated to ensure complete segmentation of OCT4-positive colonies while minimizing background inclusion. Noise was removed by using the *open* function once, touching colonies were then separated by the *watershed* function, and *particle size* was set > 5,000μm^2^ to automatically count the regions of interest. The area values of 175 to 1,042 colonies were averaged and represented with the SEM using Microsoft Excel. P-values were calculated using t-test (unpaired unequal variance). *SNHG26* knock-down efficiency was measured at D12 via purifying the RNA using Qiagen RNeasy Kit (Qiagen #74106) and reverse-transcription using Maxima H-minus Reverse (ThermoFisher Scientific #EP0751), followed by qPCR with the LightCycler 480 SYBR Green kit (Roche #04707516001), by comparing to the control condition (control ASOs) (qPCR primer sequences are listed in **Supplementary Data 13**).

### *Snhg26* knock-out (KO) in mouse ESCs

*Snhg26*^KO^ R1 ESC lines were established as described in Ran *et al*.^123^, using two sgRNAs to remove the full fourth exon of *Snhg26*. SgRNAs were designed using the CRISPR 10K Tracks of the UCSC genome browser^119^ and ordered from IDT^TM^. 0.350ng of annealed sgRNA pairs were cloned in 50ng of the pCCC backbone^124^ (gifted by Dr. Francis Lynn, University of British Columbia) by BbsI HF® digestion (NEB #R3539) followed by ligation using a T4 DNA ligase (ThermoFisher Scientific #EL0016) according to the manufacturer’s instructions (sgRNA sequences are listed in **Supplementary Data 13**). The sequences of all entry clone inserts were verified by Sanger sequencing. Complete sequences of all plasmids are available upon request.

For targeting with the CRISPR-Cas9 system, ESC lines were seeded at a density of 13 x 10^3^ cells/cm^2^ on plates coated with 0.1% porcine gelatin (Sigma-Aldrich #G1890) in 2i-LIF medium. The following day, cells were transfected with 100ng of hyPBase transposase, 200ng of transposon construct corresponding to either the control cDNA of *Luciferase-antisense*, *Snhg26.217* or *Snhg26.216*, and 800ng of each sgRNA construct using Lipofectamine^TM^ Stem Transfection Reagent (ThermoFisher Scientific #STEM00015) according to the manufacturer’s instructions. Cells were then selected with 1μg/mL puromycin (Bio Basic #PJ593) for 4 days. Five days post-selection, cells were dissociated as single-cells, counted and seeded at one cell per well of a Falcon^TM^ 96-well Flat-Bottom Microplate (Fisher Scientific #08-772-2C) with a multichannel pipettor for colony isolation. Medium was replaced every other day and wells containing single colonies were selected by eye using an Axio Vert.A1 (Zeiss) inverted microscope with a 10X objective. One week post-seeding, clones were split, expanded and frozen prior to screening and further characterization. All cell lines were maintained in 2i-LIF medium on 0.1% porcine gelatin, in a humidified incubator at 37°C under 5% CO2 and were negative for mycoplasma.

For *Snhg26*^KO^ ESC clone screening and validation, genomic DNA (gDNA) was extracted using gSYNC^TM^ DNA Extraction Kit (Geneaid #GS100) following the manufacturer’s instructions, with 100μg/mL RNaseA treatment (ThermoFisher Scientific #EN0531) 20 minutes at room temperature. DNA was quantified using a NanoDrop^TM^ 2000 Spectrophotometer (ThermoFisher #ND-2000C) and 300ng of DNA were loaded onto a 1% Agarose A (Bio Basic #D0012) gel to verify the DNA integrity and purity.

Assessment of ESC clones genotype regarding *Snhg26* locus was performed by PCR on 2ng of gDNA using the dNTP Solution Set (NEB #N0446S) and the Taq DNA polymerase (NEB #M0267) according to the manufacturer’s instructions. 15-25μL of PCR-amplified product was loaded on a 1% agarose gel with 5μL of the GD 1kb plus DNA Ladder RTU (FroggaBio #DM015-R500F) and revealed using an Azure 200 gel image (Azure Biosystems). PCR-amplification of *Snhg26* fourth exon screening followed by Sanger sequencing validation was used to assess the copy number of *Snhg26* allele in each clone (genotyping primer sequences are described in **Supplementary Data 13**).

### Blastoid differentiation and characterization

The blastoids were generated following a protocol adapted from Jana *et al.*^125^. Briefly, wild-type or *Snhg26*^KO^ mouse ESC lines routinely cultured in 2i-LIF medium were seeded at 9.4 x 10^3^ cells/cm^2^ on a 0.1% porcine gelatin (Sigma-Aldrich #G1890) coating in serum + LIF (SL) medium and passaged three times using TrypLE^TM^ Express Enzyme (ThermoFisher Scientific #12604021). On the third passage, SL medium was supplemented with 1μM PD0325901 (Selleck Chemicals #S1036) for 48 hours. Then, an Aggrewell^TM^ 400 plate (StemCell Technologies #34411) was treated with Anti-Adherence Rinsing Solution (StemCell Technologies #07010) as per the manufacturer’s instructions, and 48,000 cells were seeded per well in RBF medium. RBF media was composed of 12.5g/L GMEM with 10% FBS (vol/vol), 32.7mM NaHCO3, 1mM sodium pyruvate, 0.1mM non-essential amino acids, 0.1mM β-Mercaptoethanol, supplemented with 1μM retinoic acid, 25ng/mL bFGF and 20ng/mL BMP4. 24 hours later, 500µL of RBF medium was carefully added to each well. On the following day, 500µL of RBF medium containing CHIR-99021 (Selleck Chemicals #S2924) was added to each well for a final concentration of 3µM.

72h post seeding, brightfield images of the blastoids were acquired using an Axio Vert.A1 (Zeiss) microscope with a 5X objective coupled to an Axiocam 305 color camera (Zeiss) and the ZEN 2 (blue edition) software (v3.9.100). Blastoids were then carefully transferred from into two 1.5mL Eppendorf® LoBind® microcentrifuge tubes (Avantor #CA80077-230) using ART^TM^ wide bore filtered pipette tips (ThermoFisher Scientific #2079G). They were washed with DPBS (ThermoFisher Scientific #14190250) once by centrifuging 2 minutes at 300g at room temperature. Finally, blastoids were fixed with 4% paraformaldehyde (methanol-free, Fisher Scientific #AA433689L) for 30 minutes at 4°C under light agitation and rinsed in DPBS three times. Permeabilization was performed in DPBS containing 0.1% Tween (Fisher Scientific #BP337-500) (DPBS-T 0.1%) for 10 minutes and blastoids were blocked for 1 hour in 5% donkey serum (Sigma-Aldrich #D9663) diluted in the DPBS-T 0.1%. Primary antibodies against SOX2 (Cell Signaling Technology #23064) and GATA6 (Novus Biologicals #AF1700) were respectively diluted 1:100 and 1:80 in the blocking solution and incubated at 4°C overnight under light agitation. Blastoids were then rinsed three times in DPBS-T 0.1% for 5 minutes and incubated with secondary antibodies Alexa^TM^ Fluor 488 anti-goat (ThermoFisher Scientific #A21467) and 594 anti-rabbit (ThermoFisher Scientific #A21207) diluted 1:500 in the blocking solution for 60 minutes at RT. Following 3 rinsings with DPBS-T 0.1%, samples were incubated with DAPI (Sigma-Aldrich #D9542) diluted 1:200 for 7 minutes at RT, before 3 rinsings in DPBS and mounting on Fisherbrand^TM^ Superfrost^TM^ Plus microscope slides (Fisher Scientific #12-550-15) on which two SecureSeal^TM^ Imaging Spacers (Sigma-Aldrich #GBL654002-100EA) were placed and filled each with 10µL containing the stained blastoids in ProLong^TM^ Gold Antifade Mountant (ThermoFisher Scientific #P36930).

Imaging was performed with an Axio ObserverZ.1(Zeiss) microscope with LSM900 confocal head using 2μm z-steps. LD C-Apochromat 40X 1.1 water objective was used and images were acquired with the ZEN (blue edition) software (v3.3). The following parameters were used: LSM scan speed 6; DAPI 575V detector gain and 3% laser wavelength; 488 600V detector gain and 3% laser wavelength; 594 540V detector gain and 2.8% laser wavelength. Stacks were reconstructed using the Z project plugin in Fĳi (ImageJ v2.14.0 Fiji). Channels on the reconstructed Z images were split and analyzed separately. In Fiji, the *Freehand* tool was used to manually draw the outlines of each blastoid and measure the area. The total number of cells per blastoids was quantified manually based on DAPI-positive cells. SOX2- and GATA6-positive cells were counted manually. The average values of area, total number of cells, SOX2-or GATA6-positive cells in the wild-type (n=9-16), *Snhg26*^KO^ + control (n=9-12), *Snhg26*^KO^ + *Snhg26.217* (n=10-12), or *Snhg26*^KO^ + *Snhg26.216* (n=10-14) blastoids were used and represented as box plots with the minimal and maximal values using GraphPad Prism (v7.04). P-values were calculated by ordinary one-way ANOVA.

### RNA extraction and reverse transcription-qPCR

Cell lysates were homogenized with QIAshredder columns (Qiagen #79656) and total RNA was extracted using RNeasy Kit (Qiagen #74106) according to the manufacturer’s instructions. RNA was quantified using a NanoDrop^TM^ 2000 Spectrophotometer (ThermoFisher #ND-2000C) and then DNase-treated using DNaseI (ThermoFisher Scientific #EN0521) according to the manufacturer’s instructions. 300ng of DNase-treated and non-treated RNA were loaded onto a 1% Agarose A (Bio Basic #D0012) gel to verify the RNA integrity and purity. cDNA was synthesized from 500ng of RNA by oligo-dT and random priming with iScript^TM^ gDNA Clear cDNA Synthesis Kit (Bio-Rad #1725035), according to the manufacturer’s instructions. cDNA was diluted ten times and qPCRs were performed using SYBR^TM^ Select Master Mix (ThermoFisher Scientific #4472908) on a CFX384 Touch real-time PCR detection system (Bio-Rad). Raw data was retrieved using the Bio-Rad CFX Maestro (v5.3.022.1030) software. Relative gene quantification was analyzed based on the 2^-ΔΔCt^ method by applying the comparative cycle threshold (Ct) method on Microsoft Excel. Relative expression levels were normalized to *Rpl27 or Rps13* and to *RPS13* housekeeping genes for mouse and human cells analyses, respectively (qPCR primer sequences are listed in **Supplementary Data 13**).

### Library preparation and lrRNAseq, processing and alignment

Cell lysates were homogenized with QIAshredder columns (Qiagen #79656) and total RNA was extracted using RNeasy Kit (Qiagen #74106) according to the manufacturer’s instructions (RNAseq samples are listed in **Supplementary Data 1**). RNA was quantified using a NanoDrop^TM^ 2000 Spectrophotometer (ThermoFisher #ND-2000C) and then DNase-treated using TURBO^TM^ DNase (ThermoFisher Scientific #AM2238) according to the manufacturer’s instructions. 300ng of DNase-treated and non-treated RNA were loaded onto a 1% Agarose A (Bio Basic #D0012) gel to verify the RNA integrity and purity. RNA was also quantified using the Qubit™ RNA Broad Range Kit (ThermoFisher Scientific #Q10210) on Qubit™ 4 Fluorometer (ThermoFisher Scientific #Q33238) according to the manufacturer’s instructions. RNA quality was further verified using an RNA 6000 Nano Kit (Agilent Technologies #5067-1511) on a 2100 Bioanalyzer instrument (Agilent Technologies #G2939B) according to the manufacturer’s instructions. Samples all had RNA integrity numbers of 7.8 or more.

For samples sequenced on R9 Flow Cells, polyadenylated RNA was enriched from 7.5-20μg of total RNA using NEBNext® Poly(A) mRNA Magnetic Isolation Module (NEB #E7490) according to the manufacturer’s instructions. Poly(A)+-enriched RNA was quantified using the Qubit™ RNA High Sensitivity Kit (ThermoFisher Scientific #Q32852) on Qubit™ 4 Fluorometer according to the manufacturer’s instructions and poly(A)+-enriched RNA smear quality was verified using an RNA 6000 Pico Kit (Agilent Technologies #5067-1513) on a 2100 Bioanalyzer instrument according to the manufacturer’s instructions. Libraries were generated using 55-100ng of poly(A)+ RNA and a Direct cDNA Sequencing Kit (Oxford Nanopore Technologies (ONT) #SQK-DCS109), with each sample barcoded with the Native Barcoding Expansion (ONT #EXP-NBD104), according to the manufacturer’s protocol: “Direct cDNA Native Barcoding (SQK-DCS109 with EXP-NBD104 and EXP-NBD114)” version DCB_9091_v109_revF_04Feb2019. Barcoded samples were pooled, loaded on a MinION sequencer (ONT) using eight R9.4.1 Flow Cells (ONT #FLO-MIN106D), and sequenced using the MinKNOW^TM^ (v19) software. Purification of DNA between each step was performed with KAPA HyperPure Beads (Roche #08963835001). R9.4.1 Flow Cells were washed and reloaded with the Flow Cell Wash Kit (ONT #EXP-WSH003) to increase sequencing depth obtained per Flow Cell.

For samples sequenced on R10 Flow Cells, Ligation sequencing amplicons - Native Barcoding Kit 24 V14 (ONT #SQK-NBD114.24) was used with 50ng of total RNA. Each sample was assigned a specific barcode. Briefly, an oligo-dT VN Primer (VNP, sequence: 5’-5phos/ACTTGCCTGTCGCTCTATCTTCTTTTTTTTTTTTTTTTTTTTVN-3’, where V=A, C, or G, and N=A, C, G, or T) was used to reverse-transcribe poly(A) RNA into cDNA with the Maxima H Minus Reverse Transcriptase (RT) (ThermoFisher Scientific #EP0751). A Strand Switching Primer (SSP, sequence: 5’-TTTCTGTTGGTGCTGATATTGCTmGmGmG-3’, where mG = 2’ O-Methyl RNA bases) was used for strand switching during the RT reaction. Libraries were PCR-amplified for 14 PCR cycles with ONT’s cDNA Primers (cPRM) (Forward primer sequence: 5’-ATCGCCTACCGTGACAAGAAAGTTGTCGGTGTCTTTGTGACTTGCCTGTCGCTCTATCTTC-3’, Reverse primer sequence: 5’-ATCGCCTACCGTGACAAGAAAGTTGTCGGTGTCTTTGTGTTTCTGTTGGTGCTGATATTGC-3’) and the LongAmp® Taq 2X Master Mix (NEB #M0287) or the NEBNext Ultra II Q5 Master Mix (NEB #M0544L), following ONT’s SQK-PCS109 kit recommendations for PCR incubation times and temperatures. cDNA was quantified using the Qubit™ 1X dsDNA High Sensitivity Kit (ThermoFisher Scientific #Q33230) on Qubit™ 4 Fluorometer according to the manufacturer’s instructions and cDNA smear quality was verified using a DNA 12000 Kit (Agilent Technologies #5067-1508) on a 2100 Bioanalyzer instrument (Agilent Technologies #G2939B) according to the manufacturer’s instructions. The blunt DNA ends were prepared using the NEBNext® Ultra^TM^ II End Repair/dA-Tailing module (NEB #E7546), adding a 3’ dA tail and phosphorylating the 5’ end. Barcodes were ligated to the DNA fragments with a Blunt/TA ligase (NEB #M0367), barcoded samples were pooled and motor protein adapters were finally ligated with the NEBNext® Quick Ligation Module (NEB #E6056). Purification of DNA between each step was performed with KAPA HyperPure Beads (Roche #08963835001). Library was loaded on a PromethION 2 Solo (ONT) sequencer using R10.4.1 Flow Cells (ONT #FLO-PRO114M) and sequenced using the MinKNOW^TM^ (v23.07.15) software.

Raw fast5 files from sequencing with R9.4.1 Flow Cells were converted to the more recent pod5 format using the pod5-file-format software from ONT’s “nanoporetech” GitHub page (https://github.com/nanoporetech/pod5-file-format). pod5 files were then basecalled using Guppy (v6.5.7) using super accurate settings with the guppy_basecaller command and the dna_r9.4.1_450bps_sup.cfg configuration file. Barcodes were detected and reads were demultiplexed with the guppy_barcoder command using the –barcode_kits “EXP-NBD104 EXP-NBD114” argument. Raw pod5 files from sequencing with R10.4.1 Flow Cells were basecalled with Guppy (v6.5.7) using super accurate settings with the guppy_basecaller command and the dna_r10.4.1_e8.2_400bps_sup.cfg configuration file. Barcodes were detected and reads were demultiplexed with the guppy_barcoder command using the – barcode_kits “SQK-NBD114-24” argument. The resulting fastq reads were aligned to the GRCm38/mm10 mouse and GRCh38/hg38 human reference genomes with Minimap2^126^ (v2.24-r1122) using the following parameters: -aLx splice –cs = long. Aligned reads were quality-controlled with the NanoPlot software^127^ (v1.43.2).

### Mouse transcriptome assembly and isoform classification

To properly assemble the transcriptome file, reads from each sample alignment file were oriented according to transcript’s proper strand relative to the genome using alignment files obtained from Minimap2. To do so, we used the following strategy per sample file:

1. Reads were first filtered to keep reads with MAPQ > 13.
2. Multi-exonic/split reads (i.e. reads with splicing information) were re-oriented to reflect gene orientation with respect to the genomic strand. Minimap2 aligned and contain a “ts” tag to reflect transcript strandedness or sequence orientation (i.e. sense (+) versus antisense (-) sequence) but not the genomic strand. So here, we divided these split reads into 4 categories:

a. Reads with a ‘ts:A:+’ tag, which represents sense transcript sequence, and a forward read orientation with respect to the genome (i.e. SAM flag = 0) were isolated using piped “samtools view -F 16” (Samtools v1.17) and “grep -E ‘ts:A:\+’” functions and maintained in the same read orientation. They were given an ‘XS:A:+’ tag designation by modifying the ‘ts:A:+’ tag field using the “sed” function within the SAM alignment files. This is done to reflect a forward gene and read orientation with respect to the genome.
b. Reads with an ‘ts:A:+’ tag and a reverse read orientation with respect to the genome (i.e.
c. SAM flag = 16) were isolated using piped “samtools view -f 16” and “grep -E ‘ts:A:\+’” functions and maintained in the same read orientation. However, they were given an ‘XS:A:-‘ instead of the ‘ts:A:+’ tag to switch the gene orientation to reflect antisense or reverse gene orientation with respect to the genome.
d. Reads with a ‘ts:A:-’ tag, which represent antisense transcript sequence, and a forward read orientation with respect to the genome (i.e. SAM flag = 0), were isolated using piped “samtools view -F 16” and “grep -E ‘ts:A:-’” functions and re-oriented by changing the SAM flag to 16 (i.e. reverse read orientation) using the “sed” function in terminal and were given an ‘XS:A:-‘ designation as done above, to reflect both a reverse gene and read orientation with respect to the genome.
e. Reads with a ‘ts:A:-’ tag and a reverse read orientation with respect to the genome (i.e. SAM flag = 16), were isolated using piped “samtools view -f 16” and “grep -E ‘ts:A:-’” functions and re-oriented by changing the SAM flag to 0 (i.e. forward read orientation) using the “sed” function in terminal. However, they were given an ‘XS:A:+‘ designation as done above, to switch the gene orientation to reflect sense gene orientation with respect to the genome.
3. For mono-exonic transcripts, the reads were oriented with the Pychopper software from ONT’s “epi2me-labs” github page (https://github.com/epi2me-labs/pychopper).
4. All reads from the above filtered and re-oriented files were merged using Samtools to generate one alignment file per sample.

The transcriptome assembly was performed on each sample BAM file using the TAMA software (tc_version_date_2022_02_22)^128^, with the tama_collapse.py script and “-x no_cap -c 60 -i 75 -icm ident_map -sj sj_priority” parameters. Transcriptomes derived from each sample were then merged using the tama_merge.py script to produce one assembled transcriptome BED file for the whole dataset. To obtain GTF files, we used the tama_convert_bed_gtf_ensembl_no_cds.py script to convert TAMA generated BED files to GTF files. The resulting GTF files were then processed with the isoform classification pipeline of the SQANTI3 software^55^ (v5.1.1) using the sqanti3_qc.py script and the “--CAGE_peak” and “--polyA_motif_list” options to identify transcription start and end sites, respectively, to classify isoforms using GRCm38/mm10 as reference genome and Ensembl’s gene annotations version 99 as reference transcriptome. The --isoAnnotLite argument was used with Ensembl’s annotations version 86 to generate a GFF3 file containing functional and structural annotations for novel isoforms (available as **Supplementary Data 8**). Isoforms that are considered artifacts were then filtered using SQANTI3 filter with the “Rules filter” using default rules and sqanti3_filter.py script. Filtered-out reference isoforms were then rescued using SQANTI3 sqanti3_rescue.py script and the “rules” argument. The SQANTI3 produced classification file included the structural categories and subcategories, predictions for RT switching, coding potential and non-sense mediated decay (NMD), and detection of transcription start sites (TSS) and poly(A) motifs. A total of 1,883,390 were identified. However, isoforms with RT switching events and without poly(A) motif were removed to reach a total of 1,038,414 isoforms. Then, IsoQuant^52^(v3.9.0) was used to quantify the 1,038,414 isoforms, and low expression isoforms (isoforms where expression is ≥ 1 TPM in at least 3 samples) were filtered out to obtain 288,098 transcripts. IsoQuant was re-used to re-quantify and filter out lowly expressed transcripts from this smaller list of 288,098 transcripts and more accurately quantify and obtain a final list of 279,998 isoforms. Finally, isoforms that were not supported by TSS data (e.g. CAGEseq, 5’ end sequencing, H3K4me3 ChIPseq peaks, etc.) or splice junction validation (e.g. short-read RNAseq, presence of repeat elements, or mouse ESCs proteomics data) were filtered out (see **Analysis to support and validate isoform structure and splice junctions** section), leading to a final list of 276,608 isoforms (see **Supplementary Fig. 2** and **Supplementary Data 2**). The coding potential score of detected transcripts was assessed with the Coding-Potential Assessment Tool (CPAT)^129^ (v3.0.5) (**Supplementary Data 2**), which accounts for RNA sequence characteristics such as the ORF size and coverage, using default parameters with the exception of “--min-orf”, which was set at 10. P-values between the distributions of coding probability were calculated by t-test (unpaired and unequal variance).

### Analysis of publicly available bulk and single-cell short-read RNAseq

Raw 5’ RNAseq fastq files from O’Malley *et al*.^130^ were downloaded from Array express E-MTAB-1654 (https://www.ebi.ac.uk/biostudies/ArrayExpress/studies/E-MTAB-1654?query=E-MTAB-1654). Raw paired-ends RNAseq fastq files from Stadhouders *et al*.^73^ were downloaded from GEO accession GSE96608 (https://www.ncbi.nlm.nih.gov/geo/query/acc.cgi?acc=GSE96608). Raw paired-ends RNAseq fastq reads from Qiao *et al*.^13^ were downloaded from GEO accession number GSE138760 (https://www.ncbi.nlm.nih.gov/geo/query/acc.cgi?acc=GSE138760). Raw single-cell RNAseq fastq files from Kim *et al*.^18^ were downloaded from GEO accession number GSE55291 (https://www.ncbi.nlm.nih.gov/geo/query/acc.cgi?acc=GSE55291). Read quality was assessed with fastQC (v0.11.9) (http://www.bioinformatics.babraham.ac.uk/projects/fastqc). Reads were aligned to the mouse mm10/GRCm38 genome using the STAR aligner^131^ (v2.7.11a) with the following parameters: --outSAMtype BAM SortedByCoordinate --outSAMunmapped Within --outSAMstrandField intronMotif --outSAMattributes Standard . Resulting BAM files were indexed and alignment statistics exported with Samtools (v1.17).

For the O’Malley *et al.* dataset, every reprogramming sample was merged using Samtools merge. Coverage for 5’ nucleotides across each read in every sample was calculated with bedtools^132^ (v2.30.0) using genomecov -5 -bg . Both strands were processed separately using the strand + and strand - arguments respectively. 5’ coverages were merged as peaks and coverage was summed per peak with the bedtools merge command using the following parameters: -d 10 -c 4 -o sum. Bed files for plus and minus strand peaks were concatenated and sorted by position. Coverage was converted to counts per million (CPM), and peaks with a coverage below 0.5 CPM were filtered out. The resulting bed file was used in addition to CAGEseq data for SQANTI3 analysis, presented in the **Preparation of a catalog of validated isoform 5’ ends** section.

For the Stadhouders *et al.* and Qiao *et al.* datasets, Salmon^133^ (v1.10.3) was used with the mapping-based mode to quantify expression of a subset of isoforms of interest (available in **Supplementary Data 10**). First, an index was generated using the salmon index command with a fasta file of the isoforms as transcripts and the mouse mm10 chromosomes as decoys. The salmon quant command was used on fastq format reads with the previously generated index and the following arguments: -l U –validateMappings. For Stadhouders *et al.*, TPMs were averaged across reprogramming stages based on principal component analyses presented in Stadhouders *et al*: B cells and Bα as “B cells”, D2 and D4 as “early”, D6 and D8 as “late”. TPM values are available in **Supplementary Data 10** and **Supplementary Data 13**.

For the Kim *et al*. single-cell RNAseq dataset, the salmon quant command was used on fastq format reads with the previously generated index (see the paragraph above) and the following arguments: -l U – validateMappings. To differentiate high-expressor cells from low-expressors and background noise, cells were considered positive for expression of a given isoform if its expression exceeded a per-isoform threshold set at the 33rd percentile. Gene-level read counts were obtained with the featureCounts tool (v2.0.3), from the Subread package, using the exon counting mode^134^. Resulting counts matrices were processed in R (v4.3.3) using Seurat (v5.3.1). Data was processed using Seurat’s recommended SCTransform workflow, as outlined in the official Seurat vignettes. Module scores showing normalized expression of gene sets were calculated using Seurat’s AddModuleScore, using 100 control genes per analyzed gene. Significant differences in module scores were assessed with a Mann-Whitney non-parametric U-test. Log-normalized counts, isoform level TPM values, gene sets and module scores are available as **Supplementary Data 10**.

### Analysis of publicly available ChIPseq data

H3K4me3 ChIPseq and input data from Hussein *et al*.^16^ was re-analyzed. Fastq files were first trimmed with trim_galore (v0.6.10) (https://github.com/FelixKrueger/TrimGalore) using the --paired argument and quality was evaluated with fastQC. Reads were aligned to the NCBI38/mm10 mouse reference using the Bowtie2 alignment algorithm^135^ (v2.5.4) with the pre-set sensitive parameter. Peaks were called using MACS3 (v3.0.3) callpeak using the following arguments: -g mm -f BAMPE -nomodel -B . Resulting bed files from each reprogramming sample were concatenated and sorted by position, then overlapping peaks were merged with bedtools merge.

### LrRNAseq data analysis of gene expression

Raw read counts were obtained using IsoQuant^52^ (v3.9.0) using default parameters for gene level counts. Together, the sequencing runs (45 samples in total; **Supplementary Data 1**) generated 94,777,253 quality control-passed and filtered reads in total, with an average of 2,106,161 reads and 3.8 Gb of data per replicate (**Supplementary Data 1**). EdgeR R-package^136^ (v3.42.4) was then used to normalize the data, calculate RNA abundance at the gene level (as CPM), and perform statistical analysis. Briefly, a common biological coefficient of variation (BCV) and dispersion (variance) were estimated based on a negative binomial distribution model. This estimated dispersion value was incorporated into the final EdgeR analysis for differential gene expression, and the generalized linear model (GLM) likelihood ratio test was used for statistics, as described in EdgeR user guide. Genes with CPM ≥ 1 in at least one sample were considered (n=21,757 genes) and listed in **Supplementary Data 1**. Differential gene expression (DEGs) was established as genes with a fold change ≥ 2 (up-regulation) or ≤ 0.5 (down-regulation) and adjusted p-value (FDR) ≤ 0.05. Gene expression plots were generated using Microsoft Excel by averaging the log10(CPM+1 expression values) of the different replicates of a given timepoint for the different gene lists and representing the SEM as a shaded contour (gene lists are listed in **Supplementary Data 1** and **Supplementary Data 7**). Heat map expression of manually curated alternative splicing regulators was generated using the minimum and maximum log10(CPM+1) expression values of these genes on Microsoft Excel (**Supplementary Data 7**).

Sample clustering was performed on top varying genes: Briefly, the variance of expression was calculated for each gene. All genes with a variance across samples superior to 10% above the average variance were considered as top-varying genes (n=489 genes). Gene expression was normalized as z-scores, and then distance matrices were calculated with the *dist* R function, using the Euclidean distance. Clustering was then performed using the *hclust* R function using the complete method.

### Analysis of isoform level expression and DEI-based isoform switching

Isoform level expression was quantified using IsoQuant^52^ (v3.9.0) using default parameters. Low expression isoforms (isoforms where expression is ≥ 1 TPM in at least 3 samples) and isoforms that were not supported by either TSS or splice junction data were further filtered to obtain a total of 276,608 isoforms (**Supplementary Data 2**). CPM values were converted to TPM values manually. EdgeR R-package^136^ (v3.42.4) was also used on the isoform counts values to perform statistical analysis, as described in the **LrRNAseq data analysis of gene expression** section above. Differentially expressed isoforms (DEIs) were established similar to DEGs, fold change ≥ 2 or ≤ 0.5, and FDR ≤ 0.05 when comparing reprogramming phases, but this time TPM values were used as expression values. This represented 198,247 isoforms that were selected for further analysis (**Supplementary Fig. 2** and **Supplementary Data 9**). For downstream analyses, only isoforms that were differentially expressed between D0 and the middle phase of reprogramming were selected. Finally, changes in isoform usage that were associated to the comparison between NPCs (D0) and D2 were kept only if they were not associated with the comparison between NPCs and wild-type NPCs placed in reprogramming medium for 2 days (FBSL) (i.e. with no reprogramming transgenes induction). This was done in order to select only differential isoform usage attributed to the effect of reprogramming transgenes expression and not due to media change. Isoforms were only selected if they were expressed in 40% of the samples. We obtained 35,262 isoforms that were subsequently used in Microsoft Excel to generate plots compiling the average expression in TPM of four categories (up-, down-, up-then down-, or down- then up-regulated) of DEIs between D0 and the middle phase of reprogramming (**Supplementary Data 9**). The differential usage of isoforms from a gene between D0 and the middle phase of reprogramming was used to identify genes that undergo isoform switch (IS) during this period according to an in-house matrix defined with Microsoft Excel (**Supplementary Data 9**). Observed percentages of genes undergoing IS among each biotype were calculated based on the total count of genes undergoing IS between D0 and the middle phase of reprogramming. Expected percentages were based on the total count of genes with DEIs during reprogramming, and p-values were calculated by Chi-squared test. Gene Ontology enrichment analysis of genes associated with at least one significant IS event per comparison was performed using DAVID^121^ (v2022q3). Annotations were analyzed with a cut-off of FDR ≤ 0.1 for significant enrichment (**Supplementary Data 9**). Validation of *Snhg26* isoform switching between the NPC and middle phase samples was performed by PCR-amplifying from previous PCR-amplified libraries for 34 PCR cycles with ONT’s forward cPRM and *Snhg26* isoforms’ specific reverse primers (primer sequences are listed in **Supplementary Data 11**). Resulting amplicons were prepared for lrRNAseq and processed as in the **Library preparation and lrRNAseq, processing and alignment** section. For each sample, reads were counted manually and averaged before dividing by the total number of mapped reads in the sample in order to get the expression level in CPM for both *Snhg26* isoforms (fastq reads archive is available in **Supplementary Data 12**).

### Analysis of repetitive elements within splice junctions

First, a GTF file of every repetitive genomic sequence was created: 30nt sliding windows with 10nt steps were generated across the mouse mm10/GRCm38 genome with bedtools makewindows. Then, bedtools nuc was used to identify intervals where nucleotide composition was at least 85% GC, 85% AT or 85% of a single base. Overlapping repetitive windows were merged. Repetitive regions were merged to the annotated mm10 RepeatMasker .out file (available from UCSC: https://hgdownload.soe.ucsc.edu/goldenPath/mm10/bigZips/). The resulting repetitive regions that overlapped exons were extracted with bedtools intersect using our filtered transcriptome GTF file. The effect of repetitive sequences on short-read RNA sequencing coverage was assessed on male and female mouse ESC datasets^137,138^ (GEO accession: GSE67259, https://www.ncbi.nlm.nih.gov/geo/query/acc.cgi?acc=GSE67259) using Deeptools’ bamCoverage with BPM normalization, ComputeMatrix using the exonic repetitive sequences bed file as regions file and the -b 250 -a 250 --binSize 1 --regionBodyLength 50 arguments and plotProfile using the --kmeans 1 --plotType se arguments (**Supplementary Fig. 3g**). A repetitive regions bed file is available as **Supplementary Data 2.**

### Preparation of a catalog of validated isoform 5’ ends

A bed file was generated from the refTSS version 4 transcript 5’ ends dataset^139^, which includes the FANTOM5 annotations, the TE-TSS database^140^, 5’ RNA sequencing of reprogramming cells from O’Malley *et al.*^130^ and CAGEseq data from iPSCs^141^. To account for lower sequencing depths of the short-read sequencing within repetitive regions, we also included the repetitive regions obtained in the **Analysis of repetitive elements within splice junctions** section. The centre of each region was added as columns 7 and 8 to allow for calculation of distance to TSS by SQANTI3. The 5’ ends bed file is available as **Supplementary Data 1**.

### Analysis to support and validate isoform structure and splice junctions

Following isoform filtering (see **Mouse transcriptome assembly and isoform classification** section), isoforms were re-analyzed with SQANTI3 (v5.2.2) using the sqanti3_qc.py script, but this time included short-read and TSS support. Short-read datasets presented in **Supplementary Data 3** were downloaded as fastq files and supplied for isoform validation with the --short_reads argument to assess splice junctions support, short-read expression and TSS-ratio. The --CAGE_peak argument was used with the bed file generated in the **Preparation of a catalog of validated isoform 5’ ends** section. The --polyA_motif_list argument was used with the default mouse poly(A) motifs provided in the SQANTI3 software.

To assess the presence of repetitive sequences within non-supported splice junctions, a bed file was generated from the SQANTI3 junctions.txt file corresponding to the 5’ and 3’ splice sites of each junction without short-reads support, surrounded by 150nt upstream of the 5’ and downstream of the 3’ splice site (150nt being the read length of most short-read datasets used for validation) (**Supplementary Fig. 3f**). The junctions bed file was intersected with the repetitive sequences file described in the **Analysis of repetitive elements within splice junctions** section. Junctions where at least one splice site overlapped a repetitive sequence were considered supported.

For coding isoforms with junctions remaining non-supported, the encoded protein sequences were compared to experimentally detected peptides obtained by mass spectrometry (see **Sample preparation for total proteome quantification**, **Mass spectrometry analysis**, and **DIA data analysis** sections below). First, isoforms with the same protein sequence as supported isoforms were excluded. The remaining isoforms for which peptides were detected were considered as supported.

Isoforms without TSS support by CAGEseq or comparable 5’ sequencing datasets were assessed for their TSS-ratio (i.e. ratio of reads after/before the TSS). Based on thresholds established in Pardo-Palacios *et al.*^55^, isoforms with a TSS-ratio superior to 1.5 were considered to be supported. TSS were also compared to H3K4me3 peaks established in the **Analysis of publicly available ChIPseq data** section. A TSS lying within a H3K4me3 peak or within 100bp from the centre of the peak was considered supported. A classification file detailing support for all filtered isoforms is available as **Supplementary Data 2**.

### Functional and structural annotation of isoforms

GFF3 files generated from SQANTI3’s IsoAnnotLite pipeline were converted to a SQANTI3-like classification file (**Supplementary Data 8**). Untranslated region lengths of isoforms from genes subject to alternative splicing between reprogramming stages were compared with a non-parametric Kruskal Wallis test followed by Dunn’s *post hoc* test. Percentage of non-coding or NMD isoforms from coding genes, or isoforms with PFAM domains were calculated per sample. Percentage of isoforms containing each annotation type was calculated per sample, and significant differences across reprogramming stages were assessed with an ANOVA followed by Tukey’s *post hoc* test. Annotations are available in **Supplementary Data 2**.

### Comparison to other transcriptome-building tools

Transcriptome was built using three orthogonal methods: Bambu^51^ (v3.4.1) was used in R (v4.3.3) with a Novel Discovery Rate of 0.5 and the stranded = TRUE argument. IsoQuant^52^ (v3.9.0) was used with the following arguments: --complete_genedb --sqanti_output --count_exons --check_canonical --report_canonical all --splice_correction_strategy conservative_ont --matching_strategy default --model_construction_strategy all --data_type nanopore --report_novel_unspliced true --polya_requirement never . Finally, StringTie^53^ (v3.0.0) was run two ways: 1) with reference (Mus_musculus.GRCm38.99.gtf from Ensembl) and 2) *de novo*, both using the -s 1 -f 0 -L -v -G arguments. The resulting GTF files were then merged using StringTie with the --merge -i -f 0 -T 0 -F 0 -g 0 arguments.

Resulting GTF files were filtered to remove isoforms without strand information. Then, corresponding isoforms were identified using the sqanti3_qc.py script. The four generated GTF files were used sequentially as references: (i) the GTF generated using the method described in the **Mouse transcriptome assembly and isoform classification** section, (ii) the Bambu GTF, (iii) the IsoQuant GTF, and (iv) the StringTie GTF. Each GTF file was compared against the others in a pairwise manner. Isoforms classified as Full Splice Matches were considered corresponding isoforms between two orthogonal methods. A list of corresponding isoforms is available as **Supplementary Data 3**. GTF files from orthogonal methods are available as **Supplementary Data 6**.

### Analysis of alternative splicing

The occurrence of local alternative splicing (AS) and transcript events during reprogramming was assessed using SUPPA2^142^ (v2.43) and the “suppa.py generateEvents” script with parameters “-f ioe -e SE SS MX RI FL” for AS events and parameter “-f ioi” for transcript events. A GTF file containing the expression filtered isoforms (i.e. 279,998 isoforms) was used as input for these analyses. To calculate the percent spliced in (PSI) values for each isoform and seven types of alternative splicing events, “suppa.py psiPerEvent” script was used with their respective events files (i.e. “.ioi” and “.ioe”) and the TPM reads values from IsoQuant quantification above. Only the AS variations that had an adjusted p-value ≤ 0.05 in at least one reprogramming phase comparison were selected, reducing the list of 381,230 AS events to 140,645 AS events, and isoforms to 33,657 (events and isoforms are listed in **Supplementary Data 7**). Differential splicing between reprogramming phases was then established as isoforms with a ΔPSI ≥ 0.1 (more inclusion of an AS event) or ΔPSI ≤ -0.1 (less inclusion of an AS event) and an adjusted p-value ≤ 0.05. Total number of more or less included AS events for each type of AS was represented using Microsoft Excel. Differentially spliced isoforms between reprogramming phases were established as isoforms with a ΔPSI ≥ 0.1 (more abundant isoform) or ΔPSI ≤ -0.1 (less abundant isoform) and an adjusted p-value ≤ 0.05. Total number of alternatively spliced transcripts with more or less relative abundance was represented using Microsoft Excel. Similar to isoform usage calculations, AS and transcript events were kept only if they were associated to changes due mainly to reprogramming, meaning event changes occurring due to media change as identified by comparing NPCs to FBSL were removed.

Overlap between genes with significant AS events (although not necessarily the same AS events) detected in this dataset (i.e. 10,460 genes) and other reprogramming datasets^32,35^ was presented as a Venn diagram using InteractiVenn^143^ (gene lists are presented in **Supplementary Data 7**). GO-term enrichment on genes commonly spliced was assessed using DAVID^121^ (v2022q3) and FDR ≤ 0.1 (**Supplementary Data 7**).

FYN target genes were selected using the CRISPR screen, RNAi screen and Predictability datasets from the DepMap Public 25Q3 Release dataset through the DepMap portal^69^ (https://depmap.org/portal). Their expression was averaged depending on whether they correlated negatively or positively with *Fyn* expression in the CRISPR screen, based on a correlation threshold ≥ 0.3 or ≤ -0.3 (gene lists are listed in **Supplementary Data 7**).

### Visualization of gene loci and alternatively spliced isoforms

Validations of gene loci, alternative splicing events, isoform abundance and isoform switching were performed using the Integrative Genomics Viewer (IGV)^144^ (v2.16.2) tracks.

### Identification of candidate lncRNAs

A first list of genes for which alternatively spliced isoforms were significantly more or less relatively abundant (i.e. IS analysis based on ΔPSI events method, see **Analysis of alternative splicing** section) between D0 and the middle phase of reprogramming, removing those occurring when comparing NPCs to FBSL samples, which represent changes due to media, was created (i.e. 12,428 genes). A second list of genes that undergo isoform switching (IS analysis based on DEIs method, see **Analysis of isoform level expression and isoform switching** section), between D0 and the middle phase of reprogramming, and which are not due to media change, was generated (i.e. 2,338 genes). The two lists are presented in **Supplementary Data 10**. Overlap between these two lists (i.e. 1,736 genes) was presented as a Venn diagram using Microsoft Excel. LncRNAs from this overlap were cross-referenced with lncRNAs detected in Hussein *et al.*^16,42^.

Expression of candidate lncRNAs from publicly available datasets (all listed in **Supplementary Data 11**) were represented with SEM using Microsoft Excel or GraphPad Prism (v7.04). P-values between the frequencies were calculated using t-test (unpaired and unequal variance).

### Sample preparation for total proteome quantification

25μg of cell samples were processed per sample type. Each sample was reduced with DTT at 10mM for 30 minutes at 56°C and alkylated with iodoacetamide at 30mM for 45 minutes at room temperature in the dark. Each sample was then brought up to a final volume of 50L. 6.25μL of MagReSyn HILIC beads (Resin Biosciences; Gauteng, South Africa) per sample were transferred to a microcentrifuge tube and washed 2 times with 200μL Equilibration buffer (15% ACN in 100mM Ammonium acetate, pH 4.5). Each 12.5μL initial bead slurry was moved into a new microcentrifuge tube in equilibration buffer. 50μL Binding buffer (30% ACN in 200mM Ammonium acetate, pH 4.5) was added to 50μL protein solution for a final volume of 100μL. Equilibration buffer was removed from the beads and 100μL protein solution was added and mixed at 1,200 rpm for 30 minutes at room temperature. Beads were washed twice with 200μL Wash buffer (95% ACN), mixed for 1 minute at 1,200rpm. All wash buffer was removed and beads resuspended in 100μL digestion buffer (50mM Tris-HCl, pH 8.0) with 1μg trypsin/Lys-C. Protein was digested for 1 hour at 46°C with mixing (1,200 rpm). Peptides were collected and acidified to 2% formic acid. Peptides were dried down and stored at -40°C until mass spectrometry (MS) acquisition.

### Mass spectrometry analysis

For data-independent acquisition (DIA) liquid chromatography-Tandem mass spectrometry (LC-MS/MS), 500ng protein equivalent of digested peptides were analyzed using a nano-high-performance liquid chromatography (HPLC) coupled to MS. Samples were separated on an Aurora Elite TS column (IonOpticks Pty Ltd. #AUR3-15075C18-TS). The sample in 5% formic acid was trap (300µm i.d., 0.5cm length, #174500) loaded at 800 bar, onto a 75µm i.d. x 15cm nano-spray emitter (packed with 1.7µm C18 beads) heated at 50°C with the TS Interface. Peptides were eluted from the column with an acetonitrile gradient generated by an Vanquish Neo UHPLC System (ThermoFisher Scientific), and analyzed on an Orbitrap^TM^ Astral^TM^. The gradient was delivered at 800nL/min from 0.8% acetonitrile with 0.1% formic acid to 3.2% acetonitrile with 0.1% formic acid over 1 minute, 3.2% to 6.4% acetonitrile with 0.1% formic acid over 1 minute, and 6.4% to 35.2% acetonitrile with 0.1% formic acid over 19 minutes. This was followed by a column wash of 76% acetonitrile with 0.1% formic acid over 2 minutes. The LC method ends with an equilibration of 2 column volumes at the same combined control as loading. The total DIA protocol is 23 minutes. Advanced peak determination was turned on with expected LC peak width of 12 seconds. FAIMS was used with -50V compensation voltage. MS1 scan was from 380-980m/z with an orbitrap resolution of 240,000, normalized AGC target of 500%, and 50ms maximum injection time in profile mode. DIA windows were set to auto with 4m/z window sizes with 0 overlap, in the precursor range of 380-980m/z. HCD collision energy was set to 25% using the Astral detector. The scan range was 150-2000m/z with RF lens set to 50%, AGC target to 500%, and maximum injection time of 3 ms.

### DIA data analysis

Spectronaut v20 directDIA+ (Biognosis AG; Switzerland) BGS workflow was used to search the data with the Spectronaut generated mouse spectral library (Mouse_Uniprot_2023_06_13) supplemented with the fasta file derived from the SQANTI3 output files, as defined in the **Mouse transcriptome assembly and isoform classification** section. Parameters for the search were default, normalization was on with runwise imputation. Differential abundance testing used was unpaired t-test (data is available in **Supplementary Data 3**).

## Inclusion & Ethics statement

All collaborators of this study that fulfilled Nature Portfolio journals’ criteria for authorship have been included as authors, as their participation was essential for the design and implementation of the study. Roles and responsibilities were agreed among collaborators ahead of the research. This research would not have been severely restricted or prohibited in the setting of the researchers.

## Reporting Summary

Further information on research design is available in the Nature Portfolio Reporting Summary linked to this article.

## Data Availability

Datasets from other referenced works can be accessed either in the NCBI Gene Expression Omnibus (GEO) repository using their GSE numbers or in the database of the EMBL’s European Bioinformatics Institute using their E-MTAB numbers (all reference datasets and processed datasets with links and descriptions are summarized in **Supplementary Data 3**, **10** and **11**). Long-read RNA sequencing raw data, filtered aligned reads and processed data, including mouse transcriptome annotation and expression level matrices, from this study are publicly available and have been deposited in the GEO database under the accession codes: GSE282319 (mouse reprogramming data, https://www.ncbi.nlm.nih.gov/geo/query/acc.cgi?acc=GSE282319) and GSE317557 (human knock-down data, https://www.ncbi.nlm.nih.gov/geo/query/acc.cgi?acc=GSE317557). All mass spectrometry files used in this study were deposited in MassIVE (http://massive.ucsd.edu) and assigned the identifier MSV000100321 (https://massive.ucsd.edu/ProteoSAFe/dataset.jsp?task=94497fd1dc824ed0a471a6638f159439). All cell lines and biological materials generated in this study are available from the corresponding author upon direct request.

## Acknowledgements

We thank Dr. Ian Roger and Dr. Sophie Mockly for advice and reading of the manuscript. Gaëlle Bourriquen assisted with the mouse CRISPRi sgRNA design. We also acknowledge the contribution of the Model Production and Pathology Cores at The Centre for Phenogenomics for technical support in generating chimeras and processing teratomas. We thank the Network Biology Collaborative Centre Proteomics Facility (RRID: SCR_025375) at the Lunenfeld-Tanenbaum Research Institute for total proteome analysis.

## Funding Statement

This work was funded by the Canadian Institutes of Health Research (CIHR) to S.M.I.H. (PJT-378019) and supported by the Fonds de recherche du Québec (FRQ) through the Research Centre Grant for the CHU de Québec-Université Laval Research Centre (30641). V.F. is a recipient of a training award of the Fonds de recherche du Québec - Santé (FRQS) https://doi.org/10.69777/291836. G.K. is a recipient of training awards form the Natural Sciences and Engineering Research Council of Canada (NSERC) and the FRQS https://doi.org/10.69777/307170. V.W. is a recipient of a training award of the Fonds de recherche du Québec - Nature et Technologies (FRQNT) https://doi.org/10.69777/271653. S.M.I.H. and J.P.L. are Junior 2 and Senior Research Scholars of the FRQS https://doi.org/10.69777/310750 and https://doi.org/10.69777/380310, respectively. The Network Biology Collaborative Centre Proteomics Facility is supported by the Canada Foundation for Innovation and the Ontario Government.

## Author Contributions Statement

V.F., J.P.L., Y.Y., J.W., and S.M.I.H. conceived of and designed the study. V.F., G.K., R.H., V.W., M.P., N.Y., M.G., E.I.J.L., J.L., and J.P.L. developed and performed the experiments or collected data. V.F., G.K., J.P.L, and S.M.I.H. designed and performed the computational and statistical analyses. V.F., G.K., R.H., V.W., M.G., E.I.J.L., V.M., J.P.L., Y.Y, J.W., and S.M.I.H. wrote, reviewed and edited the manuscript.

## Competing Interests Statement

The authors declare no competing interests.

## Notes

### Competing Interest Statement

The authors have declared no competing interest.

### Summary of Updates

Some Supplementary figures were added and the main figures were updated to reflect extra filtering of isoforms.

