## Supplementary Figures and Note for "Long-read RNA sequencing identifies non-coding isoform switching as a regulator of cell fate"

#### Supplementary Information

### Supplementary Figure 1

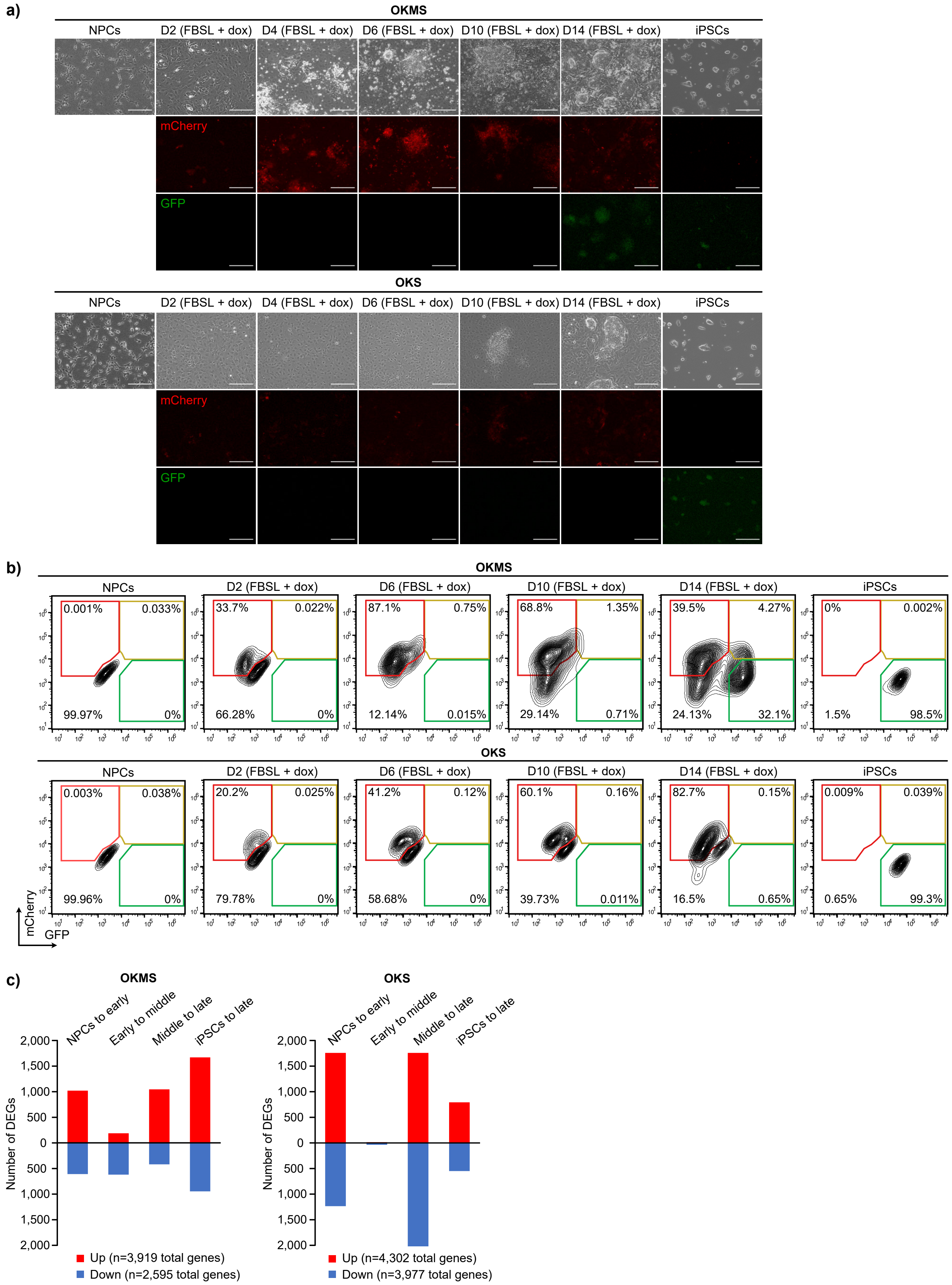

**Supplementary Figure 1: *mCherry* and *GFP* reporter genes to follow reprogramming factors and endogenous *Oct4* expression**

**a)** Representative images in brightfield or fluorescence microscopy of the NPCs, reprogramming cells at different timepoints, and final iPSCs (D20 2i-LIF) in the OKMS and OKS reprogramming systems. Scalebars = 50 µm.

**b)** Representative flow cytometry data allowing quantification of the percentages of reprogramming cells (mCherry+) and reprogrammed iPSCs (GFP+) throughout reprogramming with OKMS or OKS.

**c)** Number of up- or down-regulated genes between the early, middle and late phases of OKMS and OKS reprogramming. Significance based on FDR $\leq 0.05$ (calculated by Benjamini-Hochberg method) and log2(fold change) $\geq 2$ for up-regulated genes or $\leq 0.5$ for down-regulated genes.

### Supplementary Figure 2

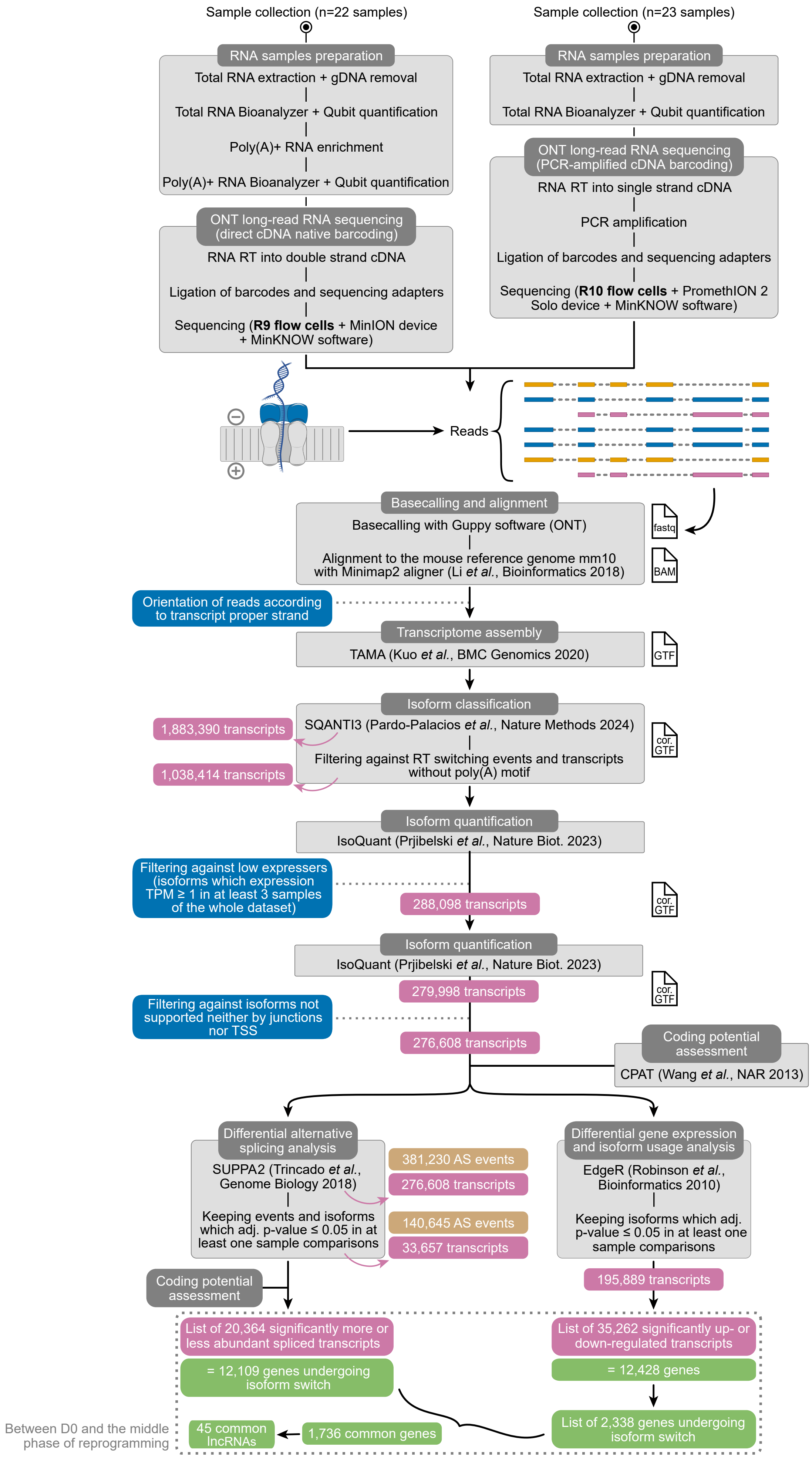

**Supplementary Figure 2: Pipeline of the lrRNAseq dataset generation and analysis**

Schematic overview of the experimental and analysis pipeline used to generate full-length transcript annotations of OKMS and OKS reprogramming samples. gDNA: genomic DNA; RT: Reverse Transcription; cDNA: complementary DNA; PCR: Polymerase Chain Reaction; ONT: Oxford Nanopore Technologies; cor. GTF: corrected GTF; adj. p-value: adjusted p-value.

### Supplementary Figure 3

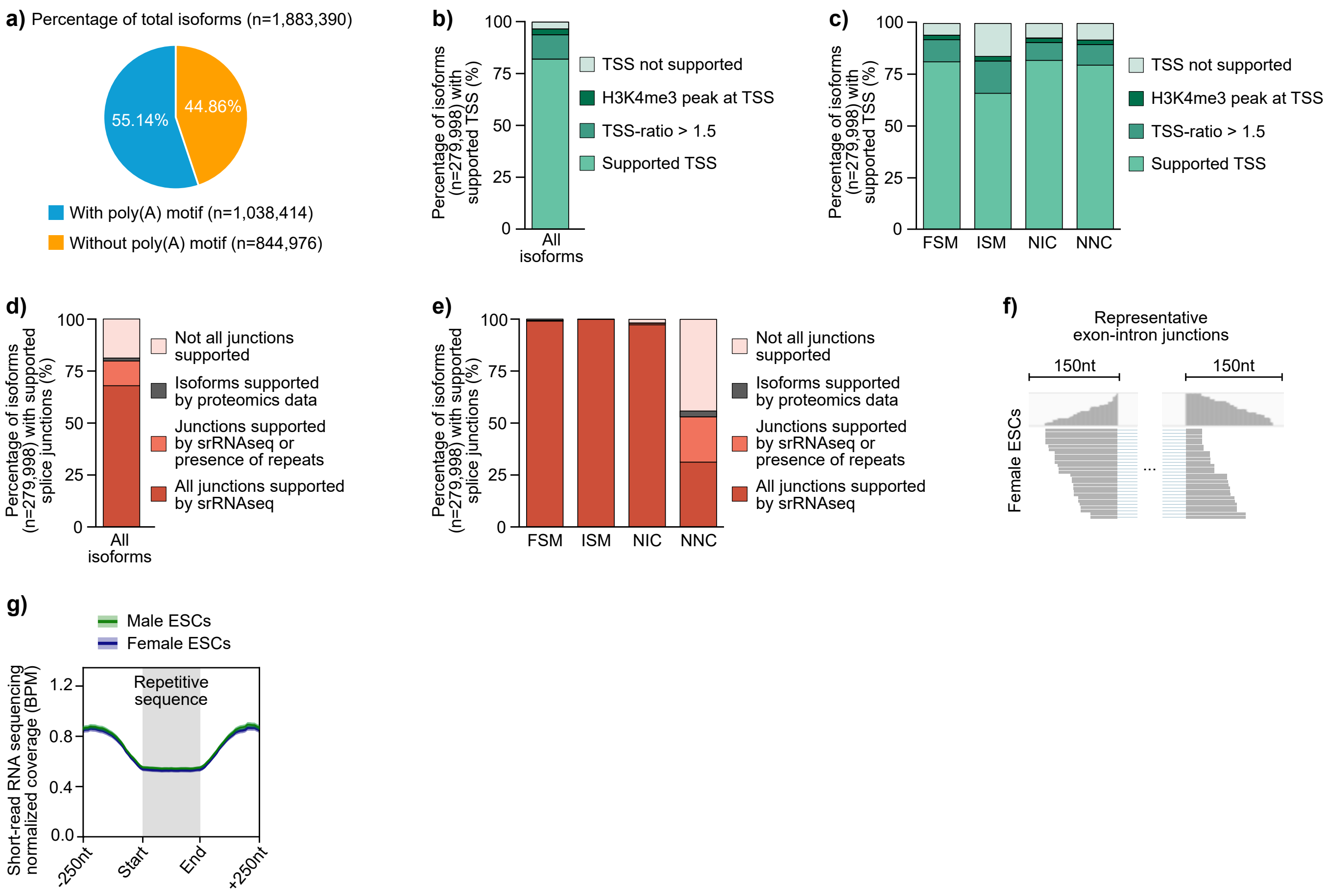

**Supplementary Figure 3: Filtering and validation of detected isoforms**

**a)** Proportion of isoforms with or without a poly(A) motif among the total detected transcripts of the generated dataset.

**b)** Percentage of isoforms with transcription start sites supported by CAGEseq data or orthogonal approaches across all isoforms or **c)** within isoform structural categories. TSS-ratio: Ratio of read coverage downstream TSS to upstream TSS.

**d)** Percentage of isoforms with supported splice junctions across all isoforms or **e)** within isoform structural categories. srRNAseq: short-read RNA sequencing.

**f)** Representative view of short-read RNAseq split reads covering an intron on a genome browser. Boxes represent reads mapped to exons, lines represent read splitting over introns.

**g)** Short-read RNAseq coverage at and around exonic repetitive sequences in ESCs, normalized as bins-per-million (BPM). Lines represent average expression and shaded areas represent standard error.

### Supplementary Figure 4

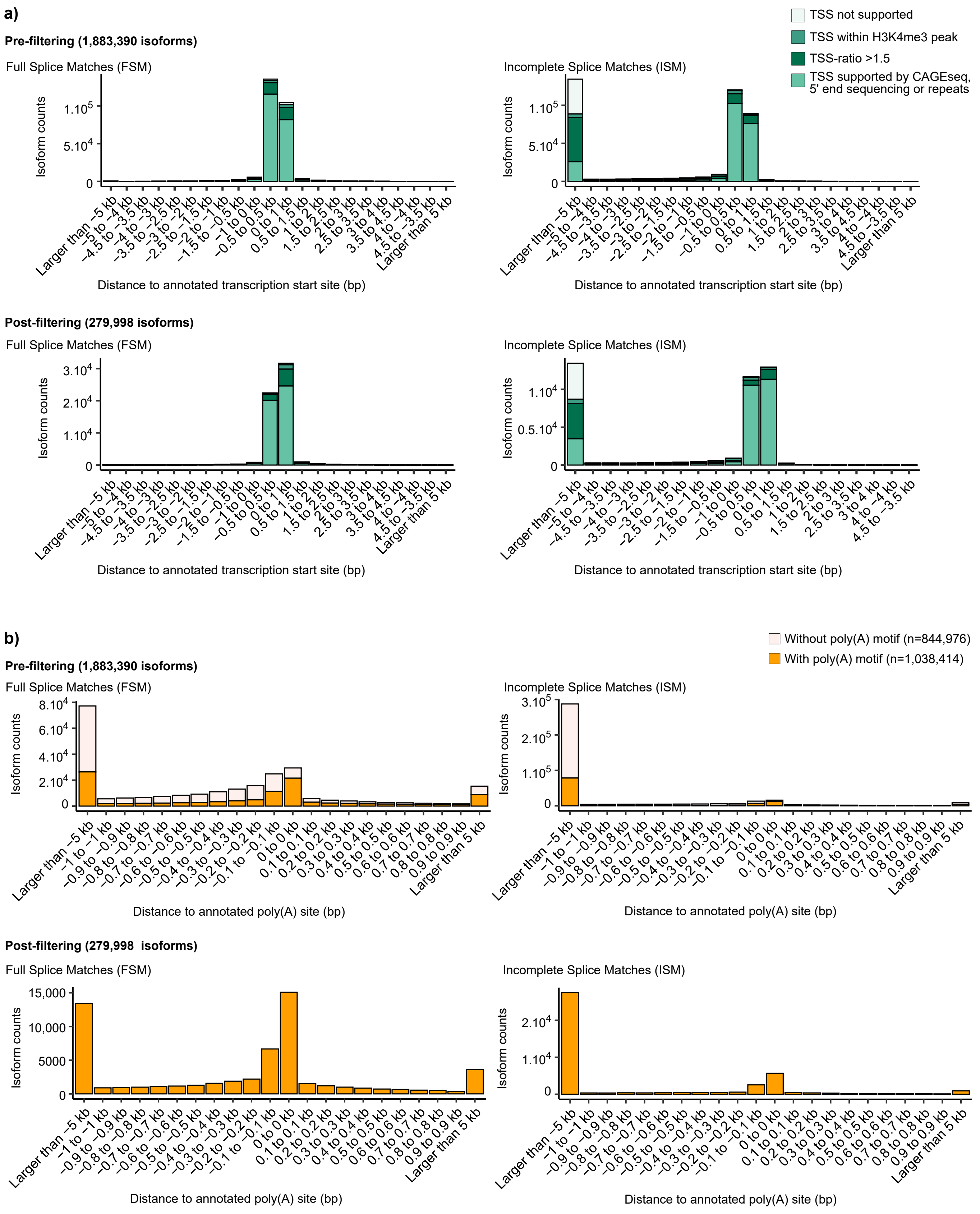

**Supplementary Figure 4: Distance from annotated isoform ends of the detected and filtered isoform datasets**

**a)** Distance to annotated transcription start site and TSS support status, or **b)** distance to annotated poly(A) site and presence of poly(A) signal within isoforms for either the total transcriptome (available as processed data for GEO Series GSE282319), or the filtered transcriptome datasets. kb: kilobases; bp: base pair.

### Supplementary Figure 5

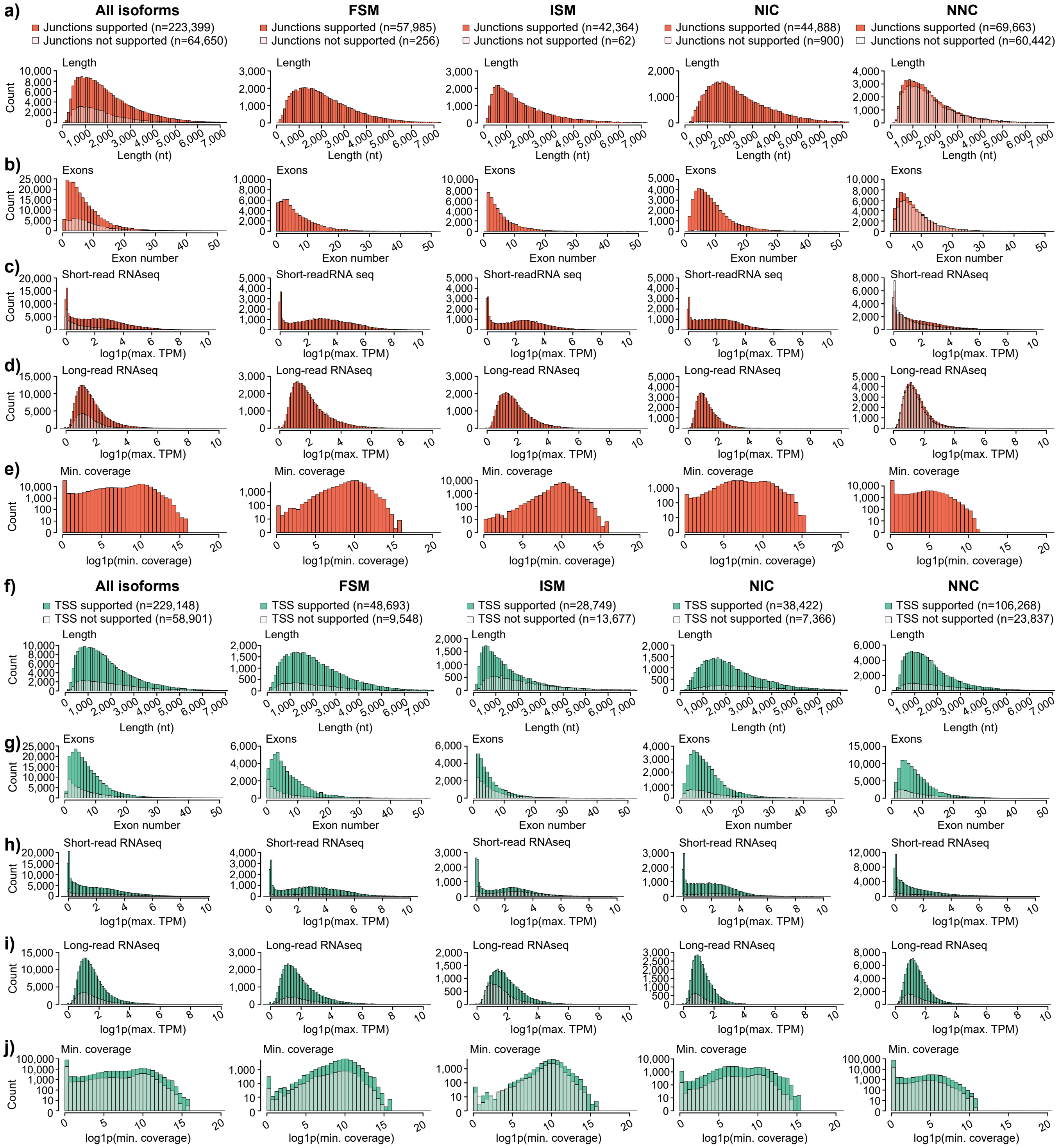

**Supplementary Figure 5: Isoform quality metrics for supported and non-supported isoforms**

**a)** to **e)** Isoform quality metrics for isoforms with or without splice junction support by short-read RNAseq, and **f)** to **j)** with or without CAGEseq support. Histograms represent **a)** and **f)** isoform lengths, **b)** and **g)** exon numbers, **c)** and **h)** expression in short-read RNAseq public datasets[130,139-141], **d)** and **i)** expression in our long-read RNAseq dataset, and **e)** and **j)** coverage at the lowest-covered splice junction (i.e. minimal (min.) coverage), shown with a log scale.

### Supplementary Figure 6

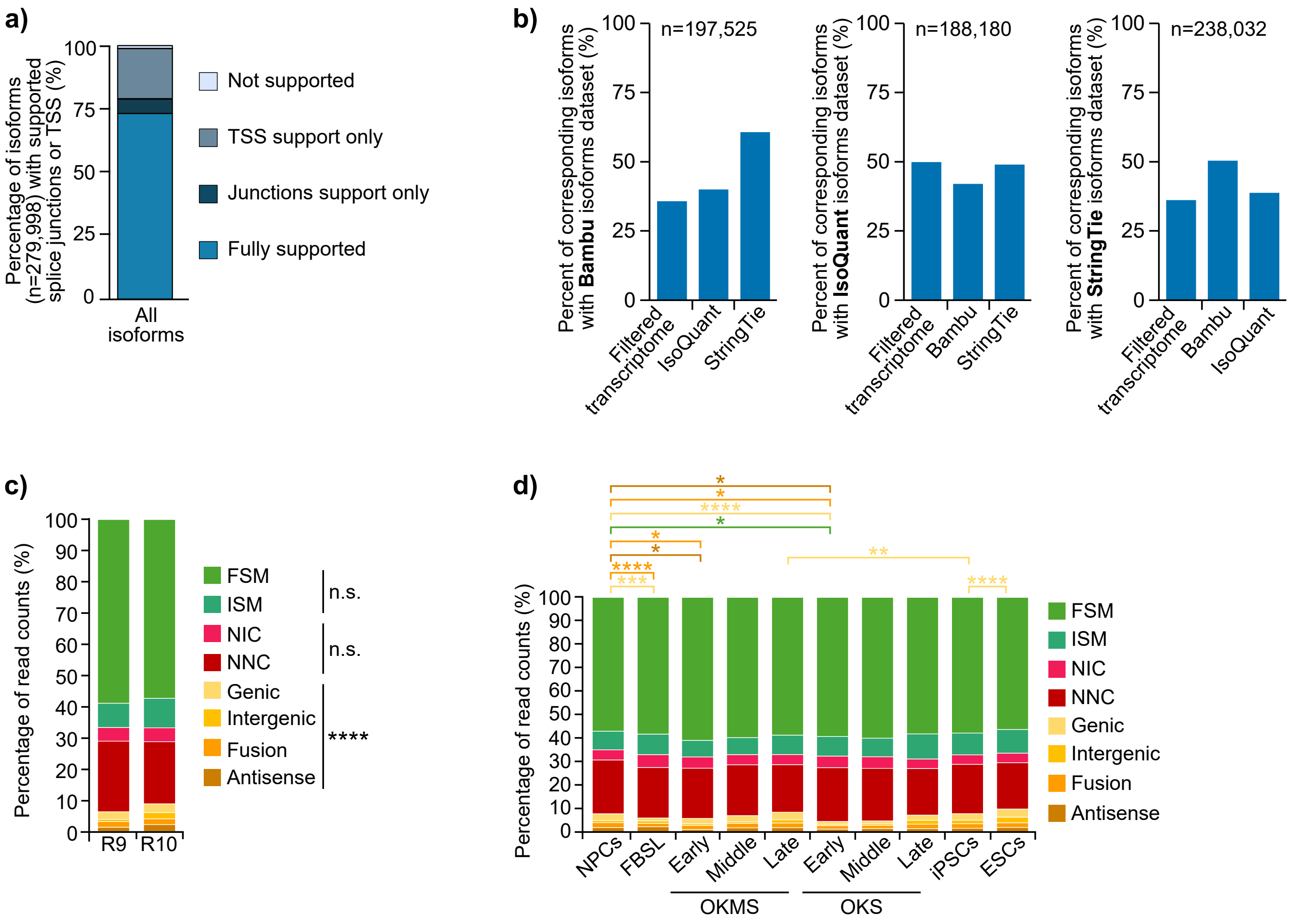

**Supplementary Figure 6: Proportion of isoform categories during reprogramming**

**a)** Percentage of isoforms with supported splice junctions and/or TSS across all isoforms.

**b)** Percentages of corresponding isoforms obtained by orthogonal transcriptome building tools (Bambu[51], IsoQuant[52] or StringTie[53]) when compared with the filtered transcriptome generated in this study and with each other.

**c)** Proportions of isoform structural categories detected in the filtered dataset, separated by flow cell technology (R9 n=22 samples, R10 n=23 samples), or **d)** separated by reprogramming phase (NPCs (n=8), FBSL (n=3), early (n=2), middle (n=4) or late (n=4 for OKMS and n=2 for OKS) phases of reprogramming, iPSCs (n=5) and ESCs (n=9)). P-values were calculated using two-sided t-test with unequal variance. n.s. $\geq 0.05$, * $< 0.05$, ** $< 0.01$, *** $< 0.001$, **** $< 0.0001$.

### Supplementary Figure 7

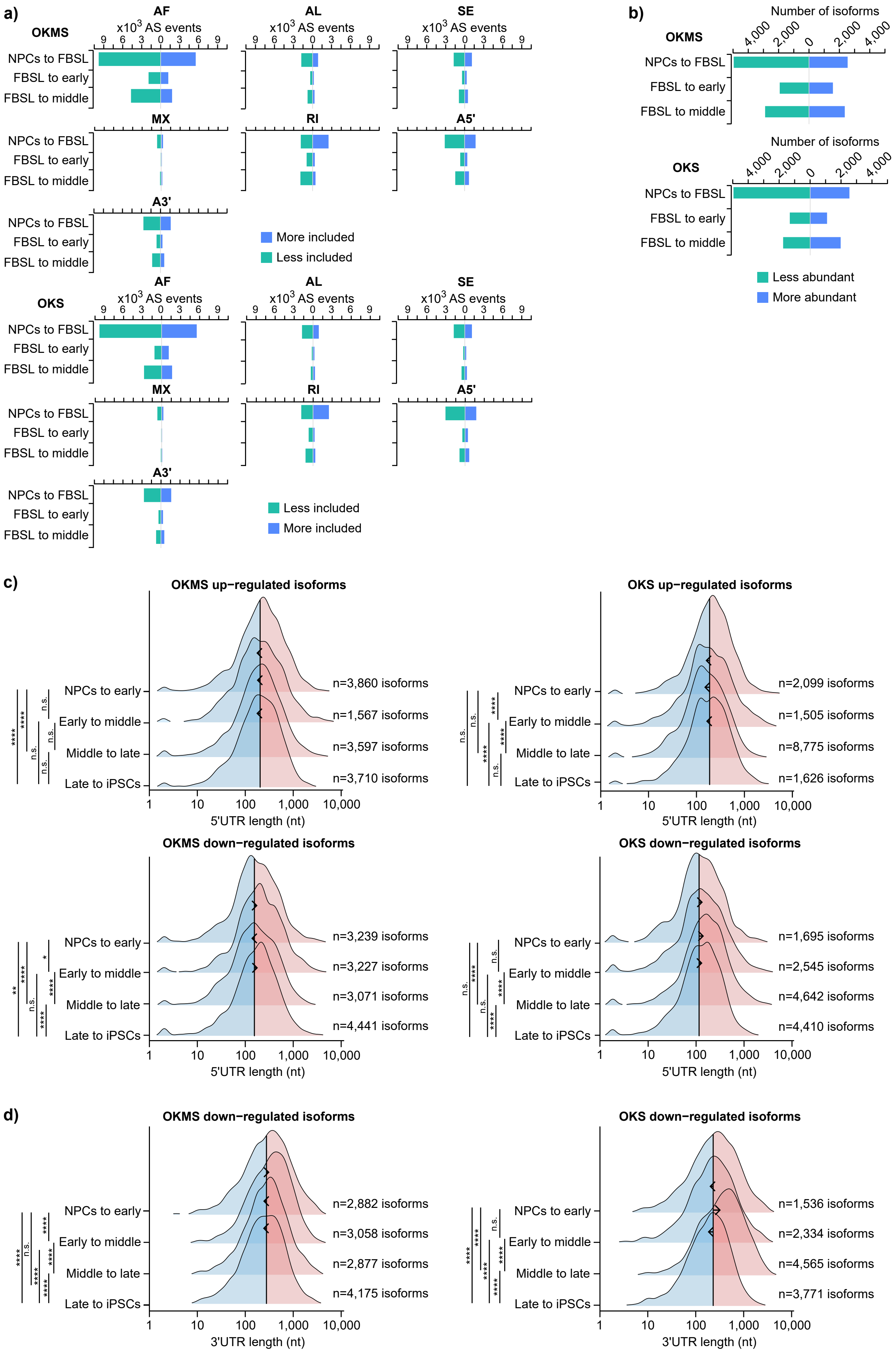

**Supplementary Figure 7: Alternative splicing variations during reprogramming**

**a)** Splicing events-based quantification of alternative splicing significantly more or less included, or **b)** isoform-based quantification of alternative splicing significantly more or less abundant, during OKMS and OKS reprogramming phase transitions (NPCs (n=8), early (n=2), middle (n=4), late (n=4 for OKMS and n=2 for OKS) phases of reprogramming, iPSCs (n=5)). Significance based on a $\Delta PSI \geq 0.1$ (more included) or $\Delta PSI \leq -0.1$ (less included) and an adjusted p-value $\leq 0.05$ (one-sided unequal variance). The data is the same in OKMS and OKS for NPCs to FBSL and ESCs to iPSCs comparisons because those samples were in common for OKMS and OKS.

**c)** Distribution of 5'UTR and **d)** 3'UTR lengths (log scale) in differentially expressed isoforms from genes subject to alternative splicing during OKMS and OKS reprogramming phase transitions (NPCs (n=8), early (n=2), middle (n=4), late (n=4 for OKMS and n=2 for OKS) phases of reprogramming, iPSCs (n=5)). Lines represent means of the NPCs to early datasets. Arrow points show change in distribution means in each transition. P-values were calculated by ordinary one-way ANOVA with Tukey HSD *post hoc* test. n.s. ≥ 0.05, * < 0.05, ** < 0.01, **** < 0.0001.

### Supplementary Figure 8

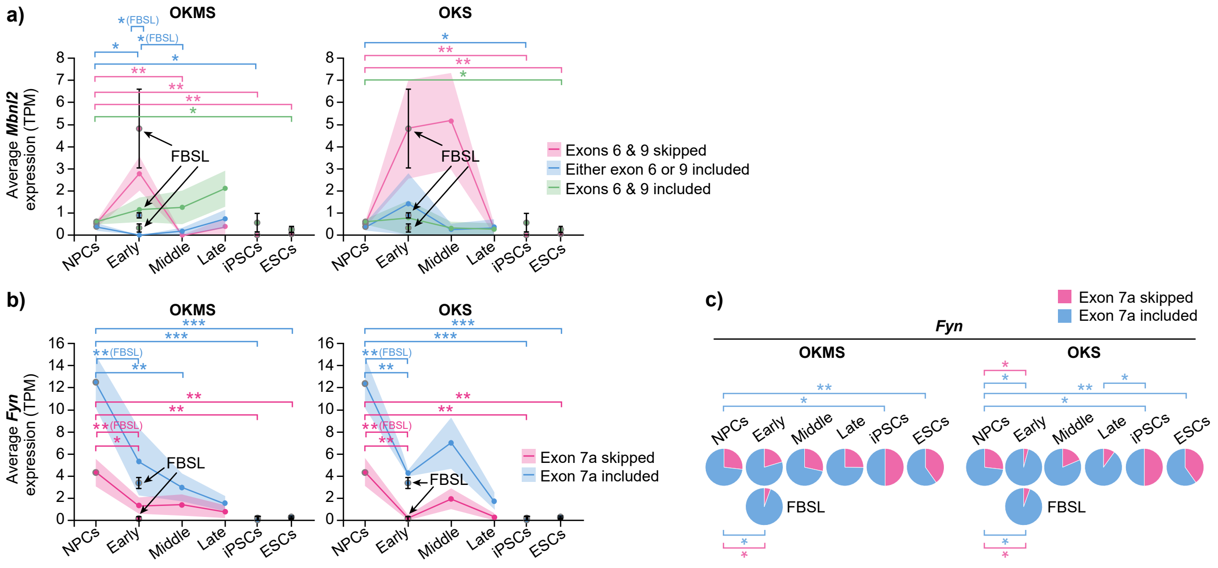

**Supplementary Figure 8: *Mbnl2* and *Fyn* isoforms expression**

**a)** Expression of *Mbnl2* or **b)** *Fyn* isoforms in NPCs (n=8), FBSL (n=3), during early (n=2), middle (n=4) or late (n=4 for OKMS and n=2 for OKS) phases of reprogramming, or in the iPSCs (n=5) and ESCs (n=9) of the dataset. Dots and line represent average and shaded area represent the SEM for reprogramming samples. For the NPCs, FBSL, iPSCs and ESCs samples, error bars represent the SEM. P-values were calculated using one-sided t-test with unequal variance. * < 0.05, ** < 0.01, *** < 0.001. The data is the same in OKMS and OKS for NPCs, FBSL, iPSCs and ESCs samples because those samples were in common for OKMS and OKS (grey circles).

**c)** Proportions of each *Fyn* transcript categories in NPCs (n=8), FBSL (n=3), during early (n=2), middle (n=4) or late (n=4 for OKMS and n=2 for OKS) phases of reprogramming, or in the iPSCs (n=5) and ESCs (n=9) of the dataset. P-values were calculated using one-sided t-test with unequal variance. * < 0.05, ** < 0.01. The data is the same in OKMS and OKS for NPCs to FBSL and ESCs to iPSCs comparisons because those samples were in common for OKMS and OKS.

### Supplementary Figure 9

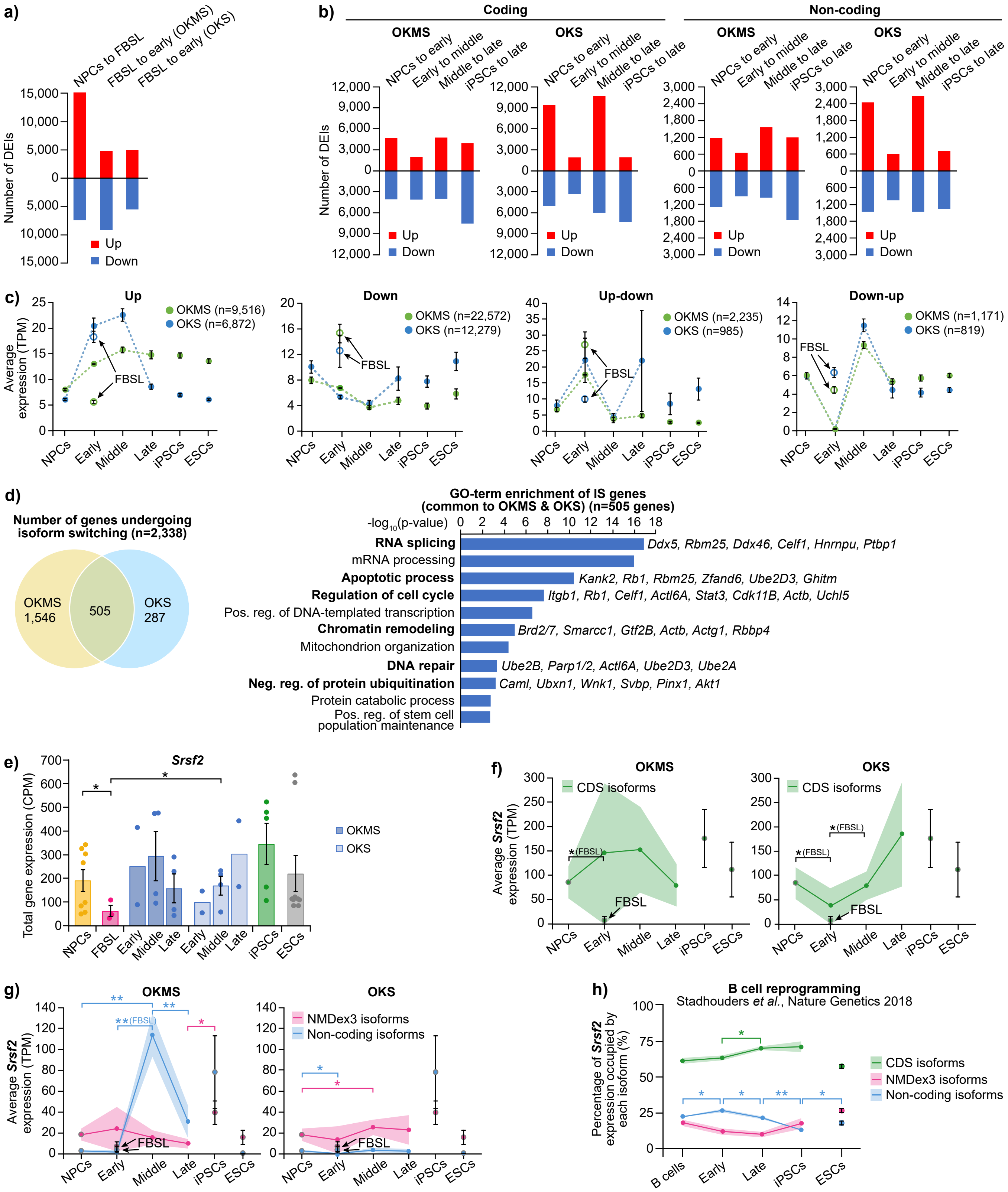

**Supplementary Figure 9: *Srsf2* isoforms switching**

**a)** Number of up- or down-regulated isoforms between different phases of OKMS and OKS reprogramming among the total isoforms (n=195,889) detected in the dataset. Significance based on log2(fold change) ≥ 2 for up-regulated isoforms or ≤ 0.5 for down-regulated isoforms with adjusted p-values ≤ 0.05 (calculated by Benjamini-Hochberg method).

**b)** Number of coding and non-coding isoforms that are significantly up- or down-regulated between the different phases of OKMS and OKS reprogramming. Significance based on log2(fold change) ≥ 2 for up-regulated isoforms or ≤ 0.5 for down-regulated isoforms with adjusted p-values ≤ 0.05 (calculated by Benjamini-Hochberg method).

**c)** Average expression of isoforms differentially expressed between D0 (n=8) and the middle phase of OKMS (n=4) or OKS (n=4) reprogramming. Error bars represent the SEM.

**d)** Overlap between the number of genes which undergo isoform switch between D0 and the middle phase of our reprogramming systems, and the corresponding GO-term enrichment analysis (biological processes) for the genes common to OKMS and OKS reprogramming. FDR ≤ 0.1. P-values were calculated by one-sided Fisher's exact test. Neg. reg.: Negative regulation.

**e)** Mean expression of *Srsf2* gene in NPCs (n=8), FBSL (n=3), during early (n=2), middle (n=4) or late (n=4 for OKMS and n=2 for OKS) phases of reprogramming, or in the iPSCs (n=5) and ESCs (n=9) of the dataset. Error bars represent the SEM and each dot represents replicates. P-values were calculated using one-sided t-test with unequal variance. * < 0.05.

**f)** and **g)** Expression of *Srsf2* isoforms in NPCs (n=8), FBSL (n=3), during early (n=2), middle (n=4) or late (n=4 for OKMS and n=2 for OKS) phases of reprogramming, or in the iPSCs (n=5) and ESCs (n=9) of the dataset. Dots and line represent average and shaded area represent the SEM for reprogramming samples. For the NPCs, FBSL, iPSCs and ESCs samples, error bars represent the SEM. P-values were calculated using one-sided t-test with unequal variance. * < 0.05, ** < 0.01. The data is the same in OKMS and OKS for NPCs, FBSL, iPSCs and ESCs samples because those samples were in common for OKMS and OKS (grey circles).

**h)** Percentage of *Srsf2* expression occupied by its isoforms in a B cell reprogramming dataset[73]. Dots and lines represent the average percentage value at each timepoint and the shaded area and error bars represent the SEM. n=4 replicates per reprogramming time point and n=2 replicates for iPSC and ESC samples. P-values were calculated for each successive phases using a two-sided t-test. * < 0.05, ** < 0.01.

### Supplementary Figure 10

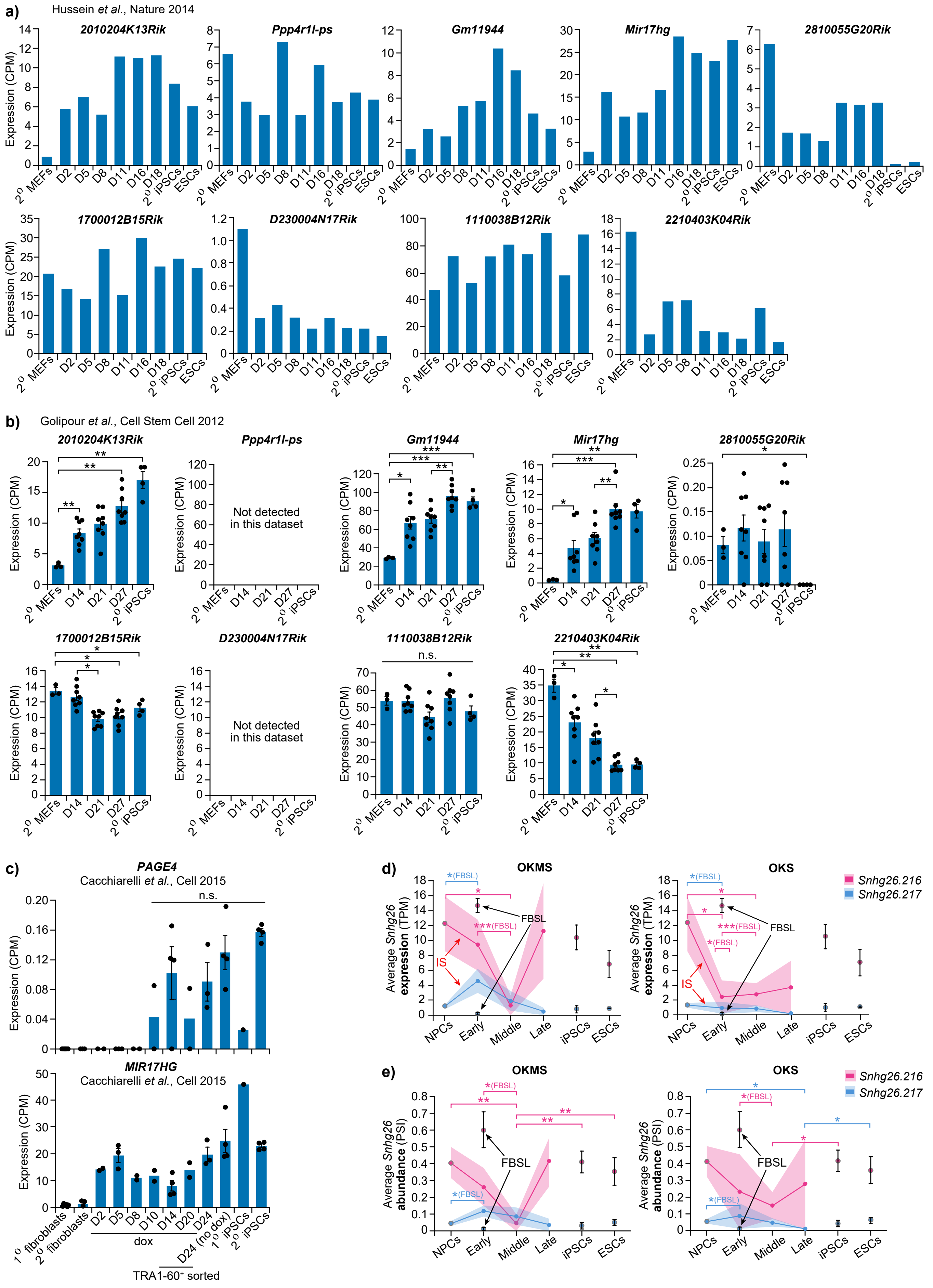

**Supplementary Figure 10: Expression of candidate non-coding transcripts during reprogramming**

**a)** and **b)** Average expression of the candidate lncRNAs other than *2700038G22Rik* at different timepoints of two 2º MEFs reprogramming RNAseq datasets[16,42] (Golipour *et al.*, Cell Stem Cell 2012: 2º MEFs (n=3), D14 (n=8), D21 (n=8), D27 (n=8), 2º iPSCs (n=4)). Error bars represent the SEM and each dot represents replicates. P-values were calculated using two-sided t-test with unequal variance. .n.s. ≥ 0.05, * < 0.05, ** < 0.01, *** < 0.001.

**c)** Average expression of *PAGE4* and *MIR17HG* lncRNAs at different timepoints of a 2º human fibroblasts reprogramming RNAseq dataset[81] (1º fibroblasts n=6, 2º fibroblasts n=3, D2 n=2, D5 n=3, D8 n=2, D10 n=2, D14 n=4, D20 n=2, D24 n=3, D24 no dox n=4, 1º iPSCs n=1, 2º iPSCs n=4). Error bars represent the SEM and each dot represents replicates. P-values were calculated using two-sided t-test with unequal variance. n.s. ≥ 0.05.

**d)** Expression of two *Snhg26* lncRNA isoforms in OKMS and OKS reprogramming phases. Lines represent average expression and shaded areas represent SEM. P-values were calculated using t-test with unequal variance. * < 0.05, *** < 0.001. The data is the same in OKMS and OKS for NPCs, FBSL, iPSCs and ESCs samples because those samples were in common for OKMS and OKS (grey circles).

**e)** PSI values of *Snhg26* transcripts abundance in the same samples of OKMS or OKS reprogramming. Lines represent the average PSI value at each timepoint and the shaded area represents the SEM. P-values were calculated using t-test with unequal variance. * < 0.05, ** < 0.01. The data is the same in OKMS and OKS for NPCs, FBSL, iPSCs and ESCs samples because those samples were in common for OKMS and OKS (grey circles).

### Supplementary Figure 11

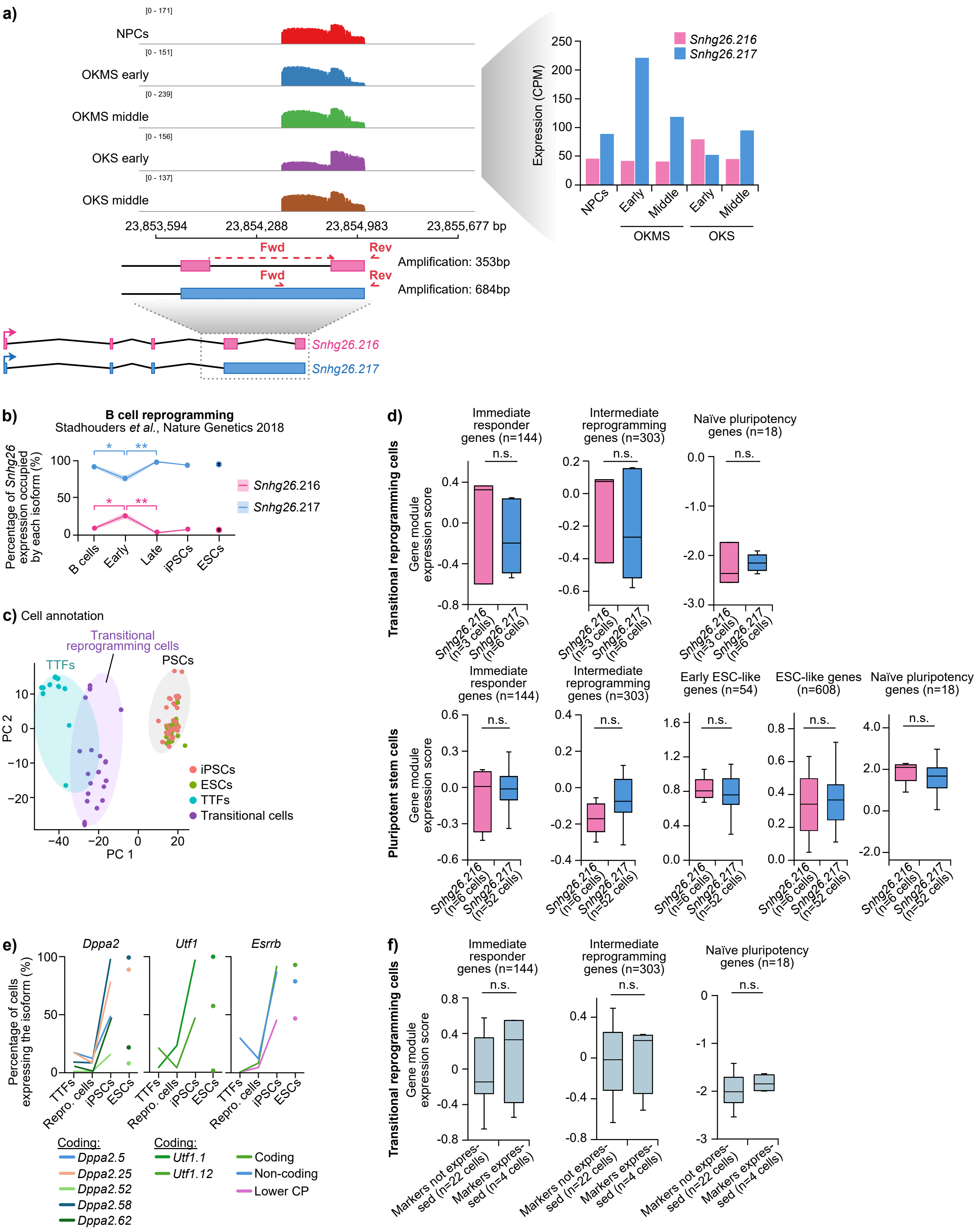

**Supplementary Figure 11: Validation and prediction of *Snhg26* isoforms' effect of the reprogramming outcome**

**a)** Histograms representing the signal from long-read sequencing of the *Snhg26* isoforms validation PCR at the genomic loci of *Snhg26*. Tracks represent NPCs (n=4) and samples from the early (n=1) and middle phases of OKMS (n=2) or OKS (n=1) reprogramming. Localization of the primers specific to *Snhg26* isoforms and size of the corresponding amplicons are represented below. Quantifications of isoforms expression is represented in on the right. Fwd: Forward; Rev: Reverse.

**b)** Percentage of *Snhg26* expression occupied by its isoforms in a B cell reprogramming dataset[73]. Dots and lines represent the average percentage value at each timepoint and the shaded area and error bars represent the SEM. n=4 replicates per reprogramming time point and n=2 replicates for iPSC and ESC samples. P-values were calculated for each successive phases using a two-sided t-test. * < 0.05, ** < 0.01.

**c)** Principal component analysis (PCA) for single-cell RNAseq[18] of reprogramming from TTFs to iPSCs. PC: Principal Component.

**d)** Seurat gene module normalized expression scores for different gene sets from Hussein *et al.*[16] between cells expressing only isoform *Snhg26.216* or *Snhg26.217,* for reprogramming transitional cells or pluripotent stem cells. Bounds of the box represent 25$^{th}$ and 75$^{th}$ quartiles, bounds of the whiskers represent minimum and maximum values, and center line represents median. P-values were calculated by Mann-Whitney non-parametric U-test. n.s. ≥ 0.05.

**e)** Percentage of cells expressing isoforms of markers of proper reprogramming (*Dppa2, Utf1,* and *Esrrb*) in a TTFs reprogramming dataset[18]. CP: Coding Potential.

**f)** Seurat gene module normalized expression scores for different gene sets from Hussein *et al.*[16] between cells expressing coding isoforms from at least two of three markers of proper reprogramming (*Dppa2, Utf1,* and *Esrrb*). Bounds of the box represent 25$^{th}$ and 75$^{th}$ quartiles, bounds of the whiskers represent minimum and maximum values, and center line represents median. P-values were calculated by Mann-Whitney non-parametric U-test. n.s. ≥ 0.05.

### Supplementary Figure 12

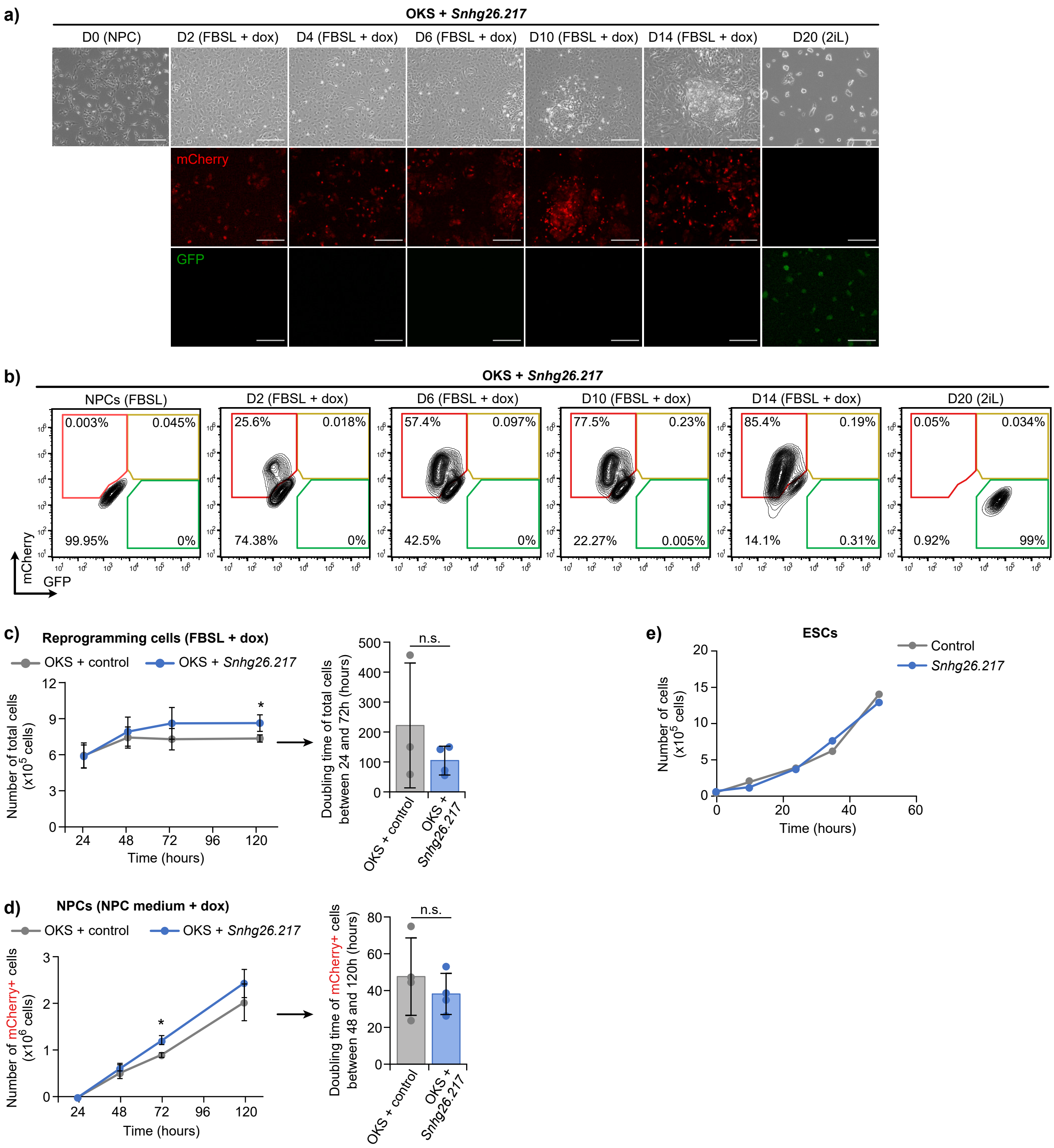

**Supplementary Figure 12: *Snhg26.217* enhances reprogramming**

**a)** Representative images in brightfield or fluorescence microscopy of the NPCs (D0), reprogramming cells at different timepoints, and final iPSCs (D20 2i-LIF) in the OKS + *Snhg26.217* reprogramming system. Scalebars = 50 µm.

**b)** Representative flow cytometry data allowing quantification of the percentages of reprogramming cells (mCherry+) and reprogrammed iPSCs (GFP+) throughout the reprogramming with OKS + *Snhg26.217*.

**c)** Average number of total cells from the OKS + control (n=3 biological replicates) and OKS + *Snhg26.217* (n=4 biological replicates) reprogramming cell lines in FBSL + doxycycline medium through the first 5 days of reprogramming. Error bars represent the SEM. P-values were calculated using one-sided t-test on paired samples. * = 0.045. Bar graph of mean cell doubling times between 24 and 72 hours of doxycycline induction. Error bars represent the STD and dot represents replicates. P-values were calculated using two-sided t-test with unequal variance. n.s. ≥ 0.05.

**d)** Average number of mCherry+ cells from the OKS + control (n=3 biological replicates) and OKS + *Snhg26.217* (n=4 biological replicates) reprogramming cell lines in NPC + doxycycline medium through the first 5 days of reprogramming. Error bars represent the SEM. P-values were calculated using two-sided t-test with unequal variance. * = 0.049. Bar graph of the mean cell doubling times between 48 and 120 hours of doxycycline induction. Error bars represent the STD and dot represents replicates. P-values were calculated using two-sided t-test with unequal variance. n.s. ≥ 0.05.

**e)** Number of cells from the ESC lines for the control (n=1) and *Snhg26.217* lncRNA (n=1) overexpression conditions in 2i-LIF + doxycycline medium through 2 days.

### Supplementary Figure 13

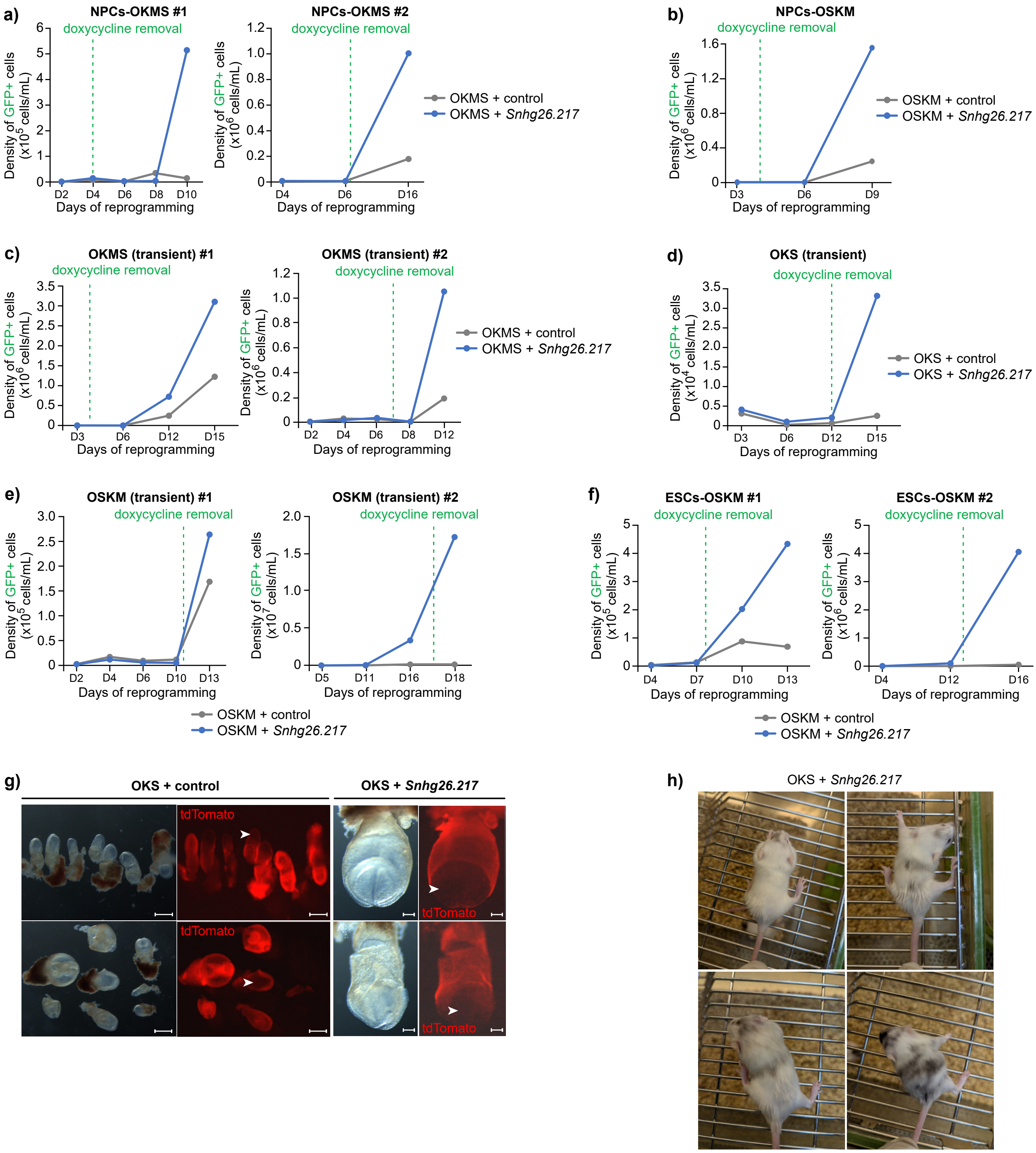

**Supplementary Figure 13: *Snhg26.217* increases the number of iPSCs independently from the reprogramming system**

**a)** Density of GFP+ cells in FBSL + doxycycline medium, or in 2i-LIF medium after doxycycline removal, during reprogramming with control or *Snhg26.217* transgenes using stable NPC lines with an OKMS (n=2) or **b)** OSKM (n=1) system.

**c)** Density of GFP+ cells in FBSL + doxycycline medium, or in 2i-LIF medium after doxycycline removal, during reprogramming with control or *Snhg26.217* transgenes using transient transfections with an OKMS (n=2), **d)** OKS (n=1), or **e)** OSKM (n=2) system.

**f)** Density of GFP+ cells in FBSL + doxycycline medium, or in 2i-LIF medium after doxycycline removal, during reprogramming with control or *Snhg26.217* transgenes using ESC line-derived NPCs with an OSKM (n=2) system.

**g)** Representative images in brightfield or fluorescence microscopy of E8-8.5 iPSC-embryo chimeras obtained from aggregation of OKS + control or OKS + *Snhg26.217* iPSC clones with 8-cell tdTomato-positive embryos for pluripotency assessment. Arrow points show examples of contribution from iPSCs. Scalebars = 200 µm (OKS + control); 100 µm (OKS + *Snhg26.217*).

**h)** Images of chimeric mice obtained from aggregation of OKS + *Snhg26.217* iPS clones with 8-cell CD-1 albino embryos, with different levels of contribution.

### Supplementary Figure 14

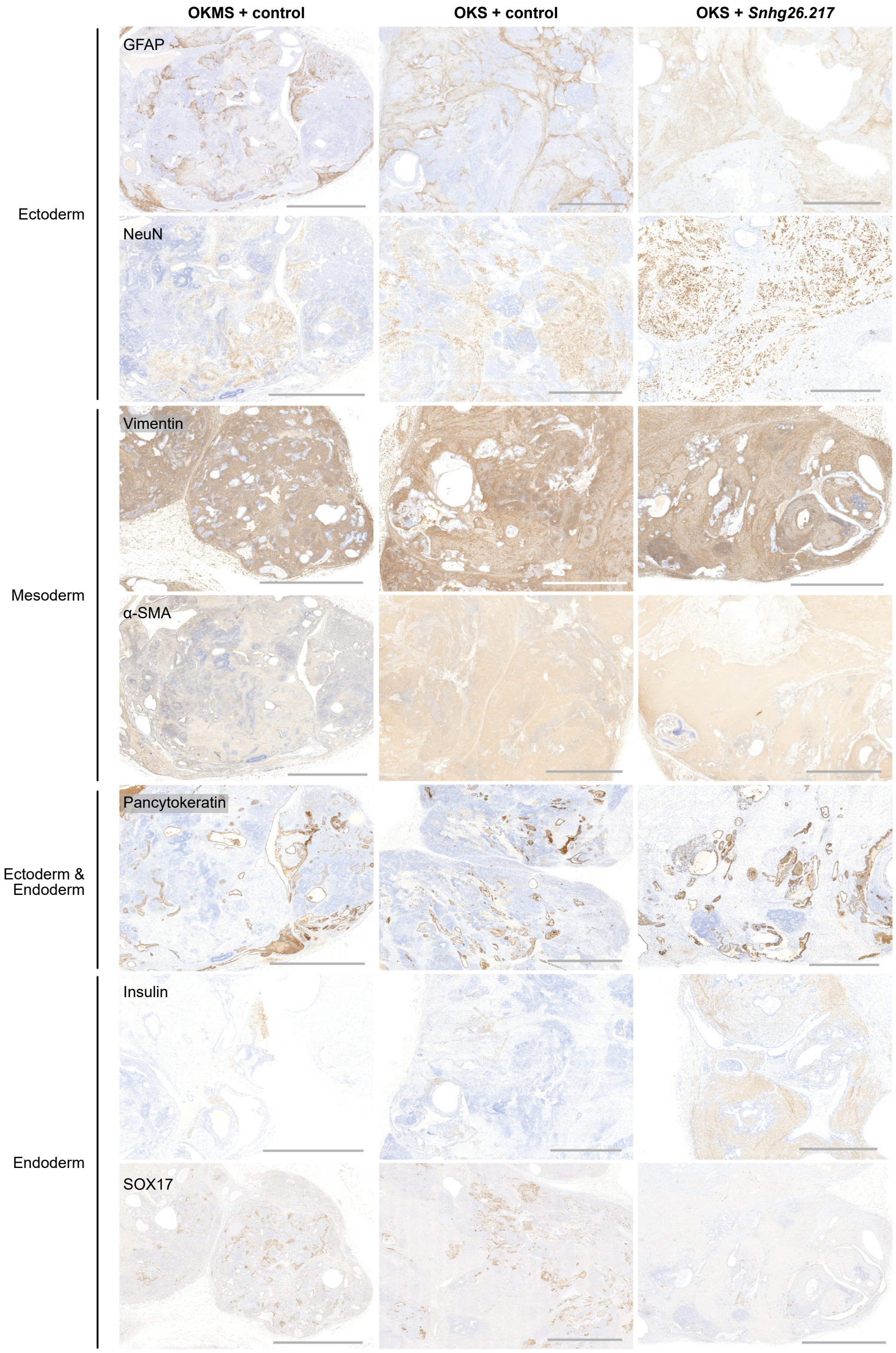

**Supplementary Figure 14: iPSC clones generate teratomas**

Representative images in brightfield of 1-month teratomas obtained from injection of OKMS/OKS + control or OKS + *Snhg26.217* iPSC clones in mice and stained with germ layer differentiation markers for pluripotency assessment. Scalebars = 2,000 μm.

### Supplementary Figure 15

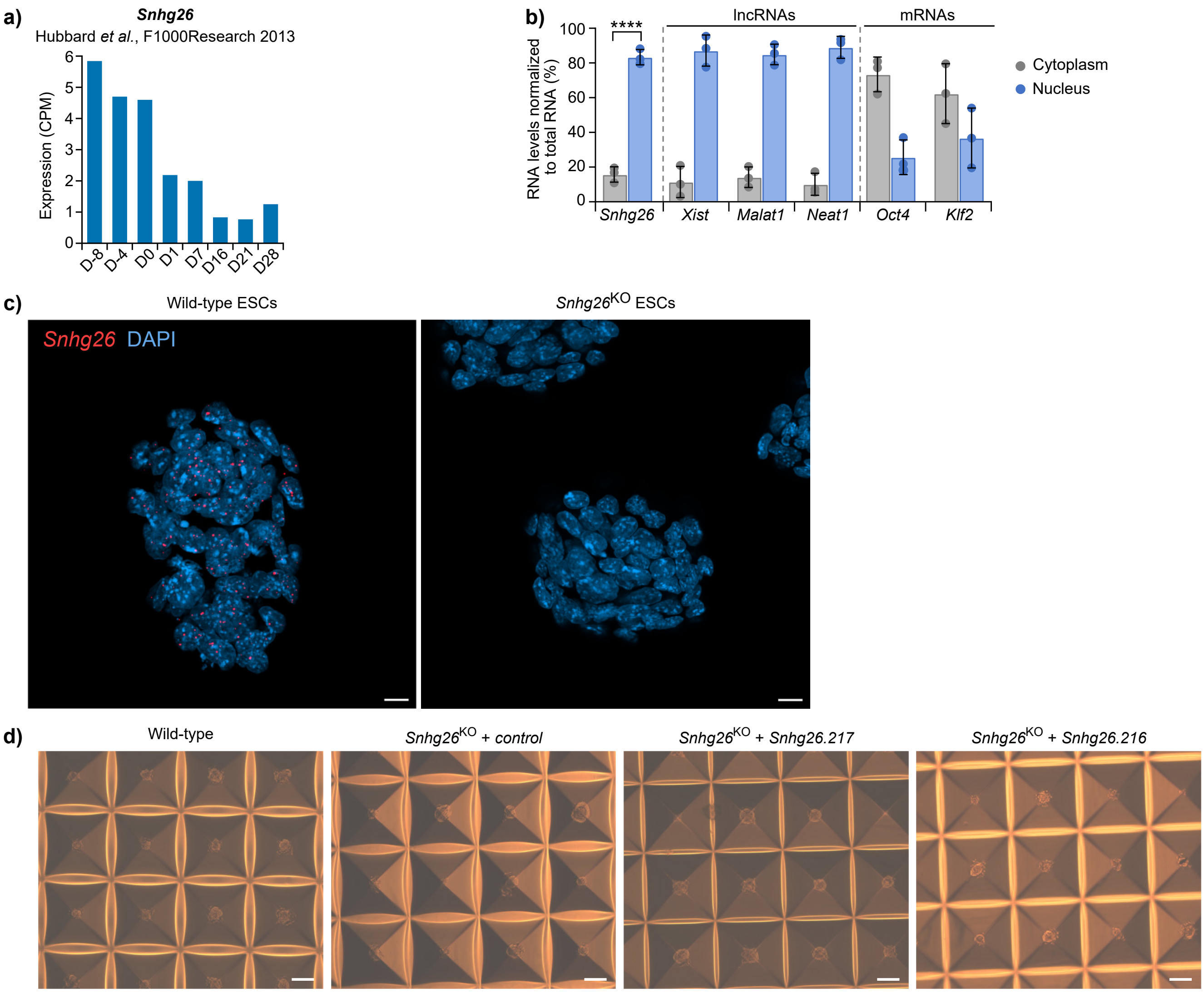

**Supplementary Figure 15: Characterization of *Snhg26* lncRNA**

**a)** Expression of *Snhg26* lncRNA in an RNAseq dataset[88] of *in vitro* differentiation of mouse ESCs into cortical neurons.

**b)** Mean percentage of *Snhg26* lncRNA and other control mRNAs and lncRNAs in the nuclear or cytoplasmic compartments of mouse ESCs after fractionation (n=3), normalized to total RNA. Dots represent the replicates and error bars represent the STD. P-values were calculated using two-sided t-test with unequal variance. **** = $4.79 \times 10^{-5}$.

**c)** Representative images of *Snhg26* lncRNA localization within mouse wild-type or *Snhg26*$^{KO}$ ESCs in 2i-LIF medium, stained with DAPI and *Snhg26*-specific probes coupled to TSA Vivid dye 650. Scalebars = 10 µm.

**d)** Representative images in brightfield microscopy of the wild-type, *Snhg26*$^{KO}$ + control, *Snhg26*$^{KO}$ + *Snhg26.217*, or *Snhg26*$^{KO}$ + *Snhg26.216* blastoids. Scalebars = 200 µm.

### Supplementary Figure 16

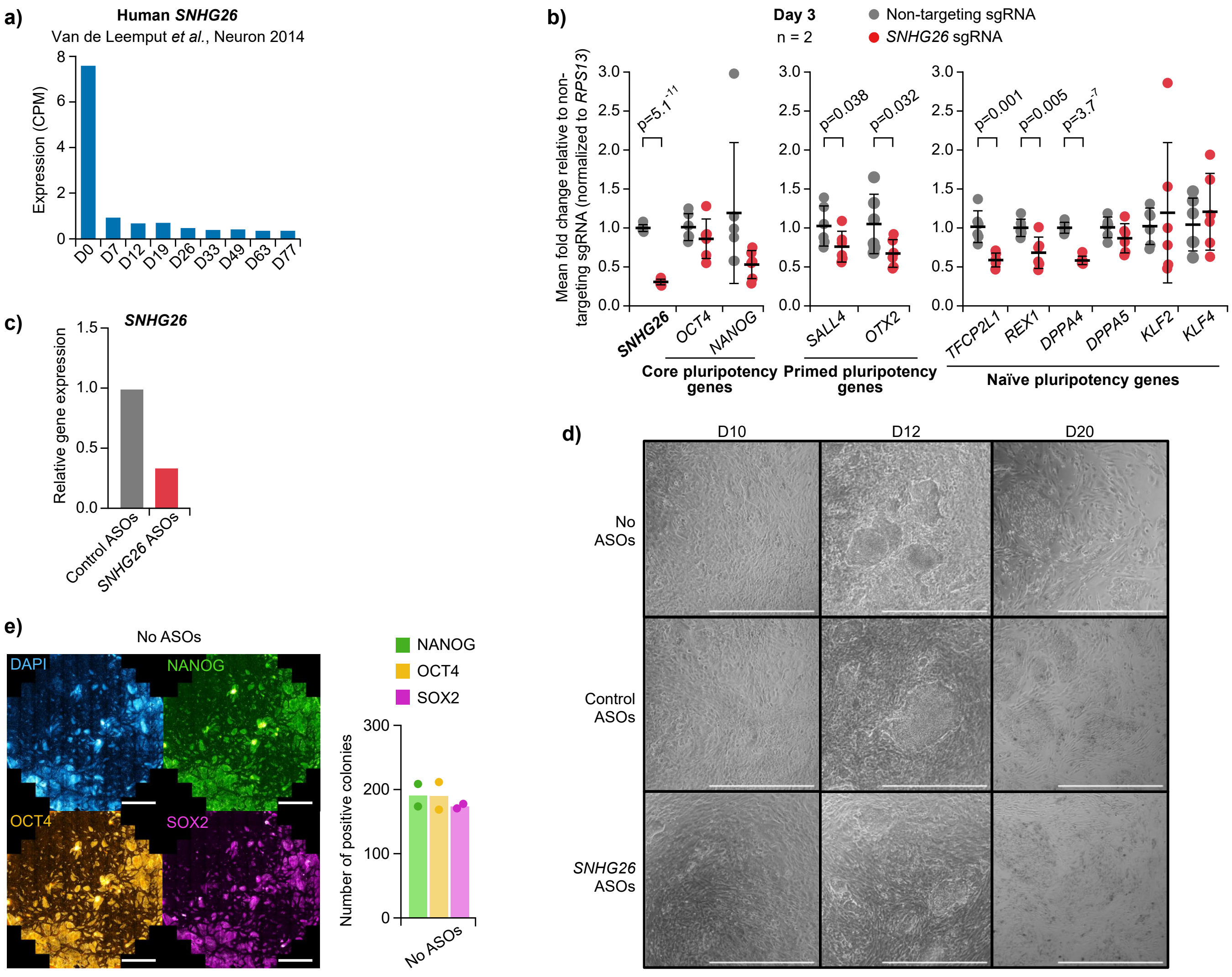

**Supplementary Figure 16: Characterization of *SNHG26* knock-down in hESCs and during reprogramming**

**a)** Expression of human *SNHG26* lncRNA in an RNAseq dataset[97] of *in vitro* differentiation of human ESCs into cortical neurons.

**b)** Mean fold changes in expression for human *SNHG26* lncRNA and pluripotency-associated genes, assessed by RT-qPCR after *SNHG26* knock-down in hESCs (n=2) using the CRISPRi system. Expression is normalized to a control sample with non-targeting sgRNA and to a housekeeping gene. Middle bars represent the average, and error bars represent the STD from 6 technical replicates. P-values were calculated using one-sided t-test with unequal variance.

**c)** Fold change in expression for *SNHG26* lncRNA 12 days after first *SNHG26*-targeting ASOs transfection in human fibroblasts. Expression is normalized to a control sample with non-targeting ASOs.

**d)** Representative brightfield images of cells without transfection, transfection of control or *SNHG26*-targeting ASOs at D10, D12, and D20 of reprogramming. Scalebars = 1,000 µm.

**e)** Representative whole-well immunofluorescence images and quantification of iPSC colonies stained with DAPI, NANOG, OCT4, and SOX2 antibodies at D20 of reprogramming without ASO transfection (n=2). Bars represent the average and dots represent biological replicates. Scalebar = 3,000 µm.

### Supplementary Figure 17

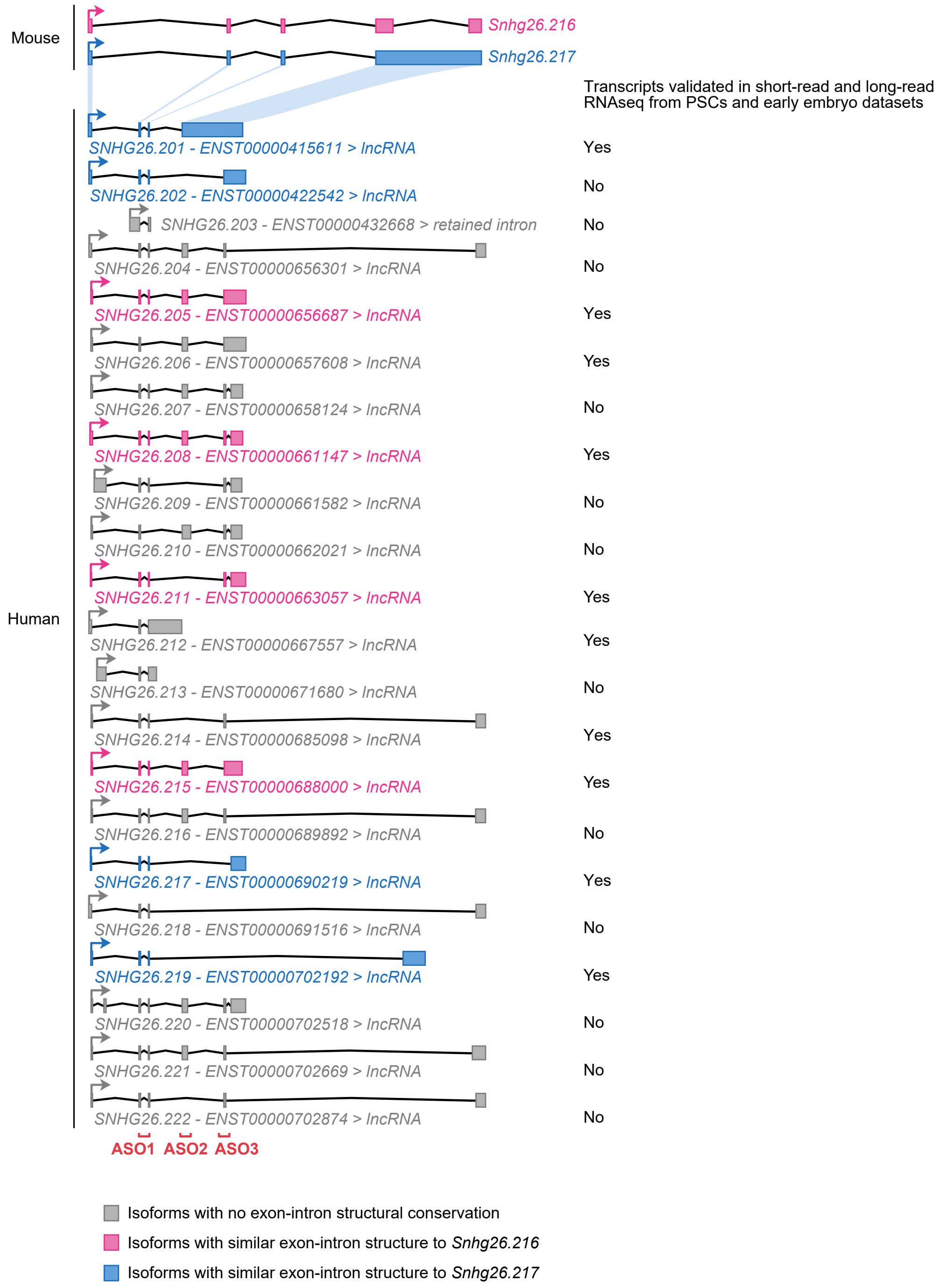

**Supplementary Figure 17: *SNHG26* isoforms**

Schematic depicting the transcripts of *SNHG26* lncRNA on the Ensembl database (release 107)[105] and whether they are expressed in short-read and long-read RNAseq datasets of human pluripotent stem cells and early embryos[93,96,97,106-110]. Transcripts with similar exon-intron structure to mouse *Snhg26.216* or *Snhg26.217* isoforms are depicted in pink or in blue, respectively. Localization of ASOs used to knock-down *SNHG26* transcripts are shown in red.

#### Supplementary Note

##### Parameters to design an efficient reprogramming system, related to Fig. 1a and Supplementary Note Fig. 1

To identify new potentially functional isoforms using long-read RNAseq, we opted to rely on the analysis of the process of reprogramming somatic cells towards induced pluripotency, as it constitutes a robust system to study the gene network ruling the pluripotent state[1,2]. However, primary and to some extent secondary reprogramming systems show efficiencies between 0.01-5%[3,4], making it almost impossible to properly define changes in isoforms with sufficiently high resolution. Moreover, most reprogramming systems use the oncogene *c-Myc* as a reprogramming factor. Although transcriptional profiles of iPSCs reprogrammed with and without *c-Myc* expression are indistinguishable, *c-Myc* promotes reprogramming by acting as a proliferation-inducing gene[5,6] and previous studies showed that *c-Myc* expression alters the quality of iPSCs generated through the reprogramming[7]. Indeed, despite triggering more reprogramming events, a bigger proportion of cells will not survive once the transgene expression is abolished[8]. On top of affecting cell proliferation and quality of the reprogramming, it also alters alternative splicing patterns of reprogramming[9]. For all these reasons, we needed to design two powerful reprogramming systems that would allow generation of sufficient amounts of reprogramming cells for analysis by lrRNAseq, and where expression of *c-Myc* could be included or omitted. Thus, patterns of alternative splicing events and differential isoform usage can be studied by comparing the two systems with and without *c-Myc*. Features that are common to the two systems thus represent alternative splicing events and isoforms that are independent from *c-Myc* proliferative-induced effects.

Many parameters affect reprogramming efficiency. First, stoichiometry of the different reprogramming factors has been shown to play an important role in the success of reprogramming, and high expression levels of both *Oct4* and *Klf4,* combined to lower levels of *Sox2* has proved to be optimal[10]. This is why we opted for doxycycline-inducible polycistronic cassettes carrying the reprogramming factors in one transcript. A main advantage of this is that the stoichiometry of the reprogramming factors is fixed, thus allowing all the transfected cells to express the same levels of reprogramming factors[11,12]. Then, a second reason why reprogramming is not very efficient is that it previously relied on lentiviruses, which were shown to be epigenetically silenced in the cells[13,14]. Even though reprogramming to pluripotency can be achieved using non-integrative systems such as adenoviruses[15,16], episomes[17], RNA[18], proteins[19] or small molecules[20,21], the efficiency of these methods remains very weak[3]. On the opposite, gene targeting allowing for transgene-based reprogramming of MEFs is up to ten-fold more efficient[22]. *PiggyBac* transposition has been shown to escape the viral silencing process and allows for random integration of large cassettes into the genome. Advantageously, *piggyBac* transposons are preferably inserted in actively transcribed regions and do not require manipulation of viruses as they can be lipotransfected[23,24]. This is why we decided to use *piggyBac* transposition, as it has proved to be more user-friendly and efficient for reprogramming[25,26].

To circumvent further the low yields of reprogramming, secondary systems such as reprogrammable mice have been generated[27-29]. Nevertheless, there are two reasons for us not using them: First, these technologies rely on the use of clonal populations that were selected based on their fitness for reprogramming. It means they are biased towards genetic or epigenetic predispositions that may hinder the decryption of the mechanisms at stake during reprogramming. Second, reprogrammable mice sometimes exhibit poor efficiency (0.2-2%) because the differentiation of the line primarily containing the reprogramming factors may select against high-expressor cells, and thus results in secondary lines able to express these factors only poorly[27].

We figured out that generating NPC stable cell lines where all cells carry doxycycline-inducible transgenes containing the reprogramming factors would allow for a higher reprogramming efficiency while conserving intrinsic cell heterogeneity. First, because NPCs are easy to grow and transfect[30,31], they are quickly expandable to reach the high numbers of cells needed for lrRNAseq protocols, specifically for early timepoint analysis. Also, they are less limited by reaching a maximum passage number like MEFs[32]. Second, unlike MEFs, NPCs already express *Sox2*, which facilitates and accelerates their reprogramming towards induced pluripotency: It has been shown that NPCs can be readily reprogrammed using only *Oct4* and *Klf4* overexpression[33,34]. In our hands, we could not reprogram NPCs with OK very efficiently (data not shown), so we opted for a combination of *Oct4*, *Klf4* and *Sox2* (OKS). Using NPCs is also useful when not including *c-Myc* in the cocktail of reprogramming factors, as timelines to reprogram MEFs with OKS extend from 14 to 21 days with a decreased efficiency[6,33]. Finally, we opted to generate stable cell lines directly from NPCs as it was significantly more efficient in reprogramming than using NPCs derived from ESCs already harboring our expression cassettes, which yielded poor reprogramming efficiency (0.3%, **Supplementary Note Fig. 1a**). As a control condition, we consistently compared the well-established *Oct4, Klf4, c-Myc, Sox2* combination of reprogramming factors to our newly generated OKS reprogramming system f o r downstream analyses.

Thus, we designed our reprogramming systems by directly introducing into mouse neural progenitor cells two *piggyBac* transposon-based expression cassettes (**Fig. 1a**): 1) A polycistronic construct to express the reprogramming factors *Oct4*, *Klf4*, *Sox2* with *Myc* (OKMS) or without *Myc* (OKS), and 2) a polycistronic *rtTA* and selection marker expression construct. These constructs are doxycycline-responsive and contain *mCherry* as a reporter. Those NPCs were differentiated from mouse Oct4-GFP ESCs[35], which harbor a *GFP* reporter to monitor acquisition of ESC-like states. Thus, upon DOX induction, reprogramming factor expression is followed by flow cytometry through mCherry and acquisition of a pluripotent state is represented by GFP expression. mCherry-expressing cells peaked at D6 (91.5% of cells) for OKMS and a week later, to lesser extent, (58.35% of cells at D14) for OKS, suggesting a very efficient activation of the transgenes and population-based reprogramming regardless of the presence or absence of *Myc* (**Supplementary Note Fig. 1b**). We monitored *Oct4-GFP* reactivation as an indication that iPSCs are reaching a stabilization phase[36-38] and found that the number of GFP-expressing cells increased at D14 for OKMS (20.71%) and at D18 for OKS (9.92%), to reach 44.44% and 21.78% by D20, respectively (**Supplementary Note Fig. 1c**). This is concomitant with spontaneous silencing of transgenes, as only 11.05% and 2.99% of D14 OKMS and OKS cells still expressed mCherry (i.e. reprogramming factor expression), respectively (**Supplementary Note Fig. 1b**). This is consistent with the fact that *bona fide* iPSCs can only be generated once the transgenes are silenced, enabling the subsequent reactivation of endogenous pluripotency genes in the later phase of reprogramming[37-39]. To determine the minimal duration of DOX induction necessary to get transgene-independent iPSCs, thus mimicking transgene silencing, we removed DOX at different timepoints of OKS reprogramming (2, 4, 6, 8, or 10 days) and analyzed for GFP expression at D14 (**Supplementary Note Fig. 1d**). We found that transgenes are needed for at least six days to achieve complete reprogramming with our OKS system, peaking at D8, and then significantly coming down with longer DOX exposure. This is consistent with our previous findings that prolonged reprograming factor expression leads to alternative routes other than iPSCs[1]. Therefore, DOX was removed at D6 for both OKMS and OKS experiments and cells were subsequently placed in 2i-LIF conditions to generate GFP+ iPSCs. DOX removal led to a rapid decrease in mCherry+ cells with most cells silencing the exogenous reprogramming factors by D14 (**Supplementary Note Fig. 1b**, right). This was followed by an increase in GFP+ cells following DOX removal, reaching 65.64% for OKMS and 63.35% of GFP+ cells for OKS by D14 (**Supplementary Note Fig. 1c**, right). We thus demonstrate that both our OKMS and OKS reprogramming systems follow population-based reprogramming and lead to the generation of iPSCs with high efficiency. Of note, OKMS dynamics were slightly faster than previously shown in secondary reprogramming systems because of differences in starting cell types (NPCs versus MEFs).

We then compared our list of differentially expressed genes (DEGs) to our previous analyses of reprogramming[1]. From this Hussein *et al.* DEG list, 72% of protein-coding genes were found to be expressed at some point in our reprogramming dataset using 2° MEFs (**Supplementary Note Fig. 1e** and **Supplementary Data 1**). This demonstrates high correlation between our systems and previous MEF-based systems. This suggests that even in the absence of *Myc*, our system efficiently induces expression profiles reminiscent of other systems that include *Myc*.

Altogether, these data demonstrate how our new design of OKMS and OKS reprogramming systems leads to very efficient models of pluripotency induction that can be used to investigate isoform diversity using lrRNAseq.

### Supplementary Note Figure 1

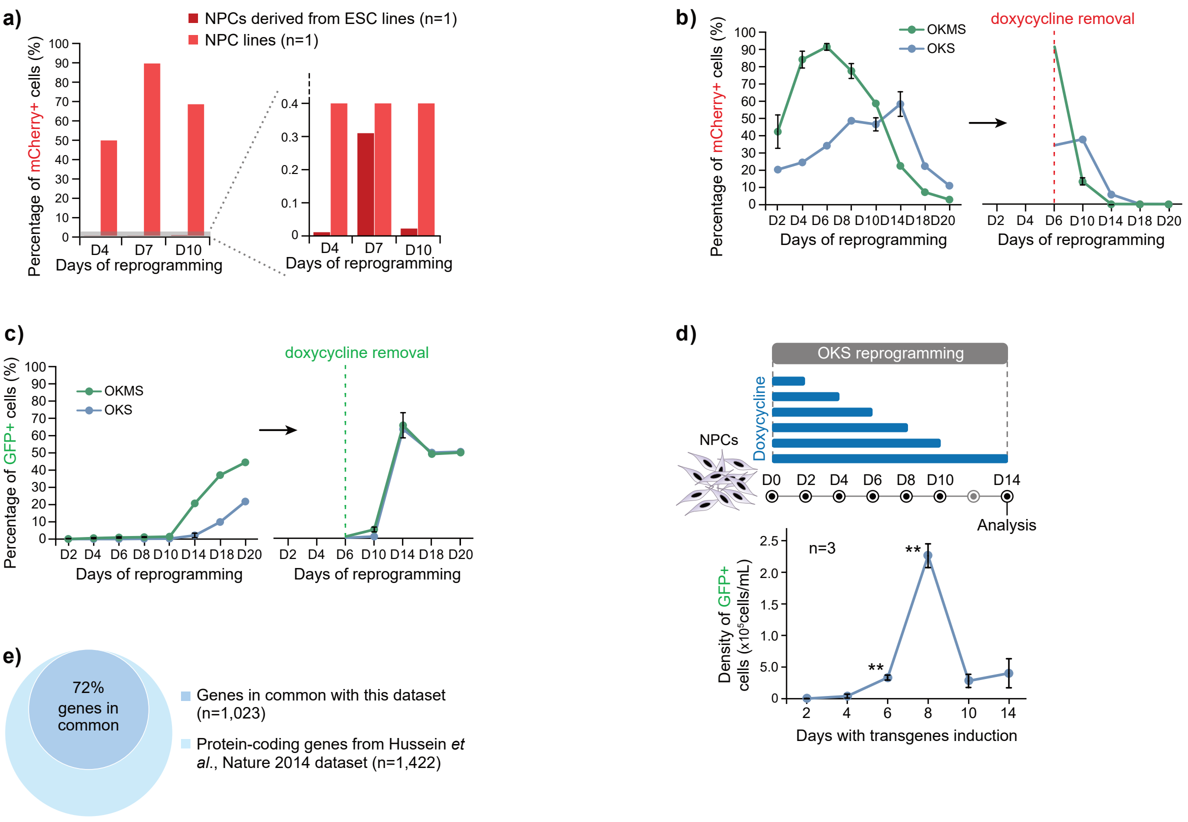


**Supplementary Note Fig. 1: Designing an efficient reprogramming system, related to Fig. 1**

**a)** Representative flow cytometry data of the percentage of mCherry+ cells (i.e. reprogramming cells), up to D10 of reprogramming, when reprogramming NPCs derived from stably transfected ESCs lines (n=1, dark red) to stably transfected NPCs lines derived from wild-type ESCs (n=1, light red).

**b)** Percentages of mCherry+ cells (i.e. expressing OKMS or OKS) or **c)** GFP+ cells (i.e. endogenous *Oct4*) in FBSL+doxycycline medium,or after doxycycline removal in 2iL medium, monitored using flow cytometry. Error bars represent the SEM (n=3-9).

**d)** Flow cytometry data of the density of reprogrammed iPSCs (GFP+) at D14 of OKS reprogramming, after removing doxycycline at different timepoints. Error bars represent the SEM (n=3). P-values were calculated using two-sided t-test with unequal variance. ** p-value = 0.01.

**e)** Overlap between the number of immediate responder, intermediate reprogramming, early ESC-like and ESC-like genes detected between our lrRNAseq dataset and the dataset from Hussein *et al*., Nature 2014.

### Supplementary Note Figure 2

#### Situation #1

**Control reprogramming**

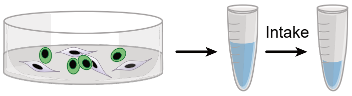


Volume intake by flow cytometer: 100µL
- Number of cells counted by flow cytometer: 100,000 cells
- Number of GFP+ cells counted by flow cytometer: **50,000 cells**

Result given by flow cytometer: **50%** GFP+ cells

**Reprogramming with *Snhg26* lncRNA**

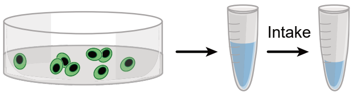


Volume intake by flow cytometer: 100µL
- Number of cells counted by flow cytometer: 100,000 cells
- Number of GFP+ cells counted by flow cytometer: **100,000 cells**

Result given by flow cytometer: **100%** GFP+ cells

↳ In this situation, the GFP+ percentages reflect well that reprogramming with *Snhg26* lncRNA yielded twice the absolute quantity of GFP+ cells compared to the control, because there was no difference in the total cell density.

#### Situation #2

**Control reprogramming**

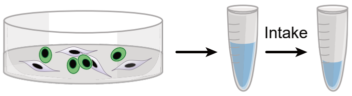


- Volume intake by flow cytometer: 100µL
- Number of cells counted by flow cytometer: 100,000 cells
- Number of GFP+ cells counted by flow cytometer: **50,000 cells**

Result given by flow cytometer: **50%** GFP+ cells

**Reprogramming with *Snhg26* lncRNA**

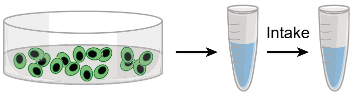


- Volume intake by flow cytometer: **50µL**
- Number of cells counted by flow cytometer: 100,000 cells
- Number of GFP+ cells counted by flow cytometer: **100,000 cells**

Result given by flow cytometer: **100%** GFP+ cells

↳ Reprogramming with *Snhg26* lncRNA yielded four times the absolute quantity of GFP+ cells compared to the control, which is is underestimated by the GFP+ percentages. Thus, because the total cell density is affected when overexpressing *Snhg26*, it is necessary to take into account the density of GFP+ cells.

#### Situation #3

**Control reprogramming**

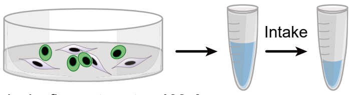


- Volume intake by flow cytometer: **100µL**
- Number of cells counted by flow cytometer: 100,000 cells
- Number of GFP+ cells counted by flow cytometer: 50,000 cells

Result given by flow cytometer: **50%** GFP+ cells

**Reprogramming with *Snhg26* lncRNA**

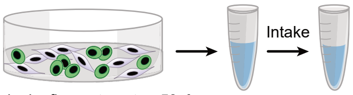


- Volume intake by flow cytometer: **50µL**
- Number of cells counted by flow cytometer: 100,000 cells
- Number of GFP+ cells counted by flow cytometer: 50,000 cells

Result given by flow cytometer: **50%** GFP+ cells

↳ Reprogramming with *Snhg26* lncRNA yielded twice the absolute quantity of GFP+ cells compared to the control, which is is underestimated by the GFP+ percentages. Thus, because the total cell density is affected when overexpressing *Snhg26*,it is necessary to take into account the density of GFP+ cells.

**Conclusion:** Overexpression of *Snhg26* lncRNA during reprogramming increases the quantity of cells committing to reprogramming (i.e. mCherry+), which leads to an increase in the total number of cells in the culture dish (Fig. 7d and Supplementary Fig. 12c) compared to a control condition. Because when analyzing the reprogramming cells with flow cytometry the same number of total events is counted, it does not take into account this difference in total density of cells (i.e. differences in volume intake) between control and *Snhg26* overexpression reprogramming. Thus, the percentage of GFP+ cells evaluated (Fig. 7b) by flow cytometry is underestimated. This is why the total density of mCherry+ and GFP+ cells (Fig. 7e-f) needs to be taken into account when assessing the yield of GFP+ cells during reprogramming.

### Supplementary Note Figure 3

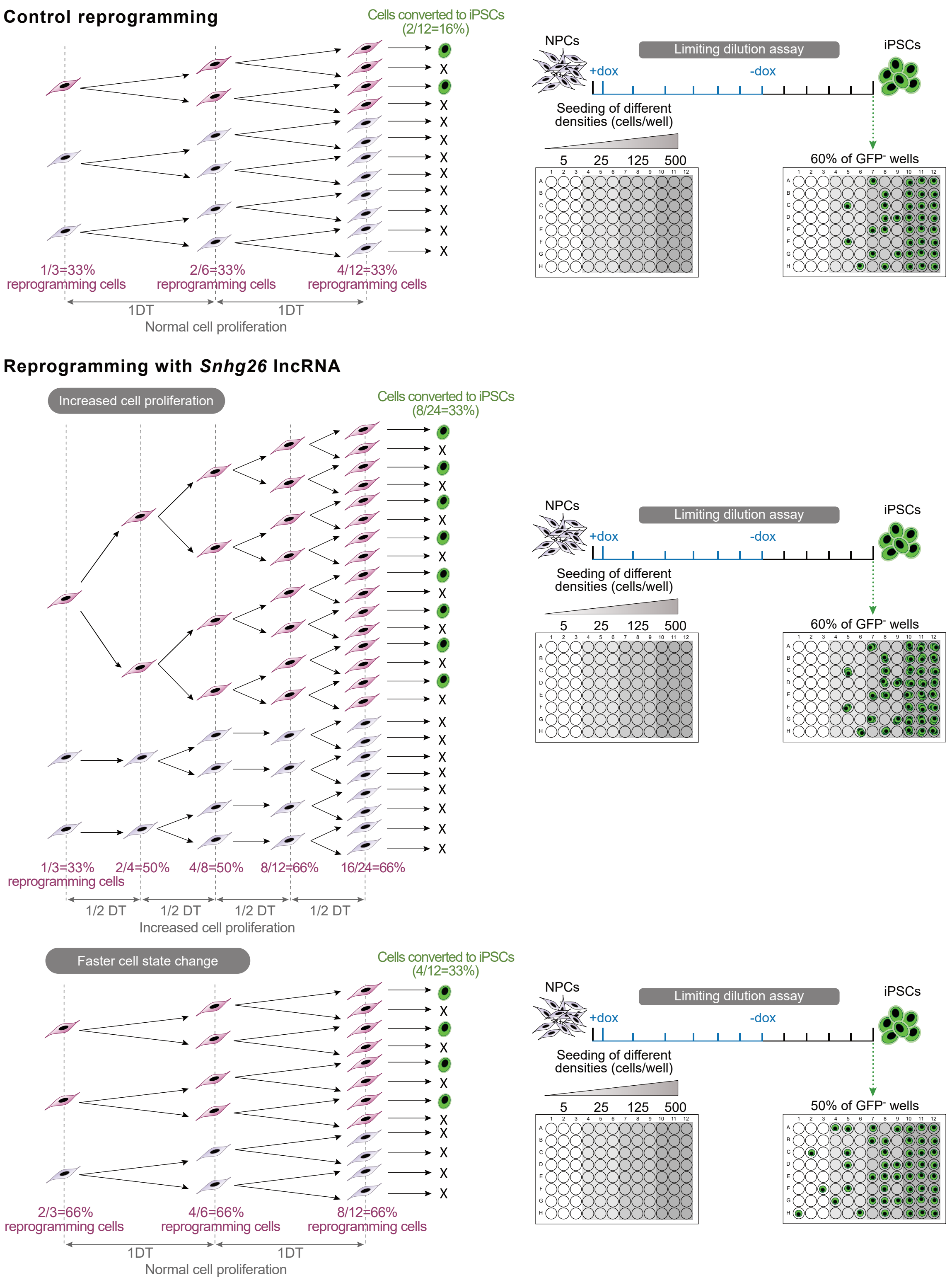


**Conclusion:** Overexpression of *Snhg26* lncRNA during reprogramming can increase the quantity of reprogramming cells (mCherry+) and iPSCs (GFP+) compared to a control condition by increasing the proliferation of reprogramming cells. This would lead to a decreased doubling time (DT) of the reprogramming cells but would not affect the percentage of wells negative for GFP+ cells in a limiting dilution assay (LDA). This is not the case when *Snhg26* is overexpressed (Fig. 7d, Fig. 7h, and Supplementary Fig. 12c).

Another possibility is that overexpression of *Snhg26* lncRNA can increase the quantity of reprogramming cells (mCherry+) and iPSCs (GFP+) compared to a control condition by increasing the number of cells able to engage in the reprogramming process (faster cell stage change). This would lead to a lower number of wells negative for GFP+ cells in an LDA, but would not affect the DT of the reprogramming cells. This is the case when *Snhg26* is expressed, as observed in Fig. 7d, Fig. 7h, and Supplementary Fig. 12c.
